# Rational development of a small-molecule activator of CK1γ2 that decreases C99 and beta-amyloid levels

**DOI:** 10.1101/2023.04.28.538773

**Authors:** Victor Hugo Bustos, Yashoda Krishna Sunkari, Anjana Sinha, Maria Pulina, Ashley Bispo, Maya Hopkins, Alison Lam, Sydney F. Kriegsman, Emily Mui, Emily Chang, Ana Jedlicki, Hannah Rosenthal, Marc Flajolet, Paul Greengard, Subhash C. Sinha

## Abstract

Alzheimer’s disease (AD) is a debilitating neurodegenerative disorder characterized by the accumulation of beta-amyloid (Aβ), C99, and Tau in vulnerable areas of the brain. Despite extensive research, current strategies to lower Aβ levels have shown limited efficacy in slowing the cognitive decline associated with AD. Recent findings suggest that C99 may also play a crucial role in the pathogenesis of AD.

Our laboratory has discovered that CK1γ2 phosphorylates Presenilin 1 at the γ-secretase complex, leading to decreased C99 and Aβ levels. Thus, CK1γ2 activation appears as a promising therapeutic target to lower both C99 and Aβ levels.

In this study, we demonstrate that CK1γ2 is inhibited by intramolecular autophosphorylation and describe a high-throughput screen designed to identify inhibitors of CK1γ2 autophosphorylation. We hypothesize that these inhibitors could lead to CK1γ2 activation and increased PS1-Ser367 phosphorylation, ultimately reducing C99 and Aβ levels.

Using cultured cells, we investigated the impact of these compounds on C99 and Aβ concentrations and confirmed that CK1γ2 activation effectively reduces their levels. Our results provide proof of concept that CK1γ2 is an attractive therapeutic target for AD.

Future studies should focus on the identification of specific compounds that can inhibit CK1γ2 autophosphorylation and evaluate their efficacy in preclinical models of AD. These studies will pave the way for the development of novel therapeutics for the treatment of AD.

## Introduction

Protein kinases regulate a wide array of cellular functions through protein phosphorylation. Since elevated kinase activity is involved in many disease states, especially in the onset and development of cancer, protein kinase inhibition has been an active area of research. Over the last two decades, more than 50 small-molecule kinase inhibitors have been approved as therapeutic drugs^1^. In contrast, just a few small-molecule protein kinase activators have been reported, although such activators may also have therapeutic applications in a growing number of diseases.

Several reports indicate that the protein kinase CK1 is linked to Alzheimer’s disease (AD). In mammals, CK1 is a family of Ser/Thr protein kinases, comprising CK1α, CK1δ, CK1ε, CK1γ1, CK1γ2 and CK1γ3 members^2^. It has been described that mRNA^3^ and protein levels of both CK1δ and CK1ε are elevated in Alzheimer’s disease. CK1α associates with neurofibrillary tangles^4^ and CK1δ accumulates in granulovacuolar bodies^5^. In addition, CK1 isoforms have been shown to phosphorylate Tau and promote pathogenesis^6,7^. CK1 isoforms have also been shown to regulate beta-amyloid levels. Overexpression of constitutively active CK1ε, leads to an increase in Aβ peptide production. Conversely, three structurally dissimilar CK1-specific inhibitors significantly reduced endogenous Aβ peptide production^8^. Recently, we showed that CK1γ, through the phosphorylation of Presenilin 1, was able to induce the degradation of C99, decreasing Aβ levels^9,10^.

Alzheimer’s disease is characterized by the accumulation of beta-amyloid, C99 and Tau in the brain. The Aβ peptide is generated through sequential proteolysis of the Amyloid Precursor Protein (APP), first by the action of beta-secretase, generating C99, and then by the Presenilin 1 (PS1) enzyme in the gamma-secretase complex. We have recently found that the protein kinase CK1γ2 phosphorylates Presenilin 1 at Ser367, leading to decreased C99 and Aβ levels^9,10^.

One potential drug discovery strategy to decrease C99 and Aβ levels involves activating CK1γ2 to increase PS1-Ser367 phosphorylation. In this study, we demonstrate that CK1γ2, similar to other CK1 family members^10-14^, is inhibited by autophosphorylation. Previous research on protein kinase CK1α indicates that this kinase employs distinct structural domains to recognize different consensus phosphorylation sites^15^. Mutational experiments suggest that various regions of the enzyme are involved in the recognition of substrate sequences. In theory, it may be possible to selectively inhibit autophosphorylation without affecting the phosphorylation of external substrates, thus preserving the enzyme’s ability to phosphorylate other targets. We designed a high-throughput screen to identify inhibitors of autophosphorylation, hypothesizing that this could enhance CK1γ2 activity towards selected substrates. In this report, we present the rational design of a kinase activator through the identification of small-molecule inhibitors of CK1γ2 autophosphorylation that increase the enzyme’s activity. We also discuss the medicinal chemistry efforts undertaken to improve the potency of these compounds and demonstrate the effects of these compounds on CK1γ2 activation and Aβ levels in cultured cells.

## Results

### CK1γ2 autophosphorylation reduces its kinase activity

Several members of the CK1 family have been shown to autophosphorylate, however autophosphorylation of mammalian CK1γ2 has not yet been investigated. To examine the ability of CK1γ2 to autophosphorylate we developed an in vitro autophosphorylation assay, using a recombinant CK1γ2 protein with a GST tag at the N-terminal. GST-CK1γ2 was purified using affinity purification with GSH-Sepharose, followed by cleavage of GST with AcTEV protease. The autophosphorylation as a function of time for a previously dephosphorylated CK1γ2 is shown in **Figure 1A**. Maximum autophosphorylation under these conditions is achieved after 30 min of incubation with ^32^P-γ-ATP.

**Figure 1.**
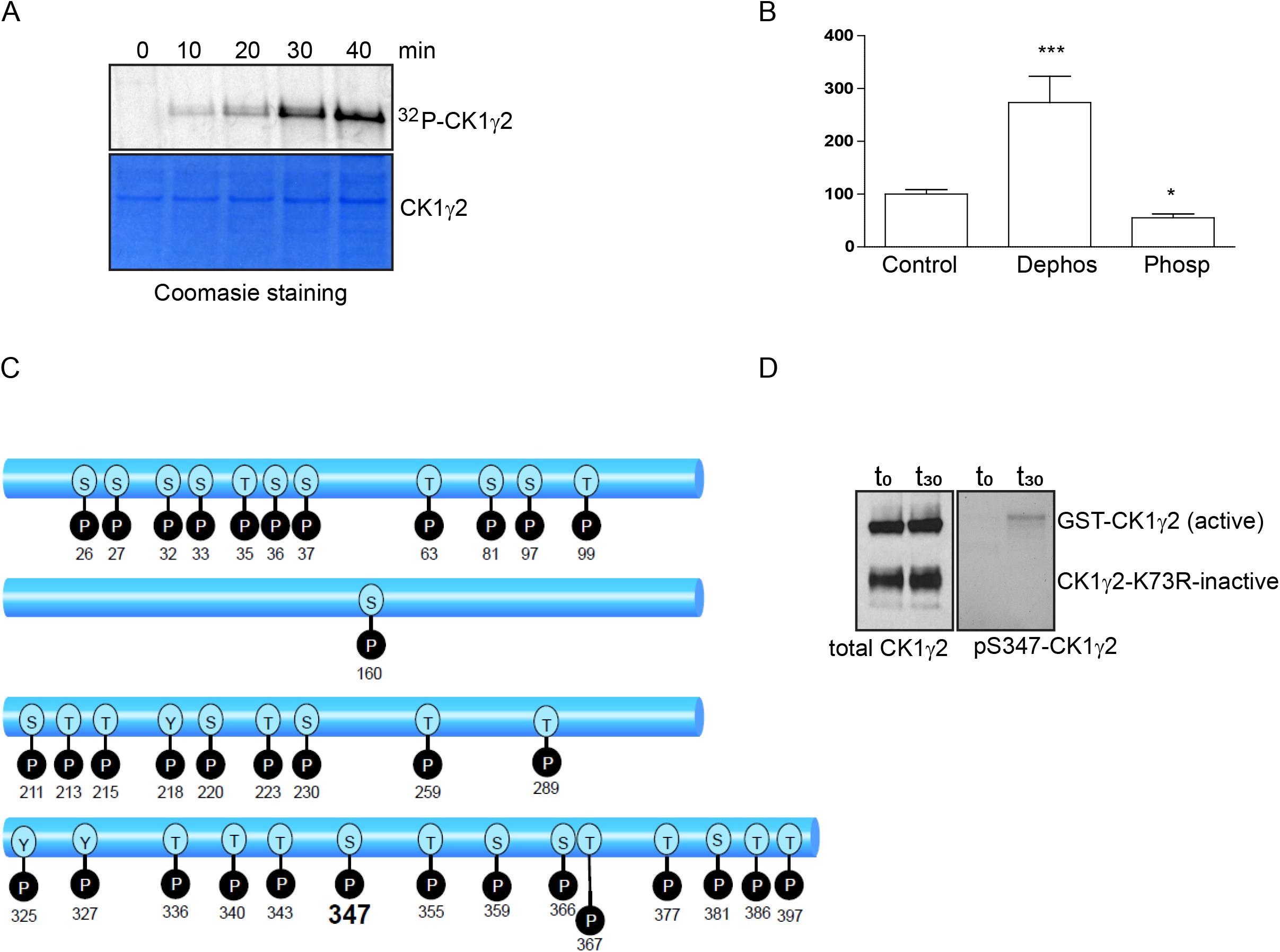
A) Autoradiograph (upper) and Coomasie staining (lower) of autophosphorylated purified recombinant CK1γ2. B) Casein phosphorylation of phosphorylated and dephosphorylated recombinant CK1γ2. Data for at least three experiments (mean ± SEM) were compared with the control condition. ⋅, P < 0.05; ⋅⋅⋅, P < 0.001; two-tailed Student’s t test, 95% significance level. C) Diagram of autophosphorylation sites in CK1γ2 as found by MS/MS D) Western blot of GST-CK1γ2 or CK1γ2-K73R mutant incubated with total CK1γ2 or pS347-CK1γ2 antibodies.

**Figure 2.**
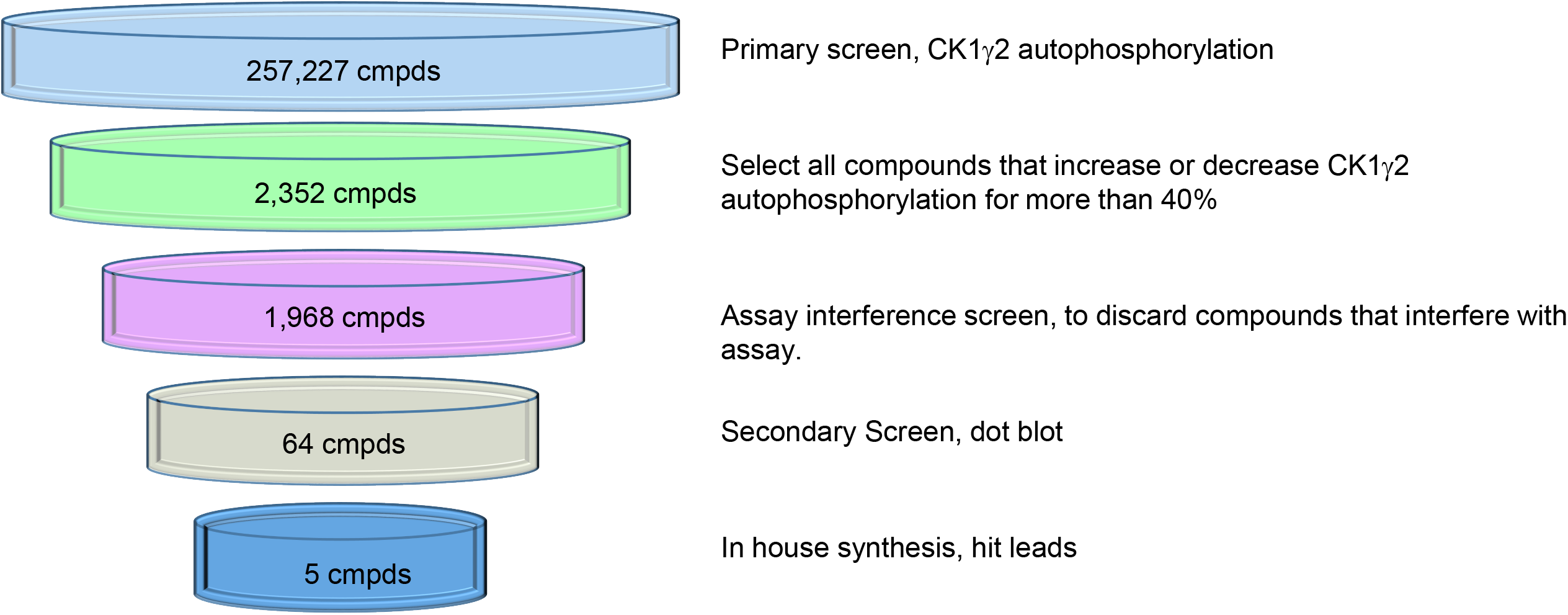
HTS assay funnel

To evaluate the effect of autophosphorylation on CK1γ2 activity, casein phosphorylation was measured. As shown in **Figure 1B**, pre-incubation of recombinant CK1γ2 with lambda phosphatase increased its activity and incubation with ATP decreased its activity. Our results show that the activity of CK1γ2, like other CK1 isoforms, is inhibited when autophosphorylated (**Figure 1B**). To determine the sites on which CK1γ2 autophosphorylates, we incubated recombinant CK1γ2 with non-radioactive ATP under autophosphorylation conditions and analyzed its autophosphorylation sites by tandem mass spectrometry (MS/MS). MS/MS analysis identified thirty-five autophosphorylation sites, distributed in clusters in the N-terminal, medial and C-terminal region of CK1γ2 (**Figure 1C**). The strongest signal was ascribed to Ser347, which accounts for 60% of all phosphate incorporation. To test whether CK1γ2 autophosphorylation at Ser347 is due to an intramolecular or intermolecular mechanism, we used WT or the inactive kinase CK1γ2-K75R as “substrates”, and GST-tagged recombinant WT-CK1γ2 as the “kinase”. We developed a phosphospecific antibody that recognizes CK1γ2 only when phosphorylated at Ser347. As shown in **Figure 1D**, autophosphorylation of CK1γ2 at Ser347 was detected only in WT-CK1γ2, while WT-CK1γ2 did not phosphorylate the GST-CK1γ2 K75R mutant. These results suggested that autophosphorylation of CK1γ2 occurs via an intramolecular mechanism. We hypothesized that a small molecule able to inhibit an intramolecular phosphorylation reaction but not an intermolecular phosphorylation would be able to activate this kinase towards an exogenous substrate. Thus, we developed and optimized a high throughput screen (HTS) to find inhibitors of CK1γ2 autophosphorylation.

### Inhibitors of CK1γ2 autophosphorylation identified by HTS

In a first step, we adapted the ADP-glo assay to measure ATP consumption in the autophosphorylation reaction. This assay is based on the breakdown of ATP to ADP plus phosphate. An enzyme consumes the remaining ATP, then ADP is converted back to ATP. Then, ATP fuels the luciferase reaction, producing light, which can be measured in a spectrophotometer. At the beginning of the screen, we performed a LOPAC validation, which mimics an HTS campaign but uses the LOPAC^1280^ compound library as a test library. It is used to calculate plate Z-factor, and signal to background. Our optimized assay had an average Z’ of 0.86. We screened 257,227 molecules at the Drug Discovery Center at the Rockefeller University. After a primary screen against CK1γ2 autophosphorylation, we selected 2352 compounds that either increased or decreased CK1γ2 autophosphorylation for more than 40%. The dose response of these compounds was determined from using ten consecutive half dilutions points ranging from 32 uM to 0.3 nM to identify 64 primary hit compounds that affected CK1γ2 autophosphorylation in a dose-dependent manner.

Next, we performed a secondary screen to investigate whether these primary hit compounds were able to modulate the phosphorylation of recombinant Presenilin 1. For this, we designed a dot blot assay, using recombinant PS1 3^rd^ loop purified from E. Coli inclusion bodies. This protein was incubated with CK1γ2 in the presence of ATP and the primary hit compounds. Next, PS1 Ser367 phosphorylation was analyzed by a specific phosphoantibody^11^. We examined all primary hit compounds using this assay to identify 55 compounds that were able to increase PS1 phosphorylation at least by 200% at 16 uM. All 55 compounds, labeled as CKR1-55, were further examined by performing Western Blots using PS1 3^rd^ loop and compounds at multiple concentrations (between 0-128 μM). Five compounds, CKR5, CKR20, CKR25, CKR31, and CKR49 (**Figure 3**), consistently increased PS1 phosphorylation at PS1-Ser367, in a dose dependent manner with IC_50_ <32 μM. We re-confirmed the activity of all 5 hit compounds using samples that were prepared, in house, to find that CKR5 and CKR49 are more potent than compounds CKR20, CKR25, and CKR31. CKR5 had been previously shown to be a potent PI3K kinase inhibitor. Therefore, we focused on compound CKR49.

**Figure 3.**
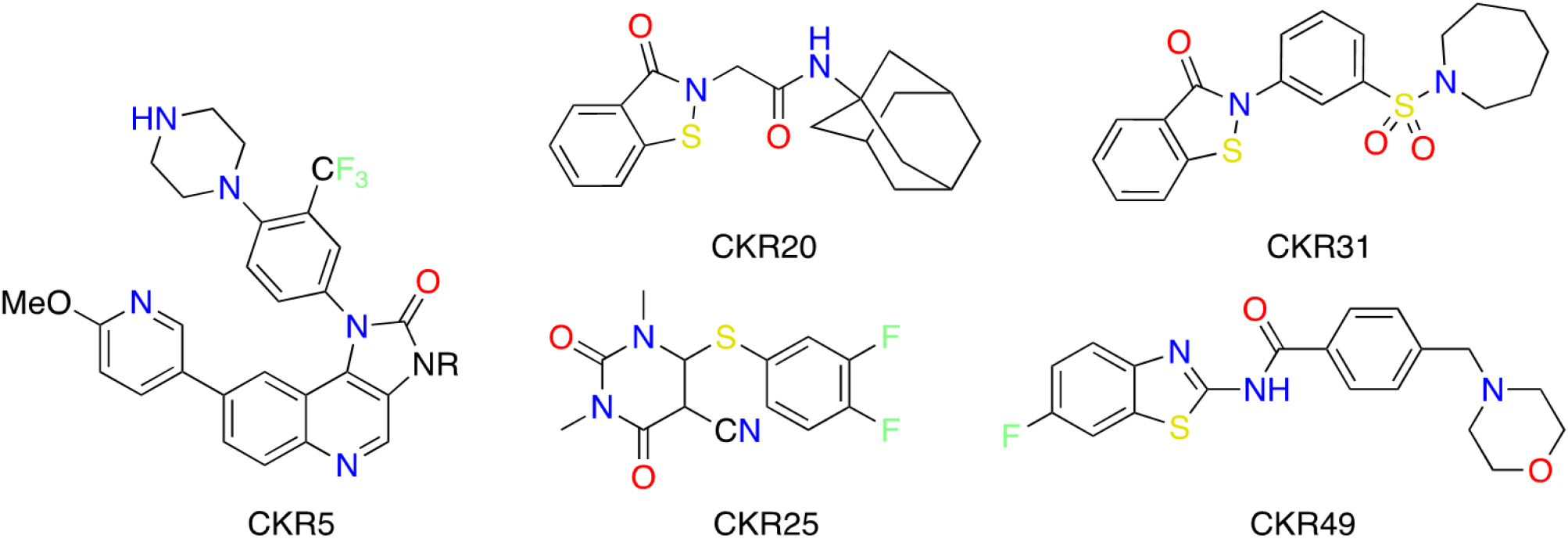
Structure of the primary hit compounds that increase phosphorylation of PS1 Ser367.

### Development of CKR49 analogs

We designed a series of compounds analogous to hit compound CKR49. We modified substituents in the thiazole ring, the -CH_2_-linker, and/or the morpholine ring. In some compounds, we also changed position of attachment of the sidechain on middle benzamide moiety from ‘*para*’ to ‘*meta*’ position (**Figure 4A**). The majority of compounds were synthesized in 2 (or 3 steps) using a general method described for compound CKR-49 (**Figure 4B**, Upp**er**). Thus, aminobenzo-thiazole derivative **Ia** underwent amide coupling with 4-chloromethylbenzoyl chloride, **IIa**, afforded product **IIIa**, and the latter reacted with morpholine **IVa** to afford CKR-49. Similarly, compound **IIIa** and analogous intermediates, prepared with various aminobenzothiazole derivatives and **IIa** or 3-chloromethylbenzoyl chloride reacted with **IVa** and analogous amines to afford analogs CKR49-1 – CKR49-9 and the Boc-protected intermediates for some compounds. Boc-protected intermediates underwent Boc-deprotection under acidic conditions to give free amine analogs of CKR49 (**SI Figure S1**). Several other compounds, including CKR49-10 – CKR49-12 were prepared by amide coupling of Ia with acid Va, Vb, or the analogous meta-substitutes acid, with or without Suzuki coupling with an appropriate boronic acid as described in **Figure 4B** (Lower). In this manner, we prepared 35 analogs of CKR49. For synthesis and characterization, please see SI Figure S1 (**SI Figure S1**).

**Figure 4.**
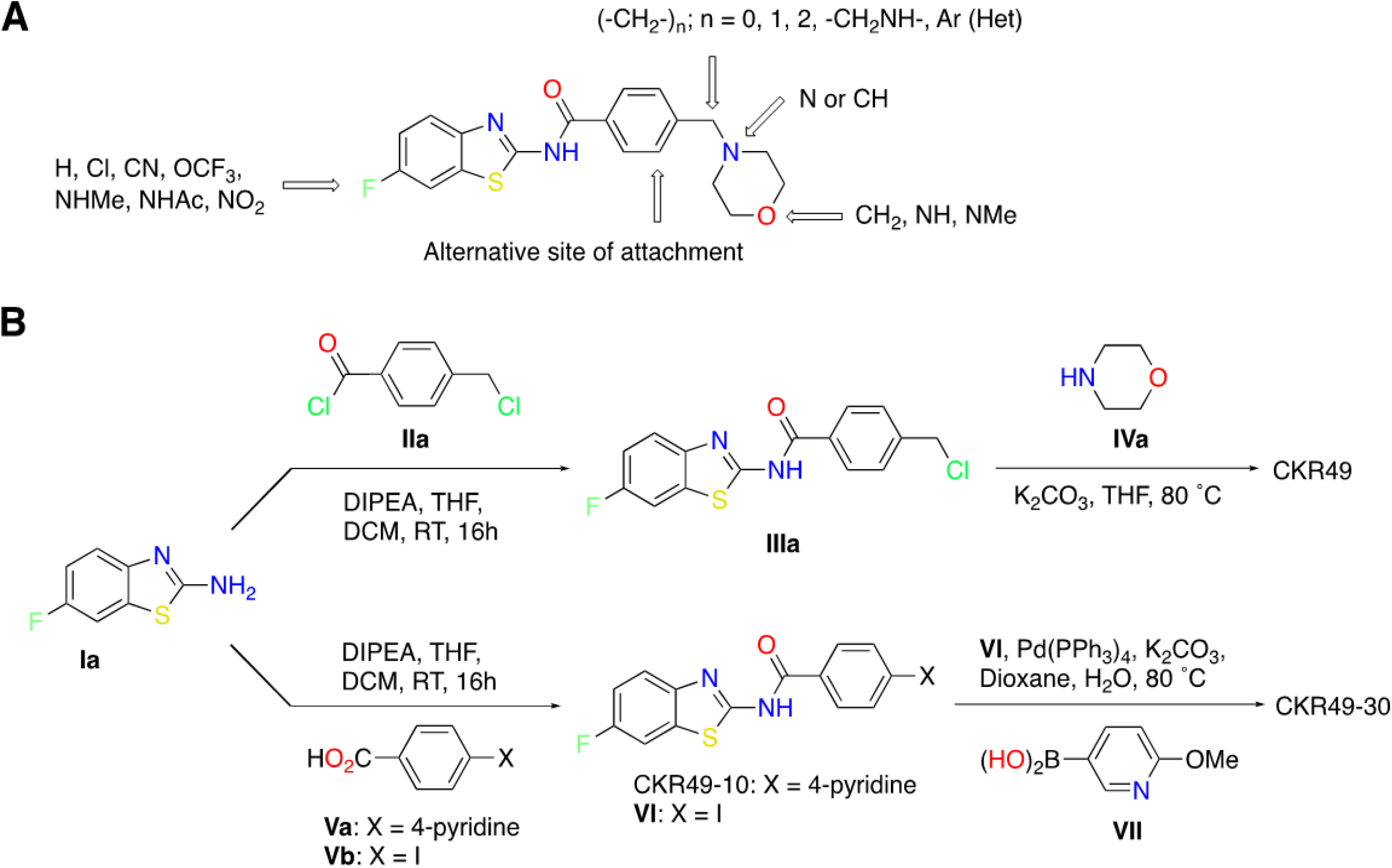
Design and synthesis of CKR49 and analogs. Shown in **A** is the general structure of CKR49 analogs and in **B** is a general scheme showing synthesis of CKR49 and analogs.

### CKR49 analogs increase PS1 Ser367 phosphorylation

Next, we evaluated all synthetic CKR49 analogs to determine their effects on PS1 Ser367 phosphorylation. For this study, we incubated recombinant PS1 3^rd^ loop with recombinant CK1γ2 kinase protein and ATP in presence and absence of the test compounds (0-128μM) for 0.5h in kinase buffer and measured phospho-Ser367 using antibody anti-phospho-Ser367 antibody in kinase buffer by Western Blot analysis. We identified 10 compounds that increase CK1γ2-catalyzed Ser367 phosphorylation by 200% at 64μM or lower concentration (**SI Table S-1**). Two compounds, CKR49-4 and CKR49-17 consistently showed higher activity among all the compounds, including the parent compound CKR49 (**Figure 5A and 5B**). It became evident that: a) free amine functions of 4-aminopiperidine in CKR49-4 and CKR49-17, b) ‘F’ or ‘CN’ substituent in benzothiazole ring, and c) -CH_2_-linker connecting the amine cyclic amine side chain to middle benzamide ring are desirable. We used compound CKR49-17 in subsequent experiments and determined its binding to CK1γ2 and its effects on C99 and Aβ production in cells in culture. The binding affinity of CKR49-17 to CK1γ2 was measured by performing microscale thermophoresis. This method is based on the directed movement of molecules in a temperature gradient, which strongly depends on a variety of molecular properties such as size, charge, hydration shell and conformation. An infrared laser induces a temperature gradient, and the directed movement of molecules through the temperature gradient is detected and quantified using either covalently attached or intrinsic fluorophores. By combining the precision of fluorescence detection with the variability and sensitivity of thermophoresis, this method can quantify the binding of small molecules to an enzyme. In our experiments, we found that CKR49-17 binds to CK1γ2 with a Kd of 180 nM (**Figure 5C**).

**Figure 5.**
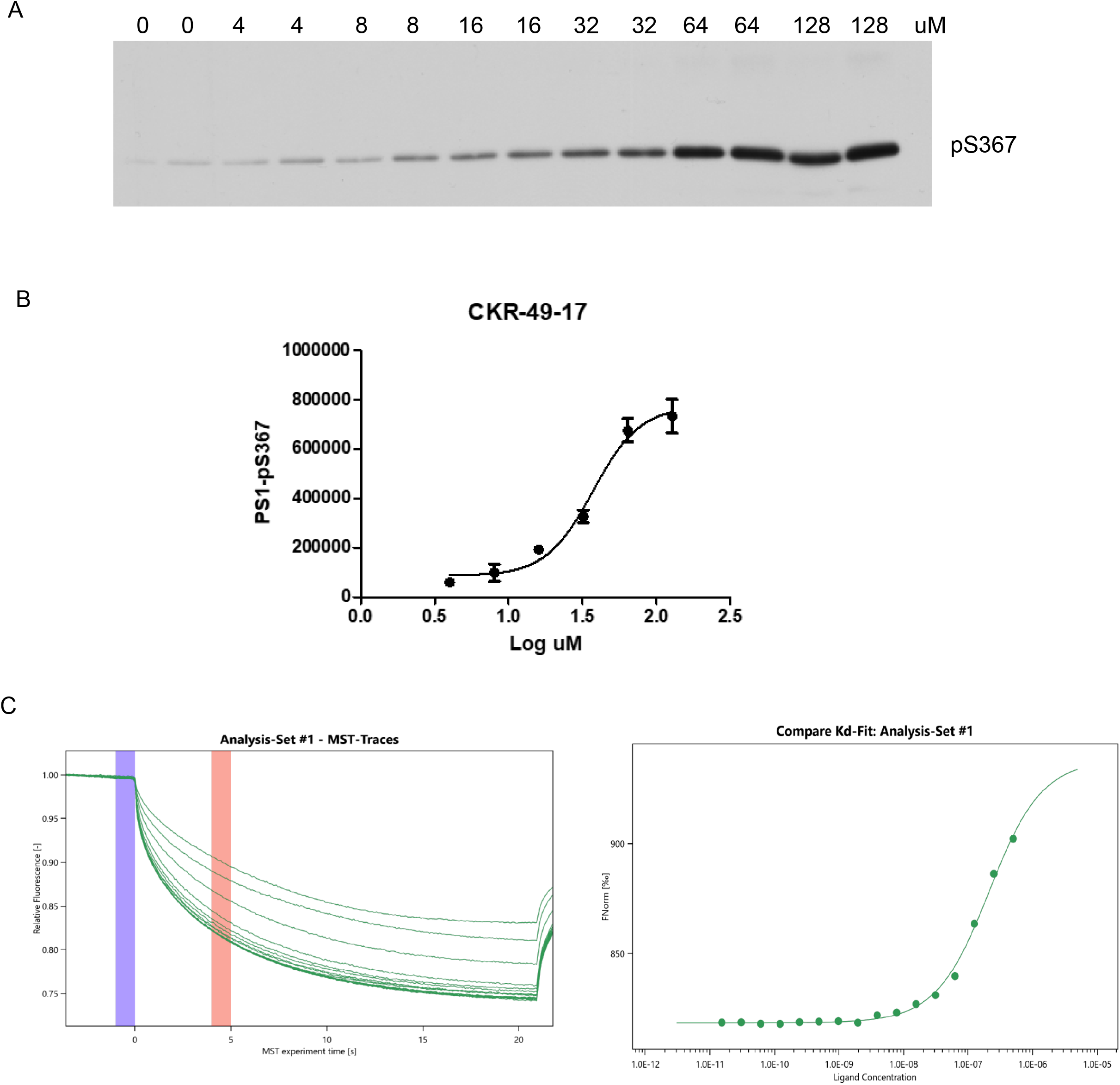
A) Effect of CKR49-17 on phosphorylation of PS1-pS367 in vitro. B) Quantification of the effect shown in A. C) Microscale thermophoresis between CKR49-17 and CK1γ2.

### Binding of CKR49-17 to an allosteric site allows the activation of CK1γ2

We hypothesized that CKR49 compounds could increase activity of CK1γ2 kinase through binding into an allosteric site in CK1γ2 kinase and inhibiting auto-phosphorylation. To test such possibility, a docking study was performed using compound CKR49-17 as well as several analogs of CKR49 and a known X-ray structure of CK1γ2 kinase (PDB: 2C47). Search for potential binding sites using its known X-ray structure was performed using the *Pocket Finder* function of LigandScout (InteLigand, Vienna). The pocket finder algorithm identifies putative pockets by creating a grid surface and calculating the buriedness value of each grid point on the surface. The iso surface obtained (grey or orange) represents empty space that may be suitable for creating a pocket. Thus, we identified two potential binding sites; however, only one site besides the catalytic site was large enough for CKR49 analogs to bind into. Each docking pose for CKR-49-17 was analyzed in their respective binding site. The estimated binding energy (kcal/mol) was calculated and the number of interacting features between the ligand and the protein were determined. These parameters were used to estimate the reliability of the docking pose observed. A preselection of the most relevant docking poses was made. Selection of the most relevant docking pose (from 19 poses) based on the score and the interacting features was performed and a pose ranking was attributed for comparison purpose.

Docking experiments with CKR-49-17 molecule were performed on each identified binding sites using autodock program (from LigandScout software). Several docking poses were generated for each binding site. After analysis of each pose, a selection of the most relevant poses was made. Finally, one docking pose was retained for each binding site and a pose ranking was proposed in order to compare different binding site poses. The most « promising » docking poses were observed for BS1 (catalytic site) and BS2 (potential allosteric site) on monomers A and C. It is expected that a small molecule interacting at the catalytic site would behave as an inhibitor, rather than an activator, so we focused in BS2, a potential allosteric site. BS2 on monomer C is located near the BS1 (catalytic site) and Asp 128 (**Figure 6A and 6B**). To validate this binding mode, a mutant CK1γ2 D128A was prepared. This mutant was catalytically active, but CKR49-17 was unable to activate this mutant, suggesting this binding mode is responsible for the activation of CK1γ2 (**Figure 6C**).

**Figure 6.**
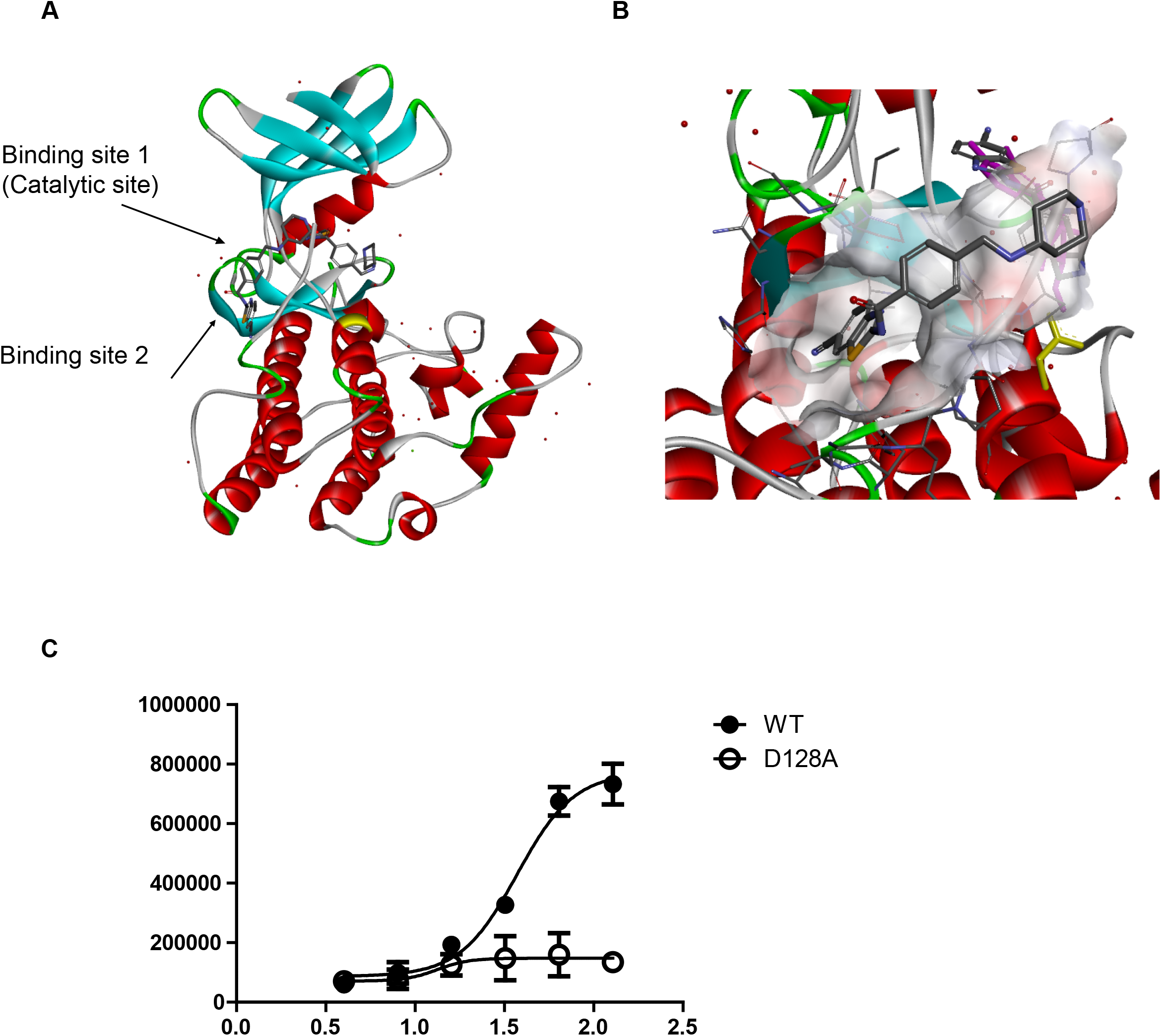
Allosteric site binding allows activation of CK1g2 by CKR49-17.Shown are in silico surface scanning of CK1γ2 kinase structure (PDB filed: 2C47) showing an allosteric cavity 1 in CK1γ2 kinase crystal structure, and (C) results of docking studies with compound CKR49 showing its strong binding into CK1 kinase allosteric site Cavity 1.

**Figure 7.**
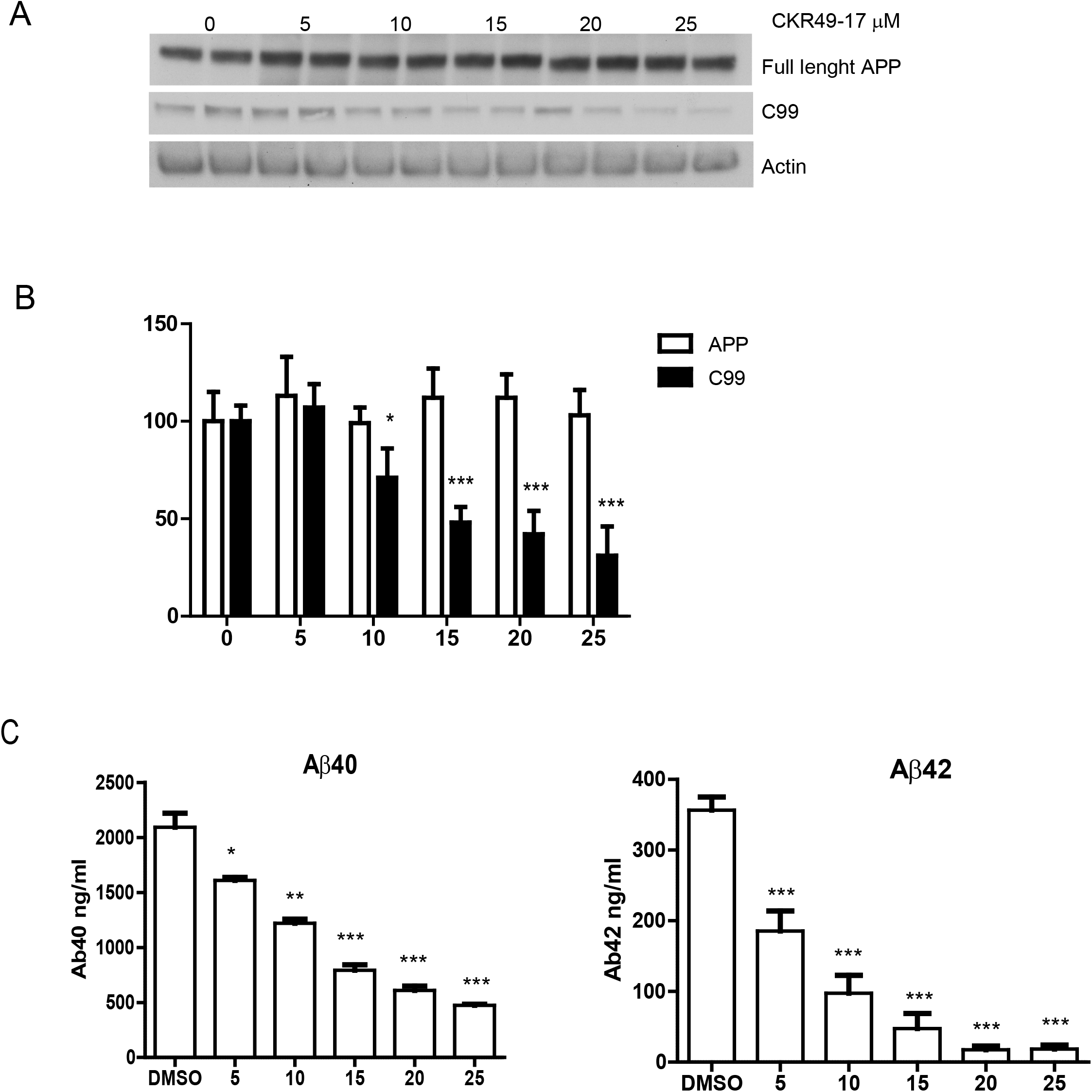
Effect of CKR49-17 on APP full length and C99 (A), quantification of A B) and effects on Aβ40 and Aβ42 levels (C) in N2A695 cells. Data for at least three experiments (mean ± SEM) are shown. ⋅, P < 0.05; ⋅⋅⋅, P < 0.001; two-tailed Student’s t test, 95% significance level.

### CKR49-17 reduces levels of C99 and Aβ peptides in N2A695 cells

To evaluate the effect of CKR49-17 on APP metabolism, N2A cells stably overexpressing APP-695 were incubated for five hrs. with increasing concentrations of CKR49-17. As shown in **Figure 6A and 6B**, CKR49-17 decreased C99 levels with an IC_50_ of 15 μM. Under these conditions, toxicity was not observed. Levels of full length APP remained stable at all concentrations tested (**Figure 6A and 6B**). In addition, CKR49-17 decreased levels of Aβ40 and Aβ42 (**Figure 6C**).

Extra:

Structure activity relationship (SAR)

In order to increase the PS1 phosphorylation at Ser367, the different structural modifications were done carefully in the benzamide and benzothiazole moieties of CKR-49. The results from PS1 phosphorylation suggested that CKR49 series compounds were remarkably influenced by various substituents on the benzamide ring and benzothiazole moiety as well. The data revealed that increasing methyl group between benzamide and morpholine(CKR49-23), the change of morpholino methyl group from para to meta position (CKR49-24), or replacing -F on benzothiazole moiety by -CN/Cl/OCF3 showed inactive on PS1 phosphorylation. Introducing of - NO2 group at the benzothiazole moiety showed active equally as CKR49 on PS1 phosphorylation and the reduction of -NO2 followed by acetylation (CKR49-18)/methylation (CKR49-19) showed inactive. The replacement of morpholine of CKR49 by piperidine (CKR49-9) did not induce the phosphorylation, however, introducing 4-aminopiperidine (CKR49-4) in place of morpholine displayed in increasing PS1 phosphorylation. Among the synthesized compounds, the modification carried out at position 6 by introducing cyano on the bezothiazole moiety and 4-aminopiperidine on the benzamide ring side (CKR49-17) exhibited the best results in PS1 phosphorylation but interestingly, none of the *N*-methylated piperidine compounds show active. The results showed that free amine of 4-aminopiperidine might be necessary for interacting with kinase. Substitution of piperazine/*N*-methyl piperazine in benzamide displayed inactive except the non-*N*-methylated compound CKR49-22. The pyridine substituents at the para/meta position of benzamide showed active, however, none showed much active than CKR49-17.

## Discussion

Protein kinase modulation has emerged as a potent strategy for drug development. Many kinase inhibitors have entered the clinic. However, the development of small molecule protein kinase activators has been elusive, even when there are several instances when this could be useful. An example is our finding that CK1γ2 phosphorylates Presenilin 1 in the gamma-secretase complex, leading to C99 and Aβ decrease. Here, we have reported the rational development of a CK1γ2 activator that decreases C99 and Aβ levels in cells in culture.

Regarding protein kinase activators, a small molecule screen identified pyrvinium, an FDA-approved anthelmintic drug, as an activator of CK1α that inhibits the Wnt/β-catenin pathway by increasing the phosphorylation of β-catenin, promoting its degradation ^12^. However, the exact mechanism by which pyrvinium activates CK1α is controversial. While reports have shown that pyrvinium allosterically activates CK1α to block Wnt/β-catenin signaling, others have shown that CK1α is not the *bona-fide* target of pyrvinium ^13-15^.

Concerning a potential involvement of CK1γ2 in AD, it has been reported that CK1γ2 gene is hypermethylated in vulnerable regions in the brain of AD patients^16^, which results in decreased CK1γ2 expression and could lead to decreased phosphorylation of PS1 at serine 367.

Studies with the protein kinase CK1α suggest that this kinase utilizes different structural domains to recognize different consensus phosphorylation sites^17^. Mutational experiments indicate that different regions of the enzyme participate in the recognition of substrate sequences. The autophosphorylation sites found here do not conform to canonical sites for the CK1 family^18-21^, suggesting that specific domains, different from the domains that recognize exogenous substrates could be involved in autophosphorylation. Our results indicate that phosphorylation of the regulatory site CK1γ2-Ser347 in the carboxy terminal domain occurs through an intramolecular mechanism. Therefore, this regulatory mechanism is independent of the concentration of CK1γ2. On the contrary, the phosphorylation of PS1-Ser367 depends on the concentration of both substrate and kinase.

CK1 family does not require T-loop phosphorylation to become active, as it is intrinsically active. As a consequence, this phosphorylation site and others, like the recently described T221 in CK1delta, have evolved to lower its kinase activity ^22^.

In our recombinant CK1γ2, many other phosphorylation sites were identified by mass spectrometry, and it remains to be elucidated the consequence of their phosphorylation.

An interesting observation is that the C-terminal domain of the CK1 family is the most divergent domain, suggesting that specific regulatory events modulate the different CK1 isoforms differently. Despite its divergence, CK1delta and epsilon show an inhibitory phosphorylation cluster. C-terminal inhibitory autophosphorylation could also be demonstrated for CK1γ1-3 as well as for CK1α and its splice variants CK1αL and CK1αS ^23,24^

Multiple levels of evidence indicate that C99 accumulation in vulnerable neurons contributes to neuronal death in AD ^25-34^. It has been shown that elevated levels of C99 in fibroblasts from Down syndrome and AD induce endosomal dysfunction by increasing the recruitment of APPL1 to rab5 endosomes, where it stabilizes active GTP-rab5, leading to accelerated endocytosis, endosome swelling and impaired axonal transport of rab5 endosomes ^25,26^. It has also been shown that C99 accumulation in the mouse brain leads to an increase in Cathepsin B and Lamp1 positive vesicles, as well as increased levels of LC3II and autophagic dysfunction ^29^.

Although the exact molecular mechanism by which C99 exert this effects is not yet known, it is becoming clear that C99 accumulation contributes to neuronal dysfunction and ultimately neuronal death.

One approach to lower C99 levels is BACE inhibition. Unfortunately, several clinical trials based on BACE inhibition were stopped due to safety concerns. For instance, atabacestat was stopped following concerning neuropsychiatric adverse events in phase2/3 trials. Verubecestat was stopped following worsening of cognitive function ^35^. One possibility is that BACE protease have more than 60 substrates, some of which may be important for maintaining cognitive abilities.^36^

Here we provide a proof of concept for an alternative approach for lowering C99 and Aβ levels.

Further work could search for brain penetrant CK1γ2 activators to test the efficacy of this approach *in vivo*.

## Acknowledgements

The authors acknowledge the Fisher Foundation for Alzheimer’s research (to PG and MF) and the Cure Alzheimer Fund (to VB, MF and SS) for their generous support. (ask Marc about funding). We also acknowledge Dr. Fraser Glickman for comments on the manuscript and the Fisher Drug Discovery Center at Rockefeller University.

## Methods

### Protein purification

For the CK1γ2 used in the high throughput screening, 500L of BL21 E. coli transformed with pGEX4-GST-CK1γ2 were grown in luria broth and induced for six hours using 100 μM isopropyl 1-thio-β-D-galactopyranoside, at the Bioexpression and Fermentation Facility at the University of Giorgia. Bacterial pellet was lysed in 50 mM Tris-HCl (pH 7.5), 150 mM NaCl, 1% Triton X-100, 2 mM EDTA, 0.1% β-mercaptoethanol, 0.2 mM PMSF, and 1 mM benzamidine and purified using glutathione-Sepharose™ 4B (GE Healthcare). Then GST was removed using 20 U/mg protein of Actev (Invitrogen) according to the instructions of the manufacturer.

### Autophosphorylation assay

In vitro assays were performed in a final volume of 50 μL in the following conditions: 50 mM Hepes pH 7.4, 150 mM NaCl, 10 mM MgCl_2_, 1 mM [γ^32^P] ATP (1,000 cpm/pmol), 100 ng of protein kinase CK1γ2. Reactions were performed at 30°C. Samples were loaded in a SDS/PAGE and autoradiography was performed in an Amersham Biosciences Typhoon 5 Variable Imager.

### High-throughput screening: Primary screen

Autophosphorylation inhibitor primary compound screening was carried out in 5Lμl final volume in 384-well flat bottom black polystyrene microplates (Lumitrac) by following any addition of reagents with a 30-s centrifugation at 180□×□*g* to ensure that all liquid was collected at the bottom of the well. The autophosphorylation reaction was composed of: 40 mM Tris pH 7.5, 20 mM MgCl2, 0.05% Tween 20, 0.1 mg/ml BSA, 30 μM ATP, 2.5 uM CK1γ2. Reagents were dispensed using a Thermo Multidrop Combi dispenser (Thermo Scientific). 0.05□μl of 5□mM stock compounds in DMSO were dispensed into assay microplates with a Janus 384 MDT NanoHead (PerkinElmer). Final concentration of the screening compounds in the assay was 16 μM. Column 23 was used to add just DMSO as a control and column 24 contained a CK1γ inhibitor. Plates were sealed using a Velocity11 PlateLoc thermal plate sealer, and incubated for 40 min at room temperature. The reaction was stopped by the addition of 15 ul of ADP-Glo reagent. The plates were incubated in a shaker for 40 min at room temperature. Then, 30 μl of kinase detection reagent were added, and plates were incubated for 60 min at room temperature. Levels of ATP were measured by the addition of 30 ul of kinase detection reagent. Plates were incubated in a shaker for 60 min at room temperature. Luminiscence was read in a Biotek Synergy Neo spectrophotomer instrument. Change in ATP levels were normalized against the positive control (column 23) and negative control (column 24) as follows: % inhibition □ = □ 100 □ × □ (RLU_sample_−RLU_average_negative_control_)/(RLU_average_positive_control_-

RLU_average_negative_control_). Compounds that inhibited CK1γ2 autophosphorylation activity by ≥40% were retested by a concentration response experiments to determine the half maximal inhibitory concentration (IC_50_). For this, stock compounds at 5 □ mM were serially diluted in half in DMSO for a total of ten dilutions. The same assay protocol as described above was used for validation of hit compounds but with the following differences: 0.1 □ μl of serially diluted compounds were dispensed with a Janus 384 MDT NanoHead (PerkinElmer) and the assay was performed 1 □ h after the addition of compounds. The highest final assay compound concentration was 32 □ μM. Subsequently, any compound selected for follow-up studies was reordered from available vending sources, dissolved in DMSO at 10 □ mM and re-tested in concentration response experiments. The Z′ factor of the primary screen ranged from 0.8 to 0.9.

### Chemical libraries

Library screening compounds were purchased from different vendors at different times over a period of 10 years using various compound selection strategies depending on factors such as cost per compound, budget, and diversity and drug-likeness (for example, Lipinski Filters, PAINS filters, QED scores). Compound vendors were contractually required to provide at least 85% purity, and provide LC-MS and NMR certificates on every compound. In some cases, the libraries were sold as existing sets with no option for removing undesirable substructures, and in other cases, the vendor offered the opportunity to select a subset of its very large commercially available compound collection. In cases where we had the option to select a subset of vendor compounds, we used the Biovia pipeline pilot software (http://accelrys.com/products/collaborative-science/biovia-pipeline-pilot). For example, we often selected a diverse number of compounds based on a yearly budget. We did this by filtering out the PAINS, non-Lipinski-like compounds and compounds similar to the existing collection from vendor files by using Tanimoto similarity calculations and then applying fingerprint-based (FCFP6) clustering algorithms, which parsed the compounds into groups of similar structures. The average group size was set to a number of groups that we could afford to purchase in a particular time, and only those compounds positioned in the center of clusters were purchased in order to maximize diversity. Supplementary Table X shows the number of compounds purchased from each vendor, and the smiles string of every compound screened is included along with the screening results deposited into NCBI screening library database. A total of 281,348 unique compounds were studied in the primary screen. Compounds were dissolved in DMSO to 5 mM in 96-well tube arrays and then partitioned into ten aliquots of 20 μl each and two master copies of 80 μl each into 384-well polypropylene deep small volume microplate (Greiner Bio-One), which were stored at −28 °C. Only a single 20-μl copy was used as a compound source plate from which to draw compound into the assay plate, to a maximum of 20 freeze–thaw cycles.

### Chemical Synthesis

A) CKR49-H, **5** (or **5a**). We prepared **5a** by reacting the commercially available aminobenzo-thiazole derivative **6a** with 4-chloromethylbenzoyl chloride, **7a**, followed by treatment of the resulting product **8a** with morpholine, **9a** (Scheme 1). Similarly, we prepared other analogs of **5a**, including **5b**-**y** via intermediates **8a**-**c** and analogs, and reacting these with various amines.

We also prepared analogs of **5a**, including **8f**-**n**, in one step by reacting **6a**-**b** and other amino-benzothiazole derivatives with m-or p-functionalized benzoic acid derivatives, such as **7c** and **7d**. A list of all new analogs of **5a**, including **5b**-**y** and **8f**-**n**, and their effects on PS1 phosphorylation is summarized in Table 1.

### Cell culture

Mouse N2A neuroblastoma WT cells and an established N2A cell line expressing APP695 were maintained in medium containing 50% DMEM and 50% Opti-MEM, supplemented with 5% FBS and 200 μg/mL G418 (Life Technologies). These cell lines were tested for mycoplasma contamination using LookOut mycoplasma PCR detection (Sigma).

### Aβ measurement

The quantitative analysis of Aβ40 and Aβ42 was performed using an ELISA kit according to the manufacturer’s instructions (Life Technologies). Briefly, supernatants of conditioned media from N2A/APP695 cells were diluted in buffer and incubated in a 96-well plate with Aβ40 or Aβ42 antibody for 3 h at room temperature. After washes of unbound material, a secondary antibody conjugated to horseradish peroxidase (HRP) was added for 30 min. Unbound secondary antibody was washed, and samples were subsequently incubated with a developing reagent for 30 min. Stop solution was added to block further reaction between HRP and the colorimetric substrate. An absorbance multiplate reader was used to quantify the colorimetric reaction at 450 nm

### Protein Quantification and Immunoblot Analysis

Cultured cells or brain tissues were lysed in buffer A [50 mM Tris⋅HCl (pH 7.5), 150 mM NaCl, and 2 mM MgCl] supplemented with 1% Triton X-100, a protease inhibitor mixture (Complete-EDTA free; Roche), and phosphatase inhibitors (30 mM NaF, 1 mM orthovanadate, and 30 mM pyrophosphate). The cell or tissue lysates were disrupted with a probe-type sonicator for 10 s twice and centrifuged, and the protein levels in the supernatant were measured by the bicinchoninic acid assay method. For Aβ40, Aβ42, and sAPPβ, conditioned cell culture media were analyzed. The samples were boiled in standard protein sample buffer; subjected to SDS/PAGE, followed by protein transfer onto a PVDF membrane; and incubated overnight at 4 °C with the following antibodies: anti-PS1 (mouse monoclonal, 1:1,000; EMD Millipore) and anti-APP (6E10, mouse monoclonal, 1:1,000; Covance) for total APP and βCTF, and antiactin (rabbit polyclonal, 1:1,000; Santa Cruz Biotechnology), anti-LC3 (rabbit polyclonal, 1:1,000; Sigma), anti-p62 (rabbit polyclonal, 1:500; Cell Signaling Technology), anti-CK1γ2 (rabbit polyclonal, 1:500; Abcam), and anti–PS1-NTF (a polyclonal antibody generated in our laboratory).

### Microscale thermophoresis

The MST assay was carried out on a Monolith NT.115 (NanoTemper Technologies). CK1γ2 was prepared at concentrations from 0.305 to 10000□nM and fluorescence dye from the Monolith His-tag Labeling kit RED-tris-NTA (NanoTemper Technologies) were added to concentrations of 80□nM and 5□nM, respectively, in 10□mM HEPES-NaOH (pH 7.5), 150□mM NaCl, 1□mM DTT, 1□mM CaCl_2_, 0.005% Tween 20, and 5% DMSO, and incubated for 30□min□at 25□°C. Samples were loaded into Monolith NT.115 Premium Capillaries (NanoTemper Technologies) and subjected to the MST measurement. Thermophoresis was measured using the IR laser at 20% MST power. MST data was analyzed in NT Analysis software (NanoTemper Technologies)

