## Supplemental Table S1 for "Rational development of a small-molecule activator of CK1γ2 that decreases C99 and beta-amyloid levels"

**Table S-1.** Structure of compound CKR49 analogs. Doubled pPS1 refers to the concentration of compound necessary to increase the phosphorylation fo pPS1 by 100%.

| **Compound ID** | **Structure** | **Doubled pPS1** |
| --- | --- | --- |
| CKR49-H | 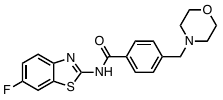 | 64 µM |
| CKR49-1 | 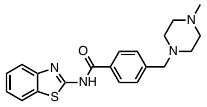 | Inactive |
| CRR49-2 | 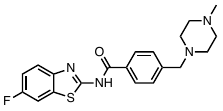 | Inactive |
| CKR49-3 | 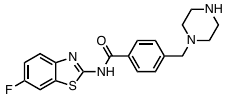 | Inactive |
| CKR49-4 | 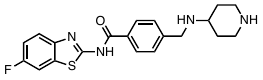 | 16 µM |
| CKR49-5 | 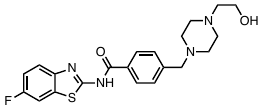 | Inactive |
| CKR49-6 | 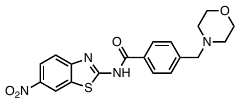 | 64 µM |
| CKR49-7 | 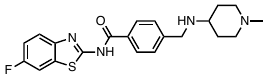 | Inactive |
| CKR49-8 | 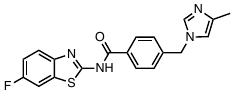 | Inactive |
| CKR49-9 | 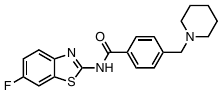 | Inactive |
| CKR49-10 | 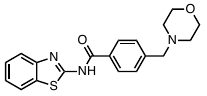 | Inactive |
| CKR49-11 | 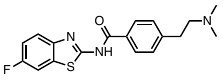 | Inactive |
| CKR49-12 | 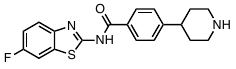 | Inactive |
| CKR49-13 | 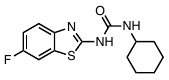 | Inactive |
| CKR49-14 | 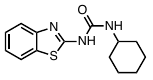 | Inactive |
| CKR49-15 | 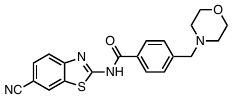 | Inactive |
| CKR49-16 | 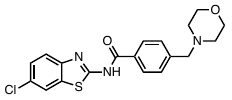 | Inactive |
| CKR49-17 | 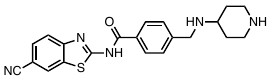 | 6 µM |
| CKR49-18 | 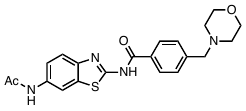 | Inactive |
| CKR49-19 | 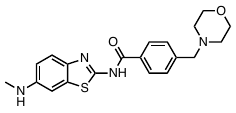 | Inactive |
| CKR49-20 | 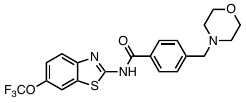 | Inactive |
| CKR49-21 | 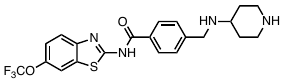 | 16 µM |
| CKR49-22 | 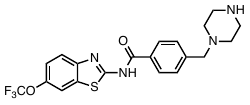 | 20 µM |
| CKR49-23 | 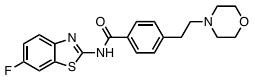 | 32 µM |
| CKR49-24 | 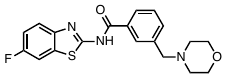 | Inactive |
| CKR49-25 | 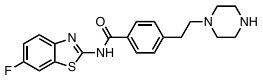 | Inactive |
| CKR49-26 | 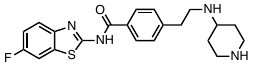 | Inactive |
| CKR49-27 | 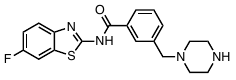 | Inactive |
| CKR49-28 | 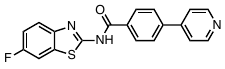 | 16 µM |
| CKR49-29 | 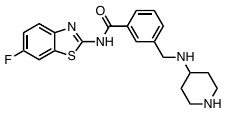 | 24 µM |
| CKR49-30 | 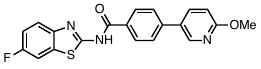 | 64 µM |
| CKR49-31 | 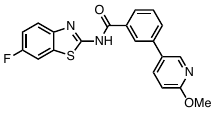 | 32 µM |
| CKR49-32 | 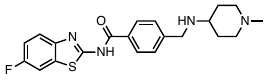 | Inactive |
| CKR49-33 | 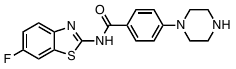 | Inactive |
| CKR49-34 | 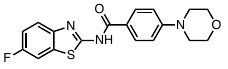 | Inactive |
| CKR49-35 | 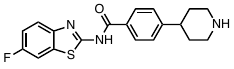 | Inactive |
| CKR49-36 | 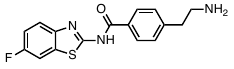 | Inactive |
| CKR49-37 | 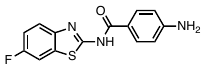 | Inactive |
| CKR49-38 | 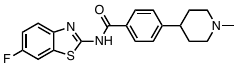 | Inactive |
| CKR49-39 | 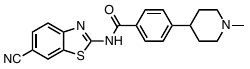 | Inactive |
| CKR49-40 | 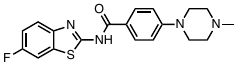 | Inactive |
| CKR49-41 | 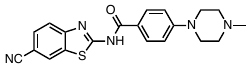 | Inactive |
| CKR49-42 | 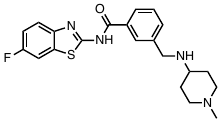 | Inactive |
