## Supplemental Figure S1 for "Rational development of a small-molecule activator of CK1γ2 that decreases C99 and beta-amyloid levels"

Supplementary Figure 1: Synthesis of CKR49 and analogues.

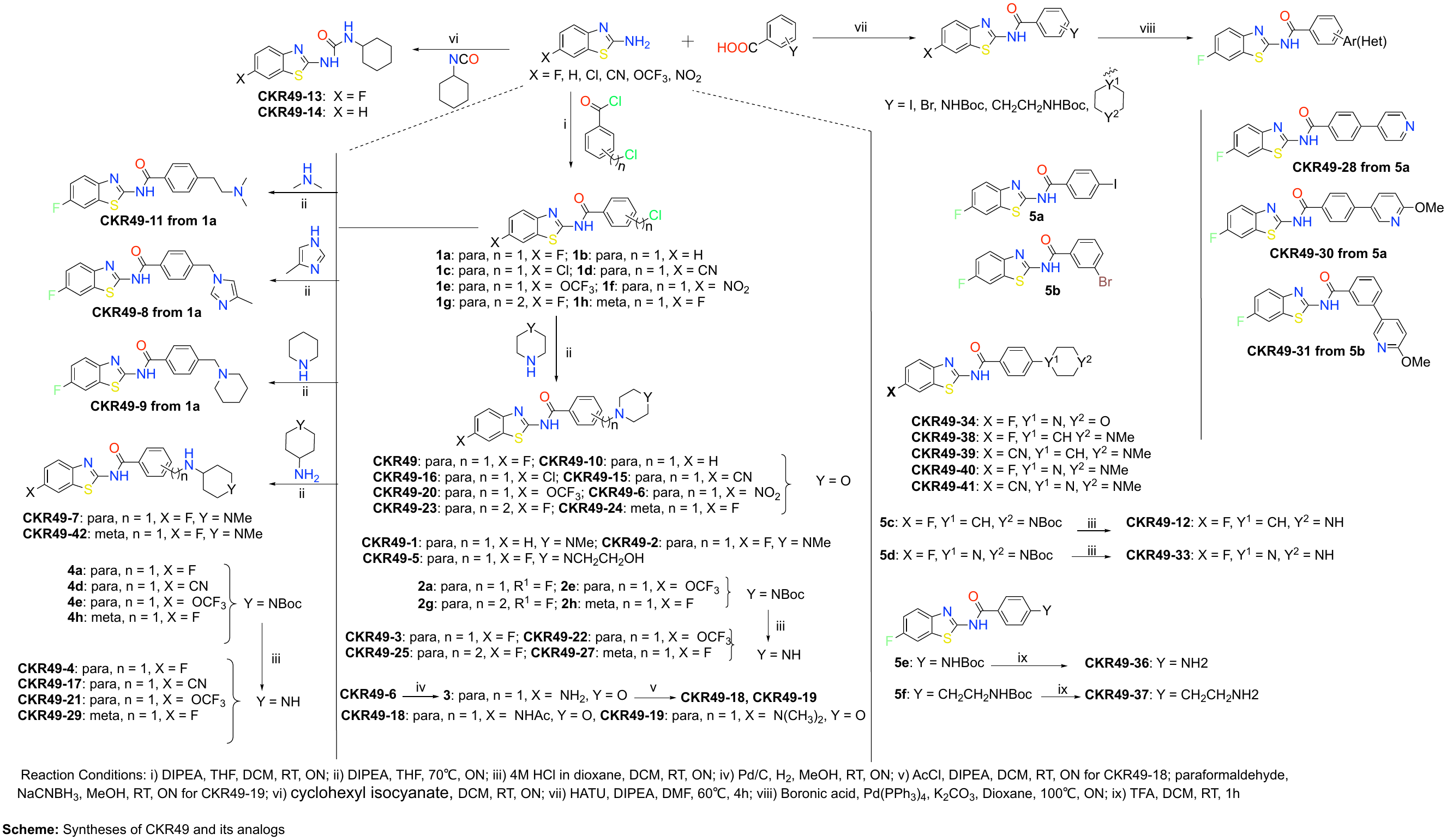

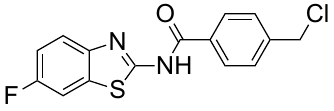
**4-(Chloromethyl)-*N*-(6-fluorobenzothiazol-2-yl)benzamide (1a).**

To a stirred solution of 2-Amino-6-fluorobenzothiazole (582 mg, 3.46 mmol) in THF (10 mL) at 0 °C, DIPEA (1.2 mL, 6.92 mmol) was added and stirred for 5 minutes. 4-(chloromethyl) benzoyl chloride (982 mg, 5.19 mmol) in CH_2_Cl_2_ (2 mL) was then slowly added to the mixture and stirred at room temperature for ON. The reaction mixture was quenched with H_2_O (4 mL) and extracted with ethyl acetate (50 mL). The organics were washed with brine, dried (Na_2_SO_4_), filtered and concentrated in vacuo to obtain a crude residue which was directly used for the next steps.

**Syntheses of 1b-1h.**

**1b-1h** compounds were synthesized as described in **1a** with the corresponding 2-aminobenzothiazole and benzoyl chloride. All the compounds were used directly for the next steps.

***N*-(6-Fluorobenzothiazol-2-yl)-4-(morpholinomethyl)benzamide (CKR-49).**

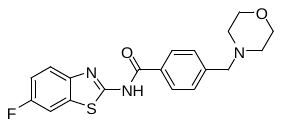

To a crude residue of compound **1a** (221 mg, 0.69 mmol) in THF (3 mL), morpholine (180 ul, 2.07 mmol) and DIPEA (180 ul, 1.035 mmol) were added, then sealed the round bottom flask with the septum and stirred at 70 °C for ON. The solvent was evaporated under vacuo, added cold ethyl acetate and hexane mixture, sonicated and filtered to afford **CKR49** (199 mg, 78%). ^1^H NMR (600 MHz, CDCl_3_): *δ* 10.22 (bs, 1H), 7.93 (d, *J* = 8.1 Hz, 2H), 7.56 - 7.472 (m, 4H), 7.10 (td, *J* = 8.7, 2.3 Hz, 1H), 3.76 -3.67 (m, 4H), 3.56 (s, 2H), 2.48 – 2.42 (m, 4H) ppm

**Syntheses of 2a, 2e, 2g, 2h, 4a, 4d, 4e, 4h, CKR49 - 1, 2, 5-11, 15, 16, 20, 23, 24 and 42.**

The above-mentioned compounds were synthesized as described in method **CKR49** with the corresponding Chloromethyl-*N*-(6-fluorobenzothiazol-2-yl)benzamide and the amine. All the compounds were purified either by flash column chromatography or washed with an appropriate cold solvent for solids to get a pure compound as mentioned in **CKR49**. All the compounds were analyzed using NMR or LCMS and all the final compounds were confirmed their purity by LCMS UV chromatogram at 260 nm before testing their activity.

**2a.**

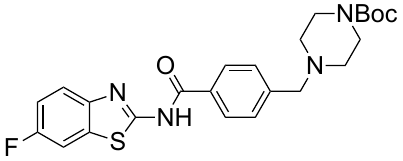

^1^H NMR (600 MHz, CDCl_3_): *δ* 10.07 (s, 1H), 7.92 (d, *J* = 8.1 Hz, 1H), 7.58 - 7.52 (m, 2H), 7.49 (d, *J* = 7.8 Hz, 2H), 7.12 (td, *J* = 8.8, 2.4 Hz, 1H), 3.57 (s, 2H), 3.44 (bs, 4H), 2.39 (bs, 4H), 1.46 (s, 9H) ppm

**2e.**

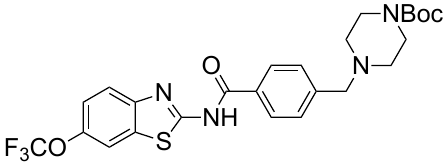

^1^H NMR (600 MHz, CDCl_3_): *δ* 10.05 (bs, 1H), 7.94 (d, *J* = 8.0, 2H), 7.72 (bs, 1H), 7.66 (d, *J* = 8.6 Hz, 1H), 7.51 (d, *J* = 7.9 Hz, 2H), 7.29 – 7.26 (m, 1H), 3.58 (s, 2H), 3.47 -3.41 (m, 4H), 2.44 – 2.37 (m, 4H), 1.45 (s, 9H) ppm

**2g.**

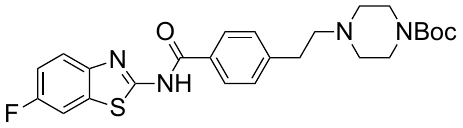

^1^H NMR (600 MHz, CDCl_3_): *δ* 9.96 (bs, 1H), 7.89 (d, *J* = 7.9, 2H), 7.61 (dd, *J* = 8.8, 4.6, 1H), 7.53 (dd, *J* = 7.9, 2.7 Hz, 1H), 7.36 (d, *J* = 8.0 Hz, 2H), 7.14 (td, *J* = 8.8, 2.7, 1H), 3.47 – 3.42 (m, 4H), 2.88 (t, J = 7.6, 2H), 2.63 (t, *J* = 8.4, 2H), 2.49 – 2.43 (m, 4H), 1.46 (s, 9H) ppm

**4a.**

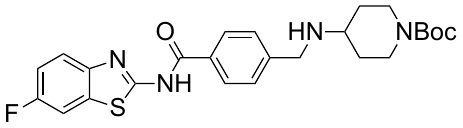

^1^H NMR (600 MHz, CDCl_3_): *δ* 7.92 (d, *J* = 8.0 Hz, 2H), 7.62 - 7.58 (m, 1H), 7.54 (dd, *J* = 8.0, 2.4 Hz, 1H), 7.50 (d, *J* = 7.49 Hz, 2H), 7.14 (td, *J* = 8.9, 2.5 Hz, 1H), 4.10 - 3.96 (m, 2H), 3.91 (s, 2H), 2.88 - 2.76 (m, 2H), 2.70 - 2.63 (m, 1H), 1.91-1.82 (m, 2H), 1.46 (s, 9H), 1.34 - 1.27 (m, 2H) ppm

**4d.**

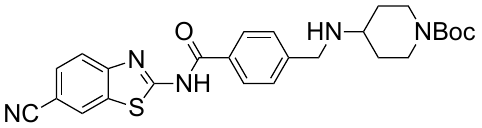

^1^H NMR (600 MHz, CDCl_3_): *δ* 8.18 (s, 1H), 7.97 (d, *J* = 8.1 Hz, 2H), 7.80 (d, *J* = 8.3 Hz, 1H), 7.69 (dd, *J* = 8.3, 1.2 Hz, 1H), 7.55 (d, *J* = 7.9 Hz, 2H), 4.11 - 3.97 (m, 2H), 3.94 (s, 2H), 2.87 - 2.75 (m, 2H), 2.72 - 2.63 (m, 1H), 1.90 -1.83 (m, 2H), 1.45 (s, 9H), 1.36 - 1.27 (m, 2H) ppm

**4e.**

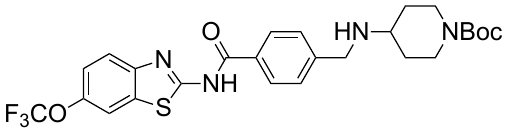

^1^H NMR (600 MHz, CDCl_3_): *δ* 7.95 (d, *J* = 8.1, 2H), 7.72 (bs, 1H), 7.67 (d, *J* = 8.8 Hz, 1H), 7.51 (d, *J* = 8.1 Hz, 2H), 7.30 – 7.27 (m, 1H), 4.12 - 3.96 (m, 2H), 3.91 (s, 2H), 2.87 - 2.74 (m, 2H), 2.70 - 2.62 (m, 1H), 1.91 -1.81 (m, 2H), 1.45 (s, 9H), 1.36 - 1.26 (m, 2H) ppm

**CKR49-1.**

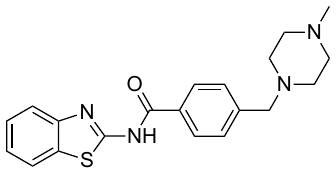

^1^H NMR (600 MHz, CDCl_3_): *δ* 7.93 (d, *J* = 8.2 Hz, 2H), 7.85 (d, *J* = 7.6 Hz, 1H), 7.56 (d, *J* = 7.8 Hz, 1H), 7.46 (d, *J* = 7.9 Hz, 2H), 7.35 (t, *J* = 6.9 Hz, 1H), 7.31 (t, *J* = 7.6 Hz, 1H), 3.58 (s, 2H), 2.56 (bs, 8H), 2.38 (s, 3H) ppm

**CKR49-2.**

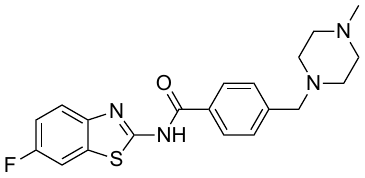

^1^H NMR (600 MHz, CDCl_3_): *δ* 10.22 (bs, 1H), 7.92 (d, *J* = 8.1 Hz, 1H), 7.53 (m, 2H), 7.48 (d, *J* = 8.0 Hz, 2H), 7.10 (td, *J* = 8.8, 2.5 Hz, 1H), 3.59 (s, 2H), 2.59 (bs, 8H), 2.40 (s, 3H) ppm

**CKR49-6.**

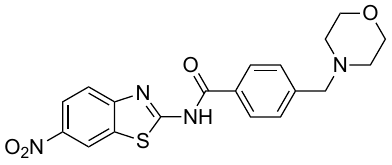

^1^H NMR (600 MHz, CDCl_3_): *δ* 8.79 (d, *J* = 2.1 Hz, 1H), 8.31 (dd, *J* = 8.9, 2.1, 1H), 7.95 (d, *J* = 8.24, 2H), 7.77 (d, *J* = 8.9, 1H), 7.55 (d, *J* = 8.0, 2H), 3.73 (t, *J* = 4.5, 4H), 3.59 (s, 2H), 2.46 (m, 4H) ppm

**CKR49-9.**

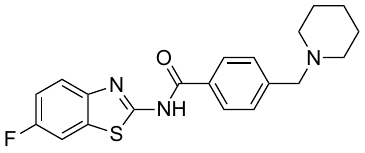

^1^H NMR (600 MHz, CDCl_3_): *δ* 9.95 (bs, 1H), 7.95 *-* 7.89 (m, 2H), 7.63 - 7.47 (m, 4H), 7.18 - 7.10 (m, 1H), 3.55 (s, 2H), 2.39 (bs, 4H), 1.59 (bs, 4H), 1.46 (bs, 2H) ppm

**CKR49-15.**

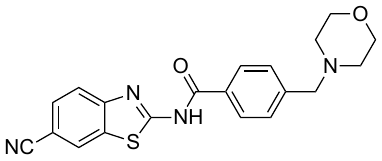

^1^H NMR (600 MHz, CDCl_3_): *δ* 10.05 (bs, 1H), 8.18 (s, 1H), 7.94 (d, *J* = 8.2, 2H), 7.73 – 7.70 (m, 1H), 7.67 – 7.64 (m, 1H), 7.53 (d, *J* = 7.9, 2H), 3.76 – 3.70 (m, 4H), 3.58 (s, 1H), 2.50 – 2.43 (m, 4H) ppm

**CKR49-16.**

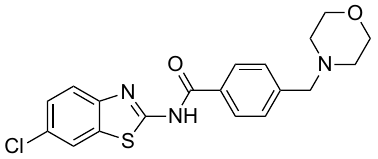

^1^H NMR (600 MHz, CDCl_3_): *δ* 9.94 (bs, 1H), 7.92 (d, *J* = 8.0 Hz, 2H), 7.83 (d, *J* = 1.8 Hz, 1H), 7.57 (d, *J* = 8.6 Hz, 1H), 7.52 (d, *J* = 7.8 Hz, 2H), 7.36 (dd, J = 8.6, 1.9, 1H), 3.74 – 3.71 (m, 4H), 3.57 (s, 2H), 2.49 – 2.43 (m, 4H) ppm

**CKR49-20.**

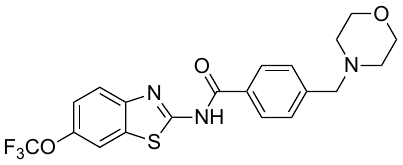

^1^H NMR (600 MHz, CDCl_3_): *δ* 10.31 (bs, 1H), 7.94 (d, *J* = 7.7 Hz, 2H), 7.72 (bs, 1H), 7.57 (dd, *J* = 8.6, 1.9 Hz, 1H), 7.50 (d, *J* = 7.7 Hz, 2H), 7.25 - 7.22 (m, 1H), 3.75 – 3.68 (m, 4H), 3.56 (s, 2H), 2.50 – 2.40 (m, 4H) ppm

**CKR49-23.**

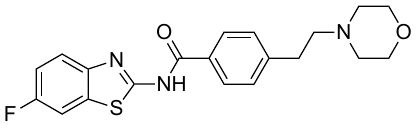

^1^H NMR (600 MHz, CDCl_3_): *δ* 9.97 (bs, 1H), 7.89 (d, *J* = 8.0, 2H), 7.60 (dd, *J* = 8.7, 4.5, 1H), 7.53 (dd, *J* = 8.0, 2.7 Hz, 1H), 7.36 (d, *J* = 7.9 Hz, 2H), 7.14 (td, *J* = 8.9, 2.7, 1H), 3.76 – 3.71 (m, 4H), 2.88 (t, J = 7.6, 2H), 2.62 (t, *J* = 8.4, 2H), 2.55 – 2.50 (m, 4H) ppm

**Syntheses of CKR49 - 3, 4, 12, 17, 21, 22, 25, 27, 29, 33, 35-37.**

The corresponding Boc protected compound was taken in CH_2_Cl_2_ (4 mL), 4M hydrogen chloride in dioxane (0.5 mL) or trifluoroacetic acid (1mL) was added at 0 C° and stirred for ON at room temperature. The solvent was evaporated under vacuo, added a few drops of MeOH, sonicated, filtered, washed with cold ether and dried to afford the free amine compound.

***N*-(6-Aminobenzothiazol-2-yl)-4-(morpholinomethyl)benzamide (3).** The solution of **CKR49-6** (360 mg, 0.90 mmol) in 50% MeOH/Ethyl Acetate (20 mL) was purged with nitrogen and then 10% Pd/C was catalytically added and the mixture was stirred for 3 hours under H_2_ (Balloon). The mixture was filtered through celite, concentrated in vacuo to get the crude residue and used directly for the next steps.

***N*-(6-Acetamidobenzothiazol-2-yl)-4-(morpholinomethyl)benzamide (CKR49-18).** To a stirred solution of **3** (40 mg, 0.11 mmol) in CH_2_Cl_2_ (3 mL), DIPEA (26 ul, 0.16 mmol) and acetyl chloride (12 ul, 0.16 mmol) were added at 0 °C and then the reaction mixture was stirred at rt for ON. The solvent was removed under vacuo and the crude was dissolved in a few drops of MeOH, sonicated, filtered and dried to afford **CKR49-18** as a solid (29 mg, 70%). The compound was analyzed using LCMS and the purity was checked by UV at 260 nm in LCMS.

***N*-(6-(Methylamino)benzothiazol-2-yl)-4-(morpholinomethyl)benzamide(CKR49-19).** To a stirred solution of **3** (40 mg, 0.11 mmol) in CH_2_Cl_2_ (3 mL), DIPEA (26 ul, 0.16 mmol) and acetyl chloride (12 ul, 0.16 mmol) were added at 0 °C and then the reaction mixture was stirred at rt for ON. The solvent was removed under vacuo and the crude was dissolved in a few drops of MeOH, sonicated, filtered and dried to afford **CKR49-19** as a solid (28 mg, 73%). The compound was analyzed using LCMS and the purity was checked by UV at 260 nm in LCMS.

**1-Cyclohexyl-3-(6-fluorobenzothiazol-2-yl)urea (CKR49-13).**

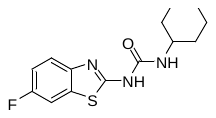

To a stirred solution of 2-amino-6-fluoroobenzothiazole (110 mg, 0.654 mmol) in CH_2_Cl_2_ (5 mL), cyclohexyl isocyanate (126 ul, 0.982 mmol) was added at 0 °C and then the reaction mixture was stirred at rt for ON. The solvent was removed under vacuo and the crude was dissolved in a few drops of MeOH, sonicated, filtered and dried to afford **CKR49-13** as a solid (119 mg, 62%). ^1^H NMR (600 MHz, CDCl_3_): *δ* 10.16 (bs, 1H), 7.69 - 7.64 (m, 1H), 7.45 - 7.41 (m, 1H), 7.16 - 7.10 (m, 1H), 3.86 - 3.75 (m, 1H), 2.05 - 1.97 (m, 2H), 1.80 - 1.72 (m, 2H), 1.66 - 1.57 (m, 2H), 1.49 - 1.39 (m, 2H), 1.38 - 1.30 (m, 2H) ppm

**1-(Benzothiazol-2-yl)-3-cyclohexylurea (CKR49-14).**

The same reaction described was used to prepare compound **CKR49-13**. 2-aminobenzothiazole (98 mg, 0.653 mmol), cyclohexyl isocyanate (126 ul, 0.982 mmol) and CH_2_Cl_2_ (5 mL). The solvent was removed under vacuo and the crude was dissolved in a few drops of MeOH, sonicated, filtered and dried to afford **CKR49-14** as a solid (106 mg, 59%).

***N*-(6-fluorobenzothiazol-2-yl)-4-iodobenzamide (5a)**.

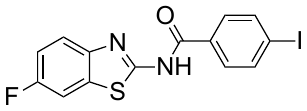

To a stirred solution of 2-amino-6-fluoroobenzothiazole (455 mg, 2.71 mmol) in DMF (5 mL), 4-Iodobenzoic acid (673 mg, 2.71 mmol), HATU (1.57 g, 4.07 mmol) and DIPEA (707 ul, 3.96 mmol) were added and stirred at 60 °C for 4 hours. DMF mostly was evaporated under vacuo, then ice was added to the mixture, sonicated the crude mixture and filtered. The filtrate was taken in CH_2_Cl_2_, then the organics were washed with brine, dried (Na_2_SO_4_), filtered and concentrated in vacuo to obtain **5a**.

^1^H NMR (600 MHz, CDCl_3_): *δ* 7.98 - 7.92 (m, 2H), 7.76 - 7.73 (m, 3H), 7.61 - 7.53 (m, 1H), 7.24 - 7.18 (m, 1H) ppm

**Syntheses of 5b-5f, CKR49 - 34, 38-41.**

The above-mentioned compounds were synthesized using HATU as described in method **5a** with the corresponding 2-amino-6-fluoroobenzothiazole and acid. All the compounds were purified either by flash column chromatography or washed with an appropriate cold solvent for solids to get a pure compound. All the compounds were analyzed using NMR or LCMS and all the final compounds were confirmed their purity by LCMS UV chromatogram at 260 nm before testing their activity.

***N*-(6-fluorobenzothiazol-2-yl)-4-(6-methoxypyridin-3-yl)benzamide (CKR49-30).**

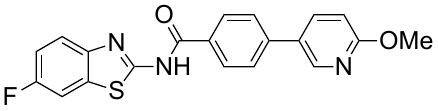

The compound 5a (201 mg, 0.51 mmol) was taken in 1,4-Dioxane (4mL). (6-methoxypyridin-3-yl)boronic acid (110 mg, 0.76 mmol) was added and then purged with nitrogen. Pd(PPh_3_)_4_ (30 mg, 0.03 mmol) and 2N K_2_CO3 (1mL) were added and the reaction mixture was irradiated in microwave at 100 °C for 30 minutes. The reaction mixture was poured into cold water, sonicated and filtered. The filtrate was taken in CH_2_Cl_2_, then the organics were washed with brine, dried (Na_2_SO_4_), filtered and concentrated in vacuo to obtain the crude residue. The residue was purified by flash column chromatography to afford **CKR49-30** as a solid (151 mg, 78%).

^1^H NMR (600 MHz, CDCl_3_): *δ* 10.27 (bs, 1H), 8.43 (d, *J* = 2.4, 1H), 8.05 (d, *J* = 8.2, 2H), 7.82 (dd, *J* = 6.2, 2.3 Hz, 1H), 7.67 (d, *J* = 8.1 Hz, 2H), 7.58 – 7.52 (m, 2H), 7.1 (td, *J* = 8.9, 2.3, 1H), 6.86 (d, *J* = 8.5, 1H), 4.00 (s, 3H) ppm

The compounds **CKR49-28 and CKR49-31** were synthesized in a similar way.

***N*-(6-Fluorobenzothiazol-2-yl)-3-(6-methoxypyridin-3-yl)benzamide (CKR49-31).**

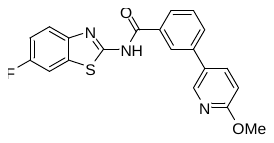

^1^H NMR (600 MHz, CDCl_3_): *δ* 10.12 (bs, 1H), 8.39 (d, *J* = 2.4, 1H), 8.13 (s, 1H), 7.92 (d, *J* = 7.7, 1H), 7.80 – 7.76 (m, 2H), 7.64 – 7.59 (m, 2H), 7.60 (dd, *J* = 8.2, 2.6, 1H), 7.14 (td, *J* = 8.9, 2.3, 1H), 6.84 (d, *J* = 8.5, 1H), 4.00 (s, 3H) ppm

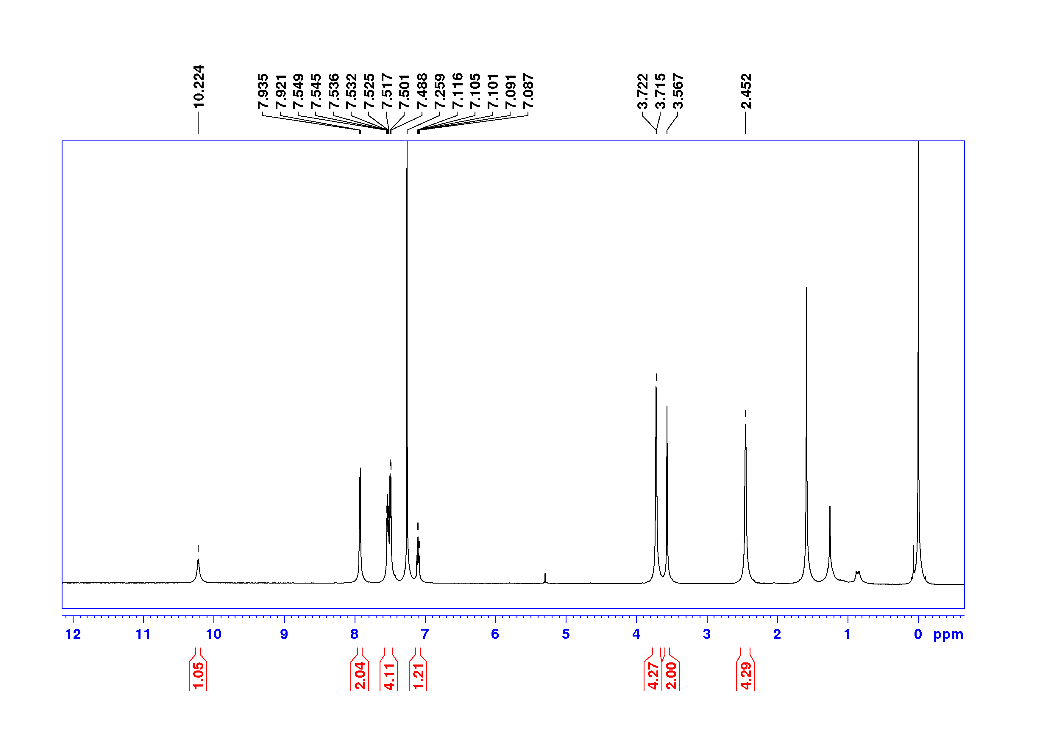

**Figure**. ^1^H NMR of CKR49

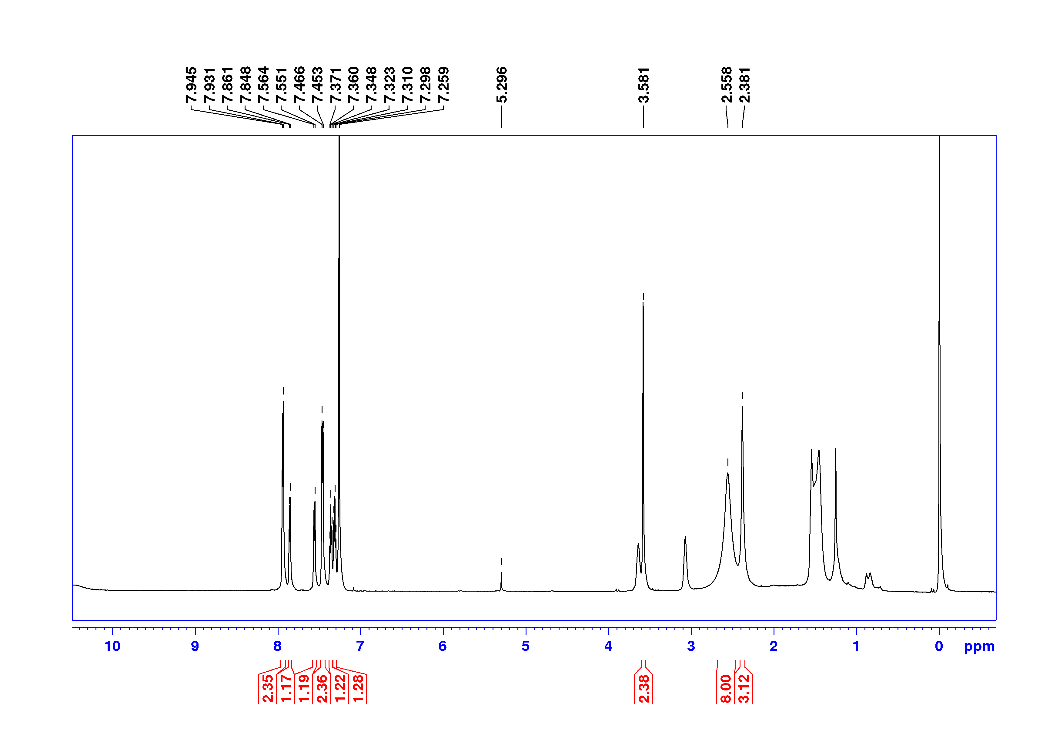

**Figure**. ^1^H NMR of CKR49-1

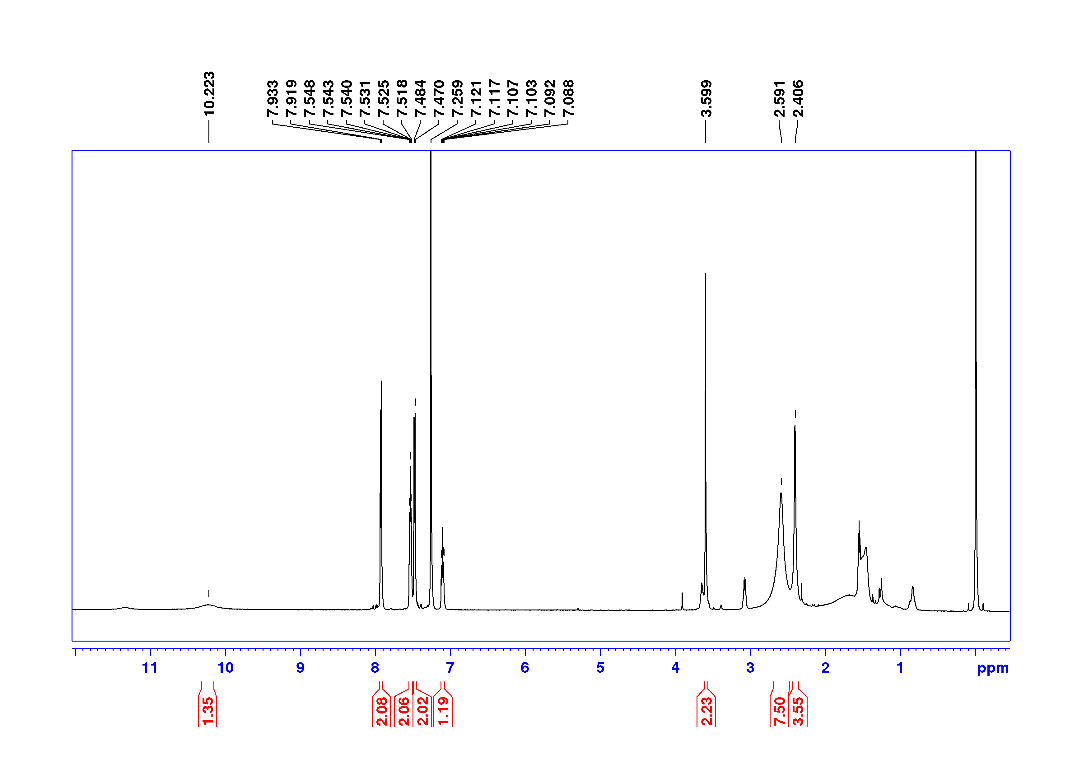

**Figure**. ^1^H NMR of CKR49-2

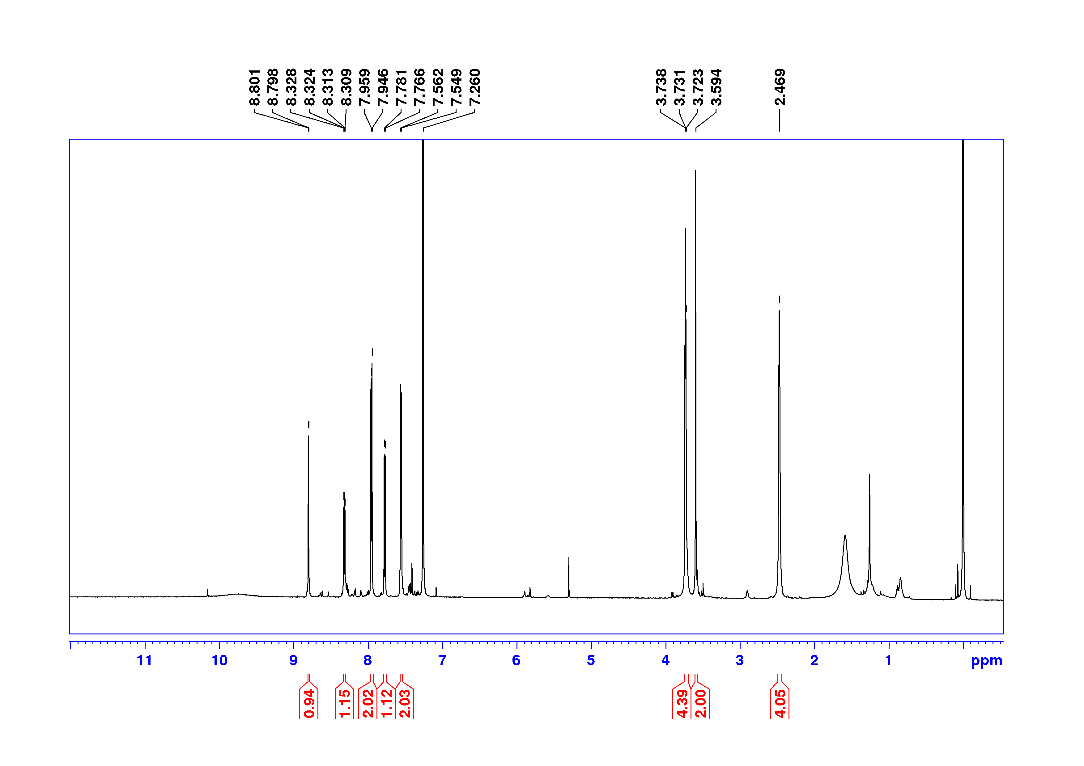

**Figure**. ^1^H NMR of CKR49-6

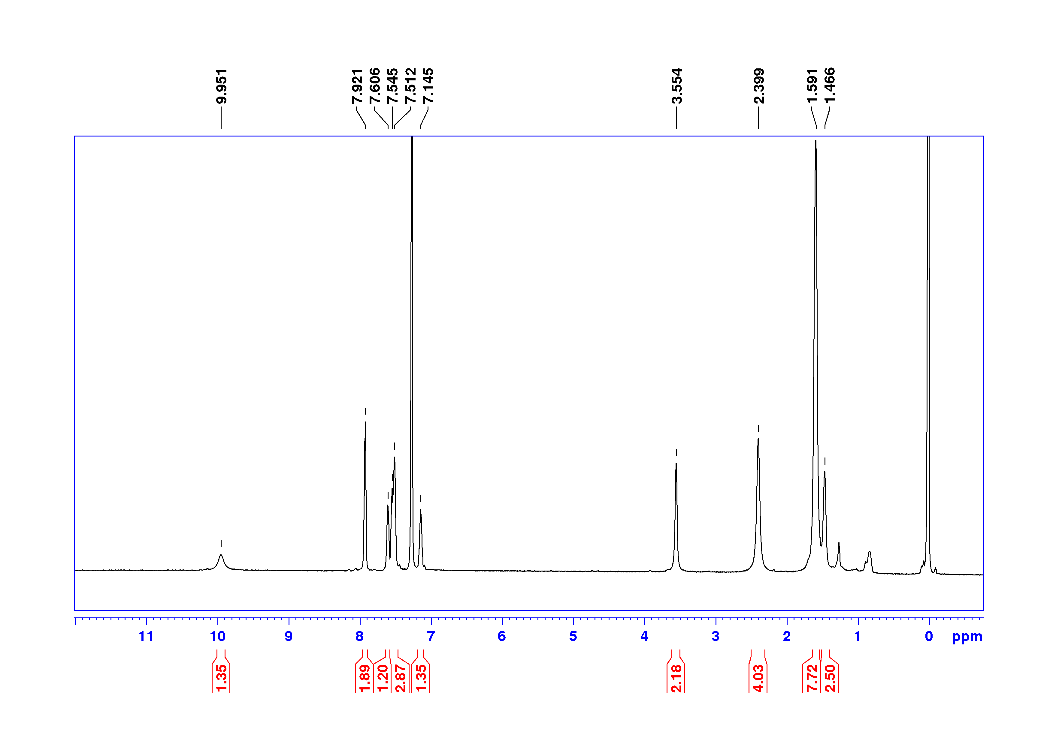

**Figure**. ^1^H NMR of CKR49-9

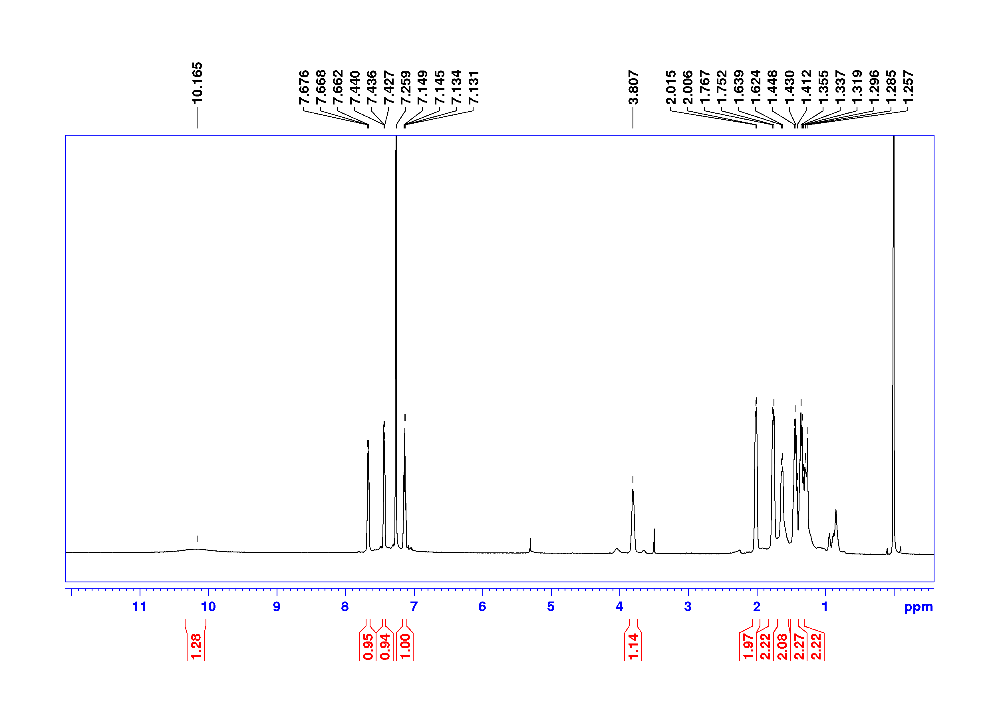

**Figure**. ^1^H NMR of CKR49-13

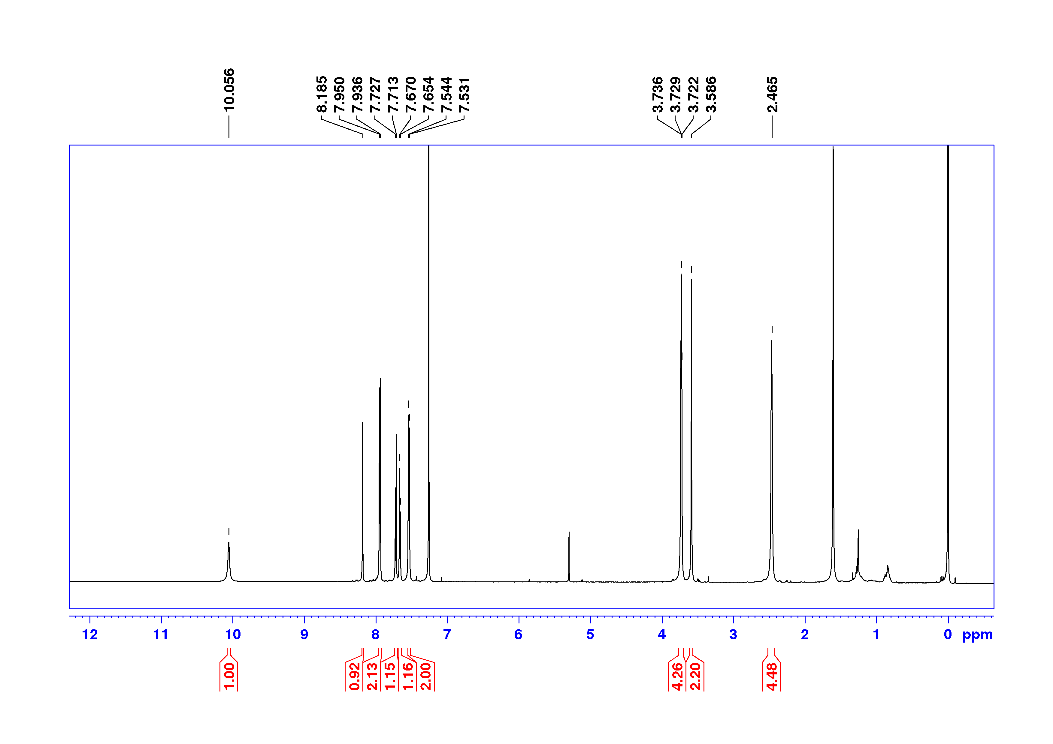

**Figure**. ^1^H NMR of CKR49-15

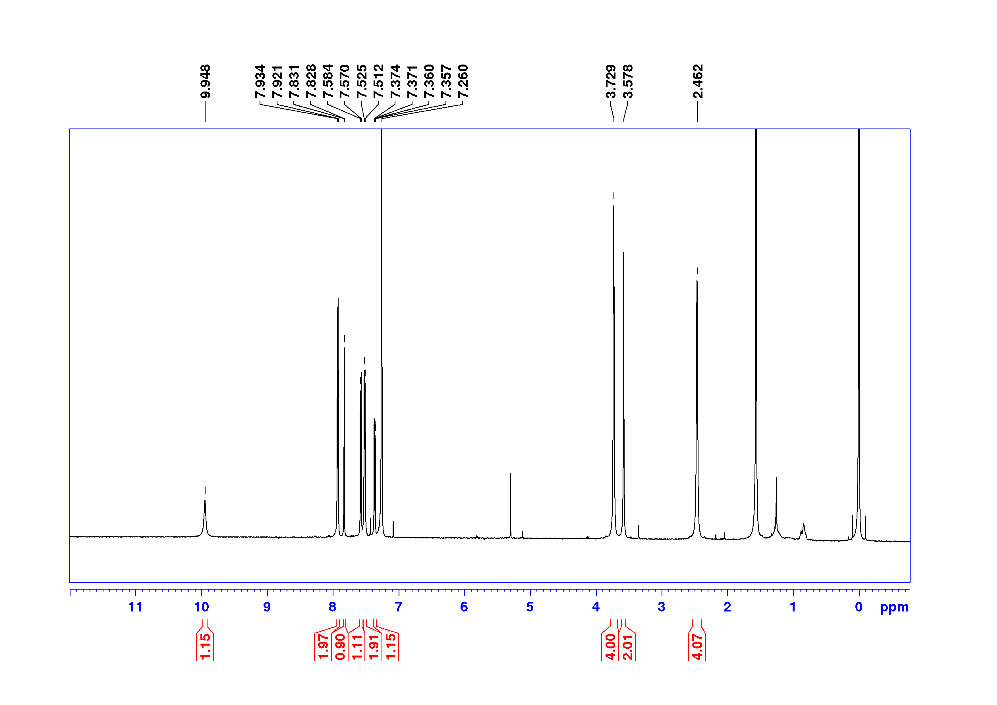

**Figure**. ^1^H NMR of CKR49-16

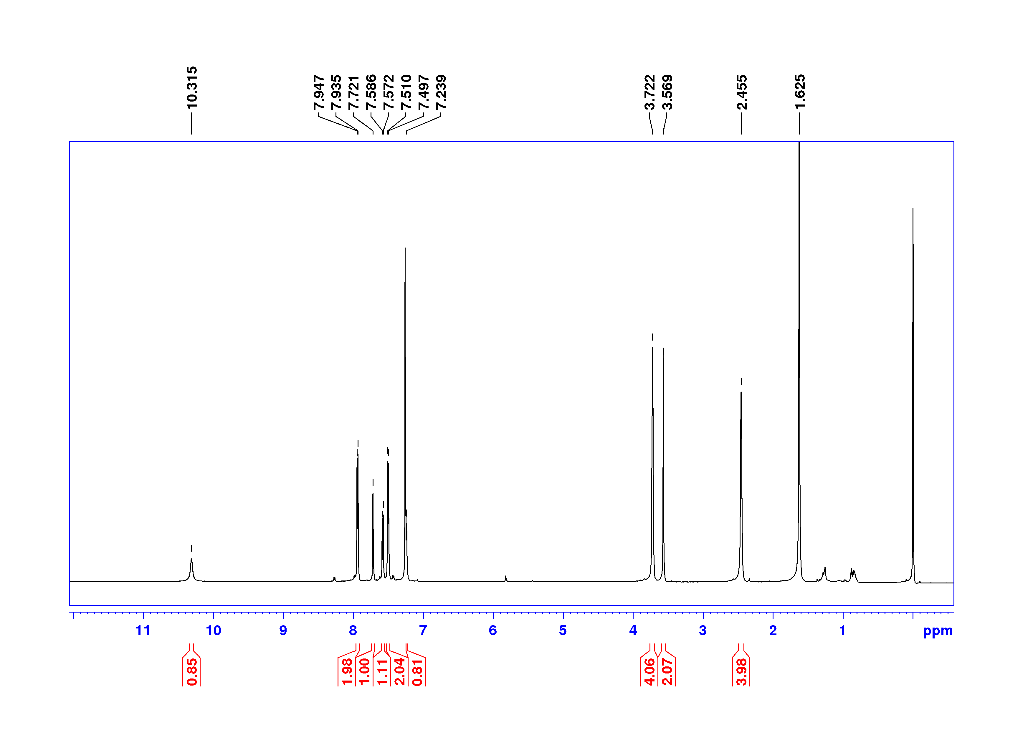

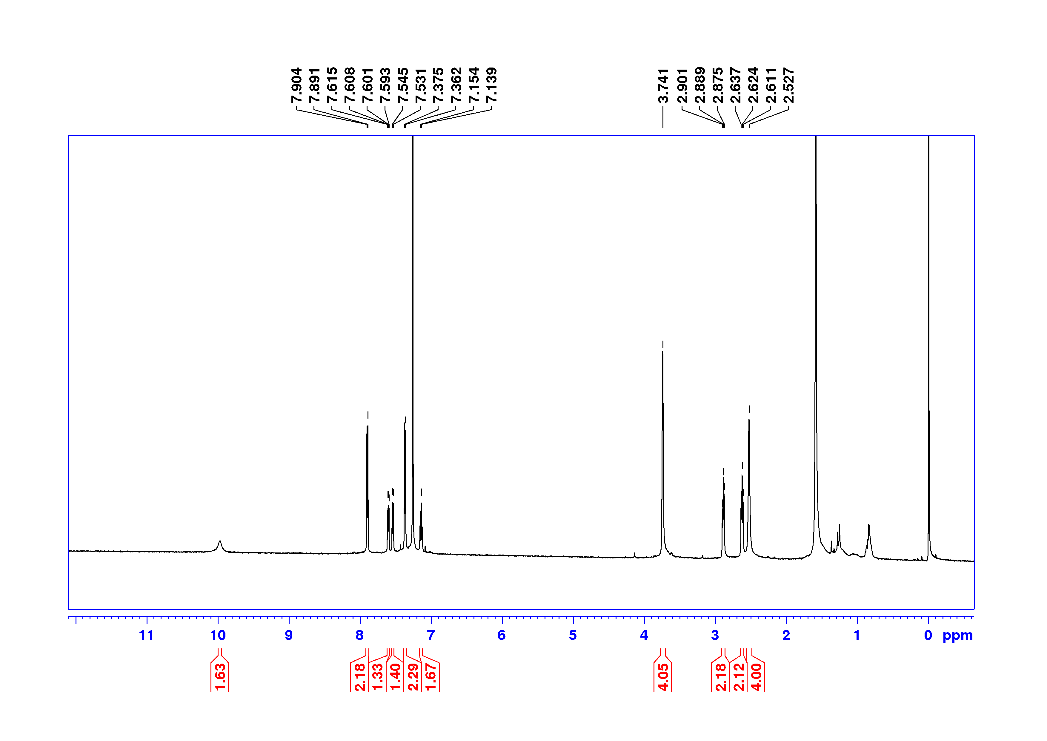
**Figure**. ^1^H NMR of CKR49-20

**Figure**. ^1^H NMR of CKR49-23

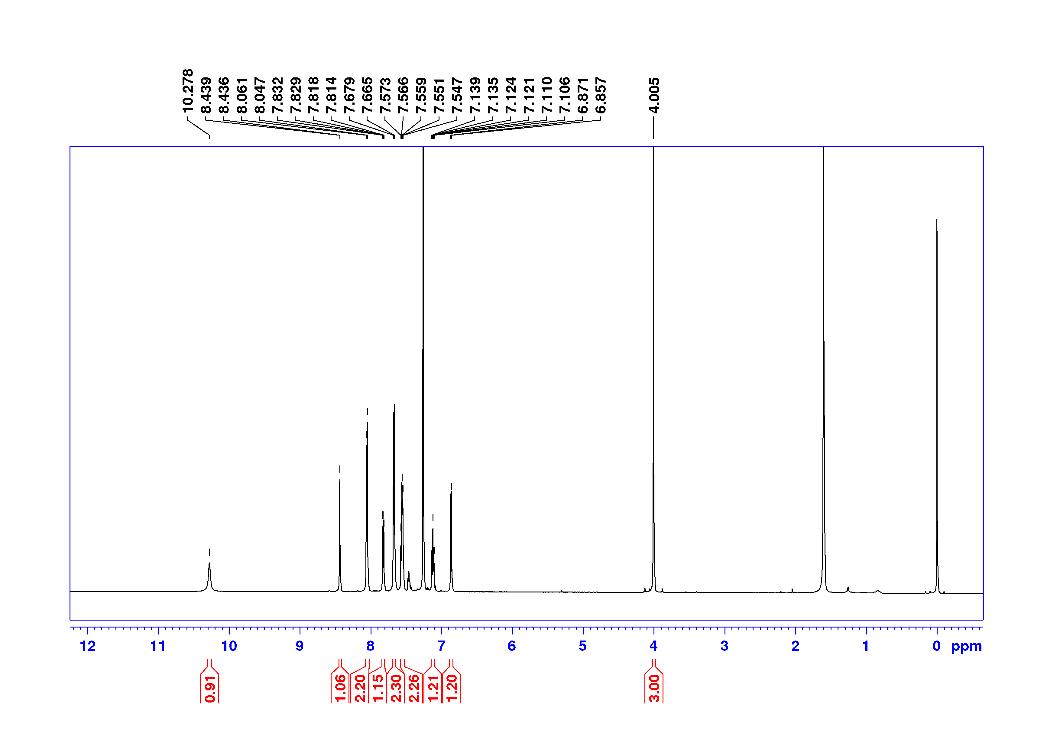

**Figure**. ^1^H NMR of CKR49-30

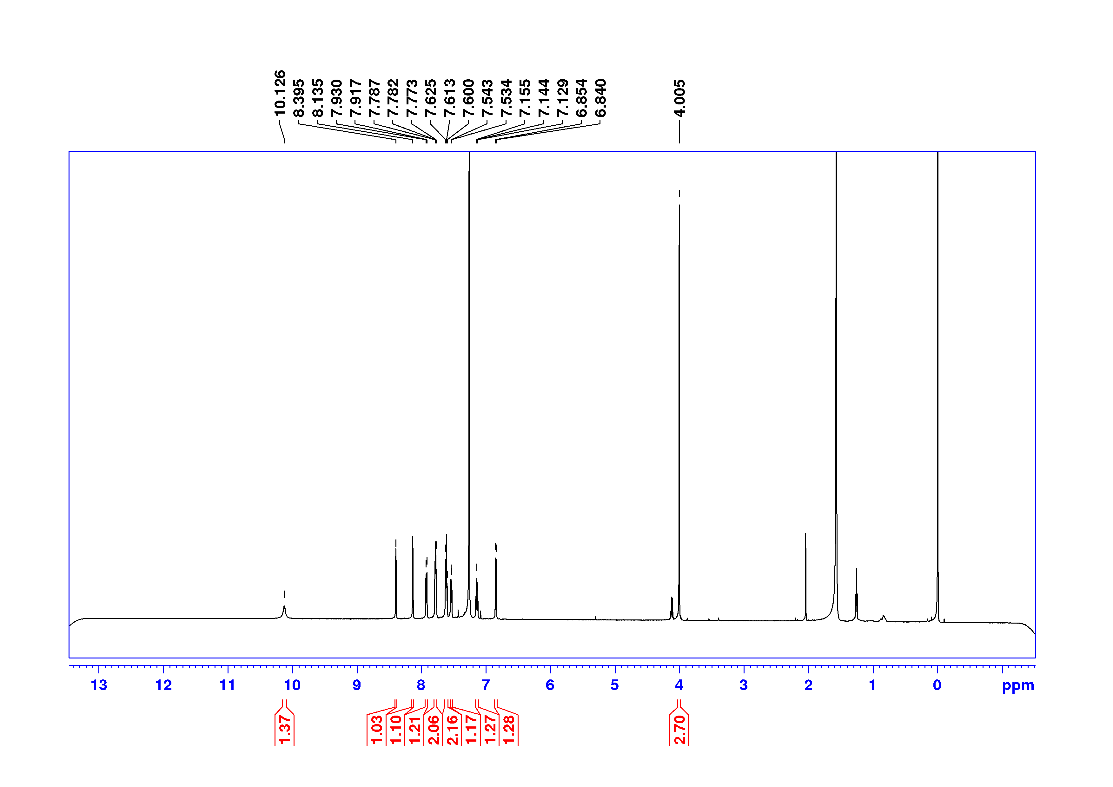

**Figure**. ^1^H NMR of CKR49-31

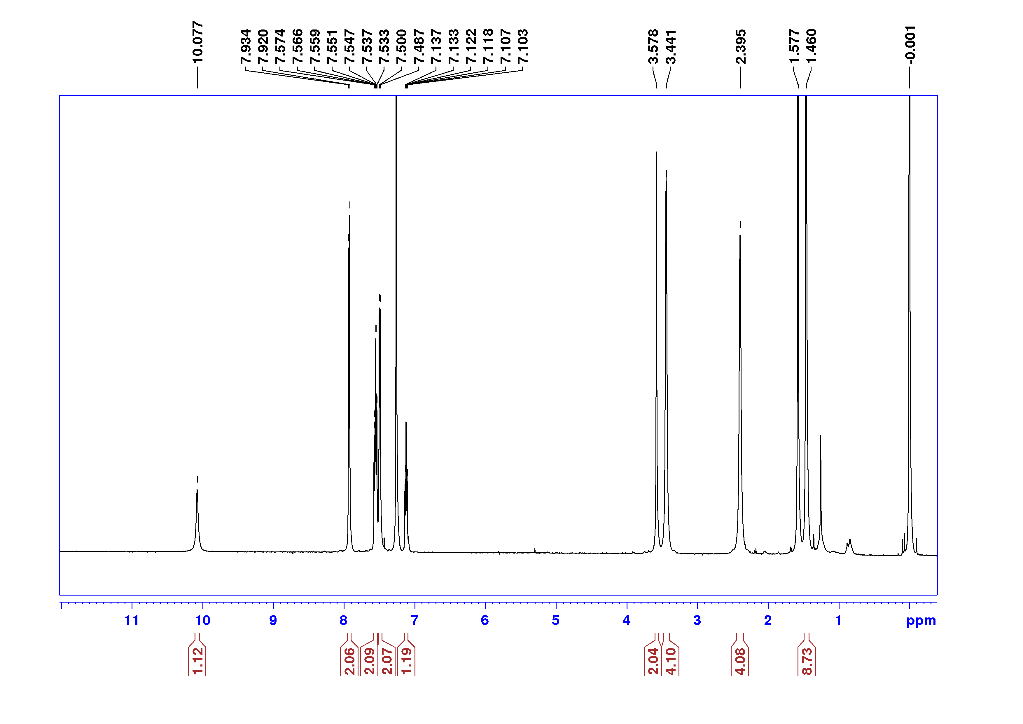

**Figure**. ^1^H NMR of 2a

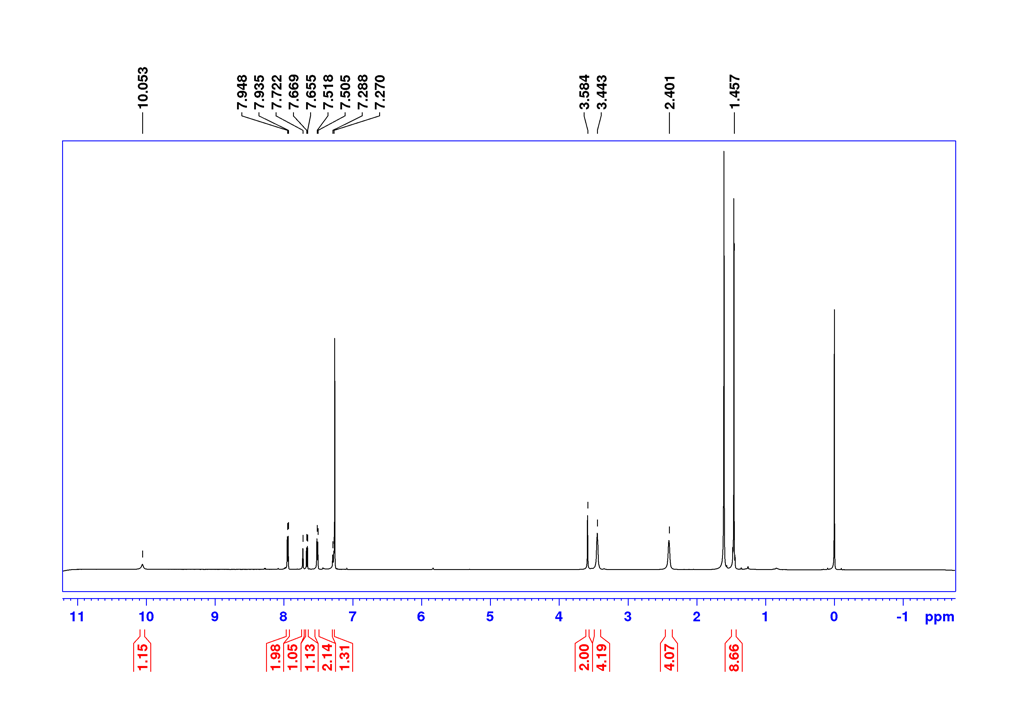

**Figure**. ^1^H NMR of 2e

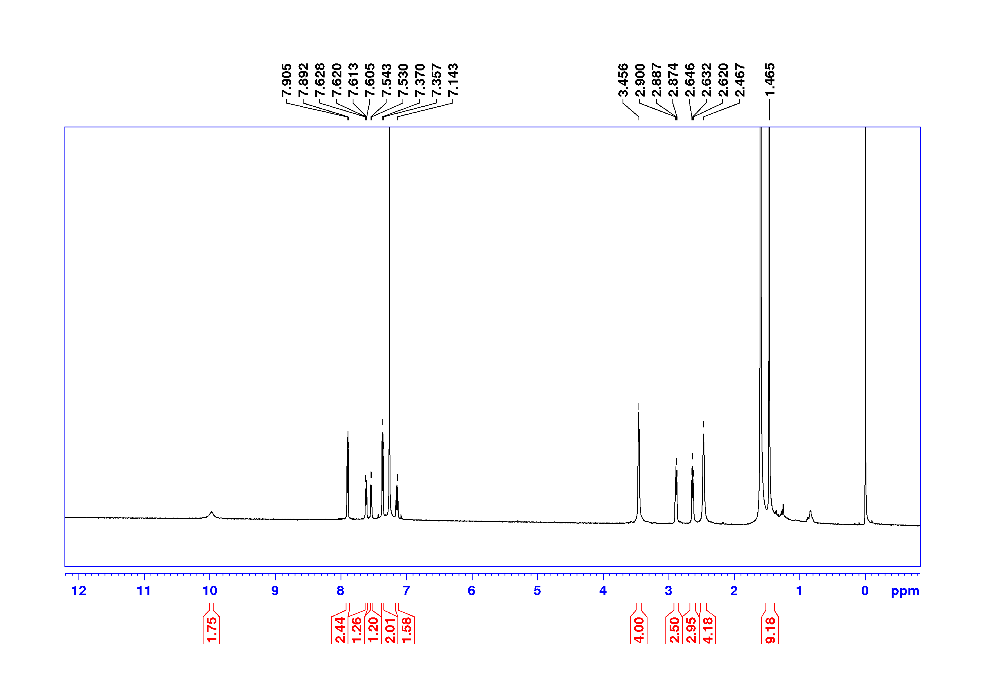

**Figure**. ^1^H NMR of 2g

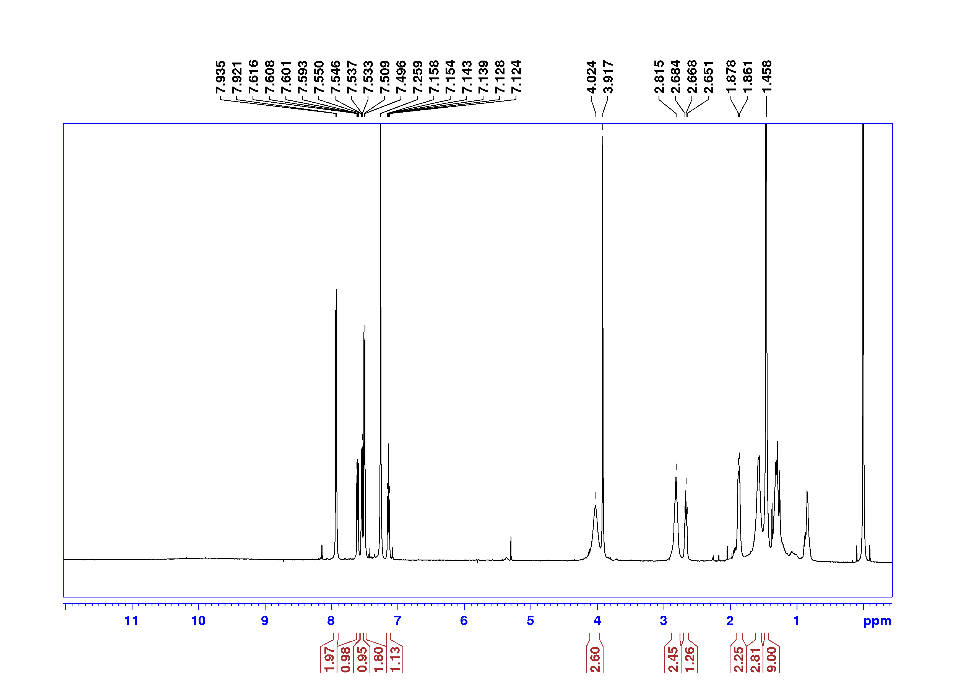

**Figure**. ^1^H NMR of 4a

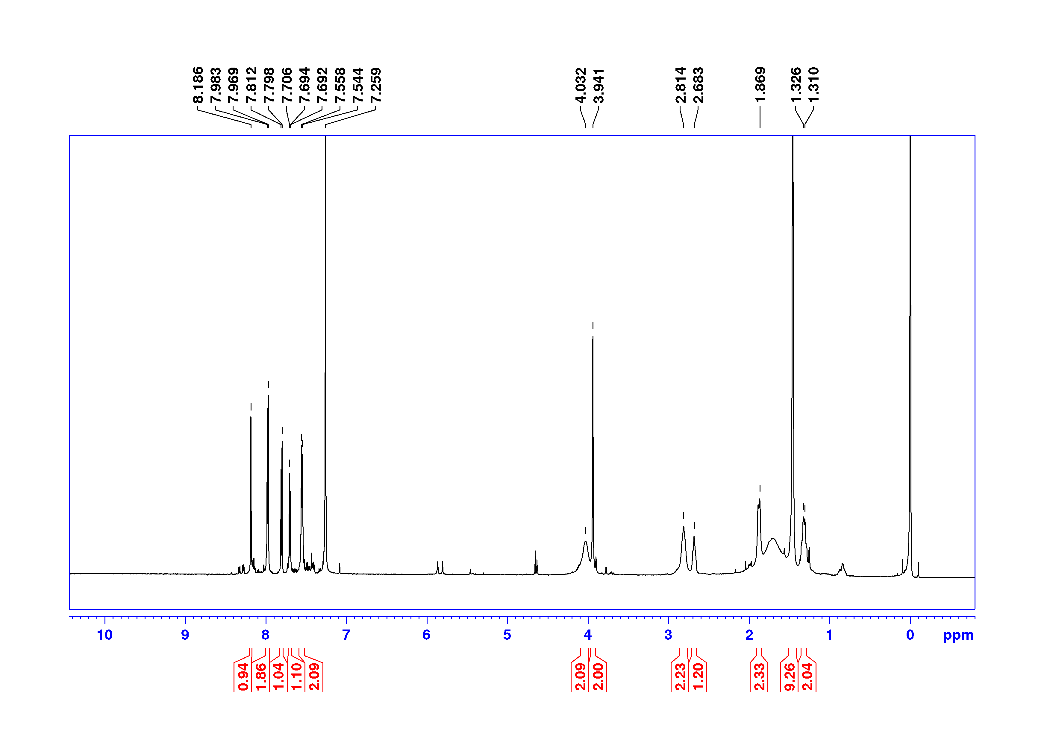

**Figure**. ^1^H NMR of 4d

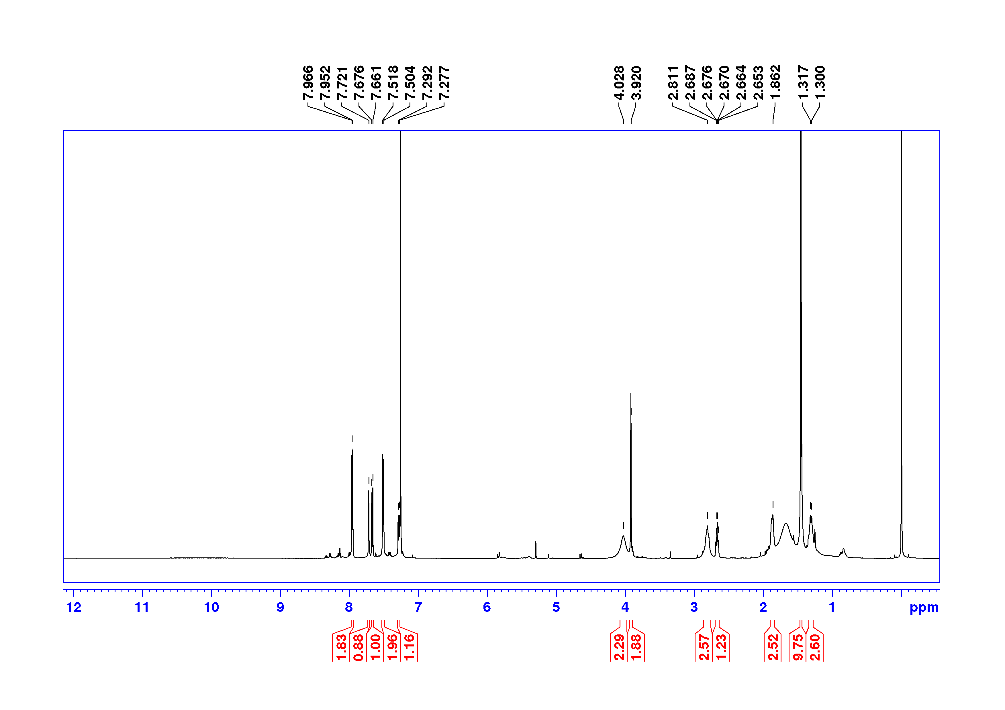

**Figure**. ^1^H NMR of 4e

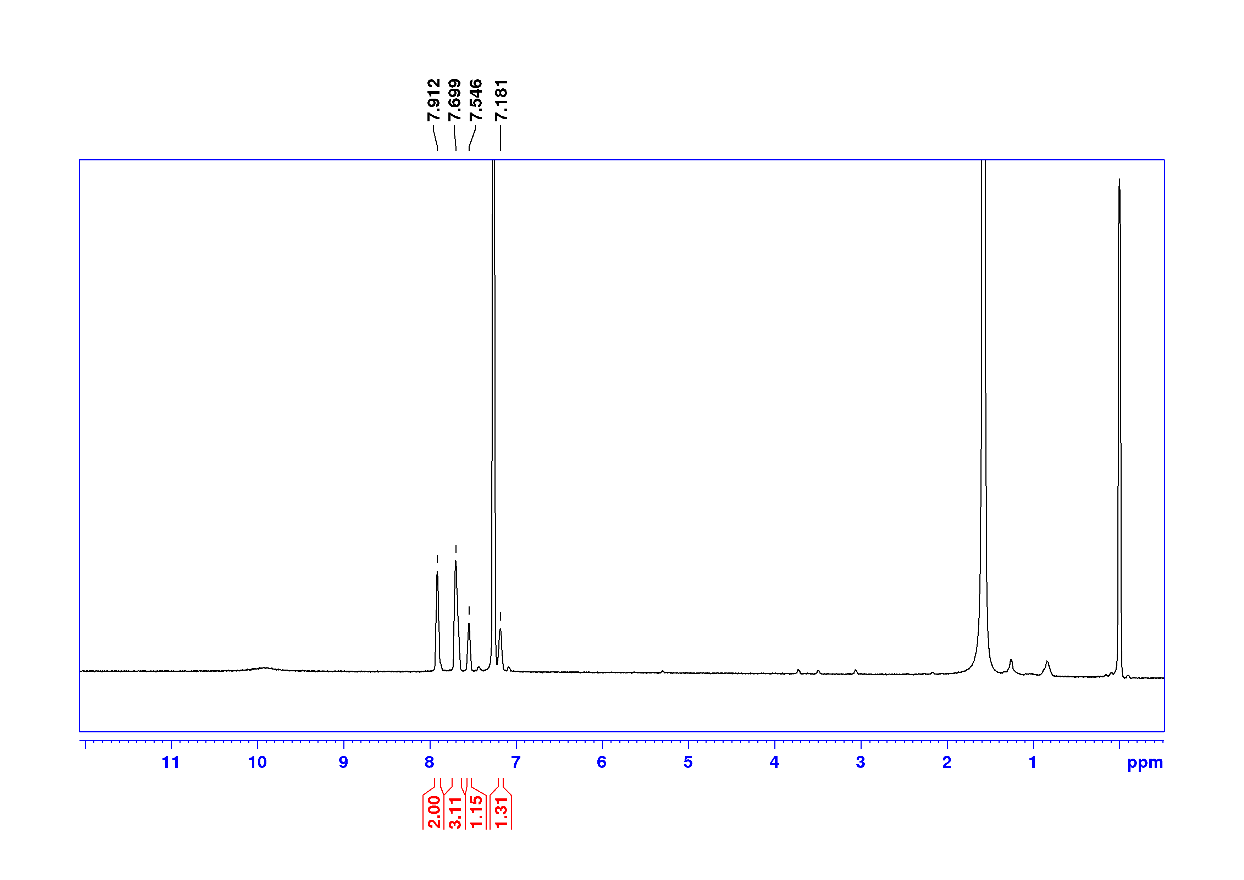

**Figure**. ^1^H NMR of 5a

LCMS analysis Conditions: The mobile phase consisted of 0.1% formic acid in deionized water (A) and acetonitrile (B). The samples were injected onto a RP chromatography column (Targa C18, 5 μm, 50 x 2.1 mm, 120 A^°^) in ESI positive ion mode, and gradient elution was as follows: 1% B hold for 0.5 minute; 1%-95% B for 5 minutes; 95% B continued until 8 minutes and set the post time for 3 minutes to equilibrate; at a flow rate of 0.4 mL/min and the column temperature at 40 °C.

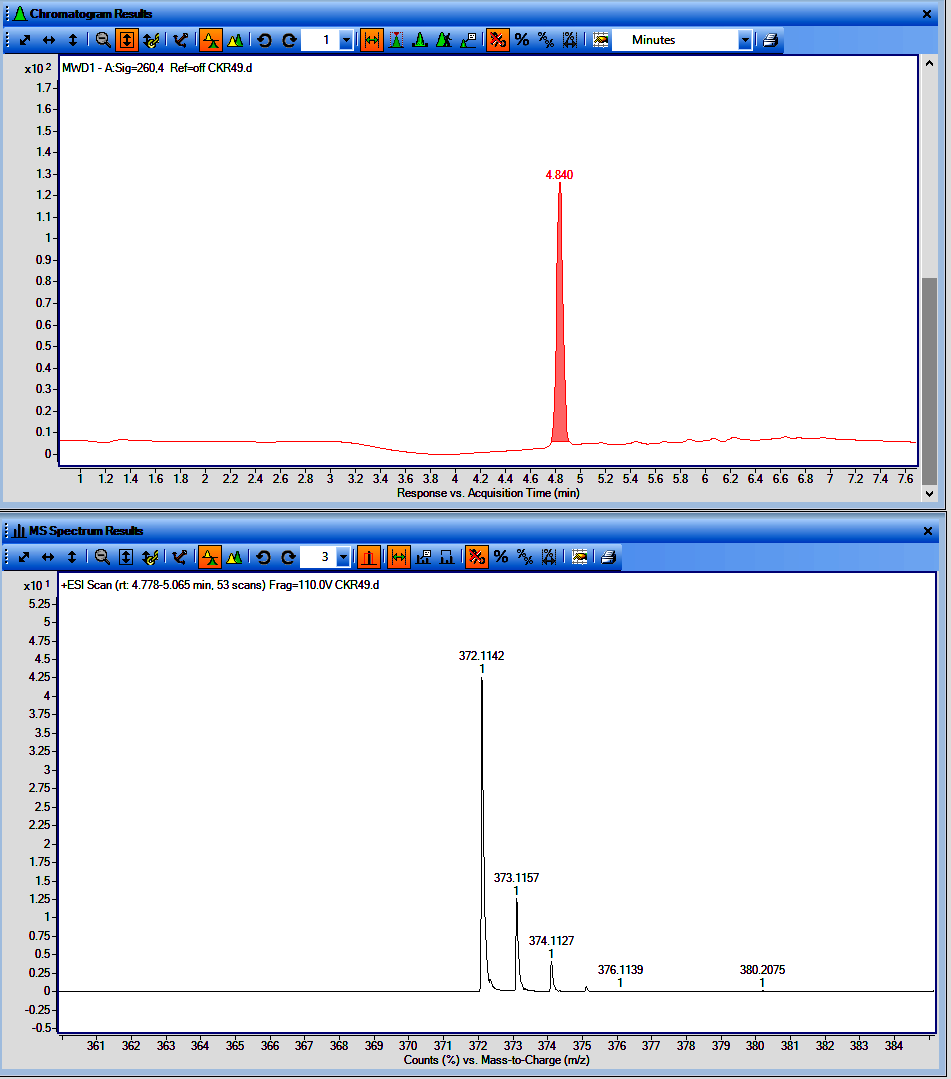

**Figure**. LCMS chromatogram of CKR49 (UV with t_R_ and MS spectrum)

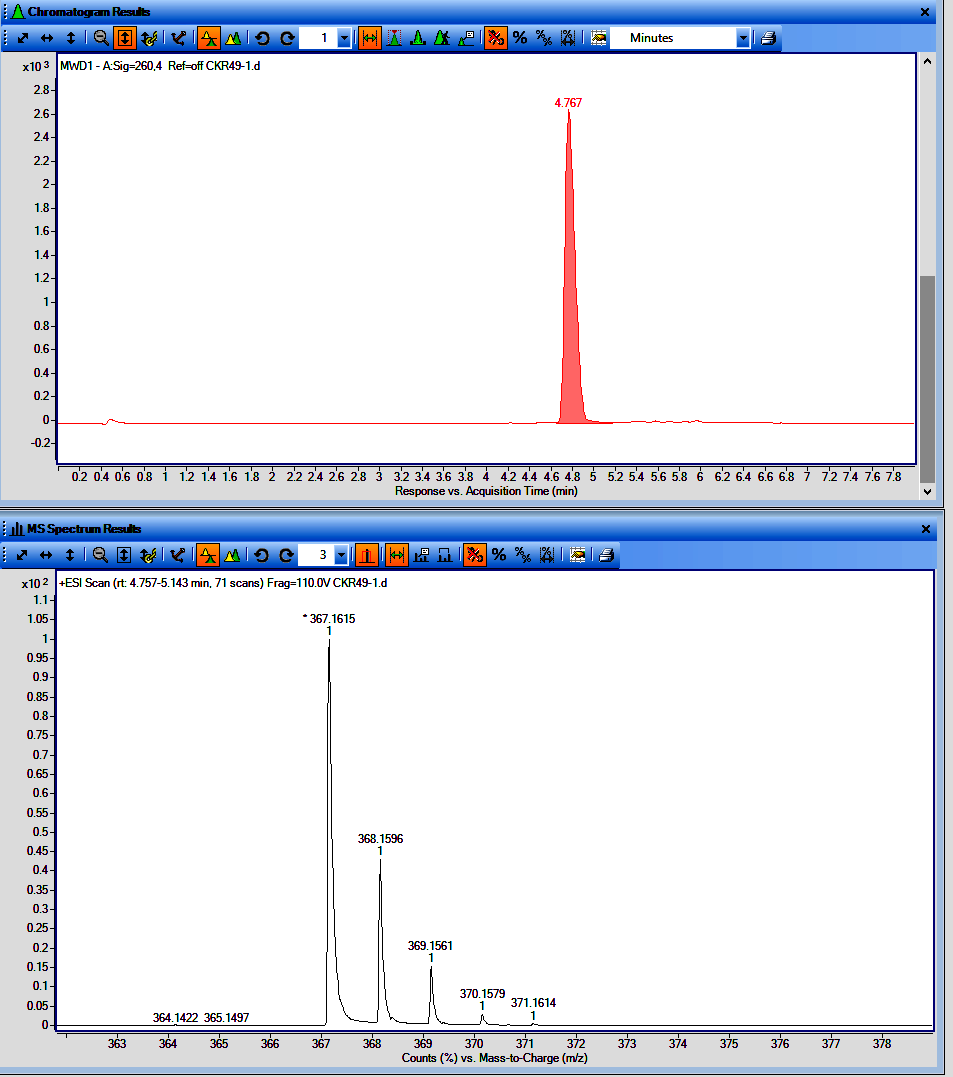

**Figure**. LCMS chromatogram of CKR49-1 (UV with t_R_ and MS spectrum)

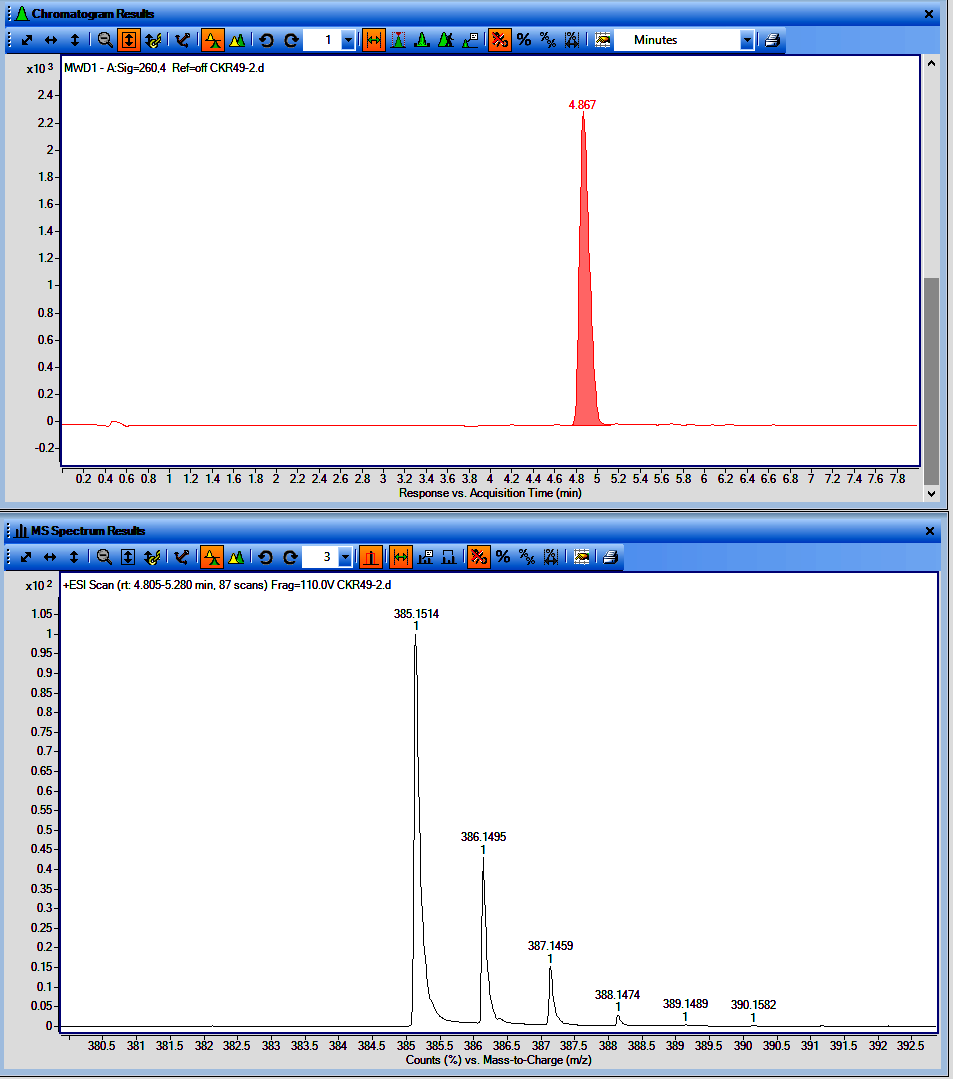

**Figure**. LCMS chromatogram of CKR49-2 (UV with t_R_ and MS spectrum)

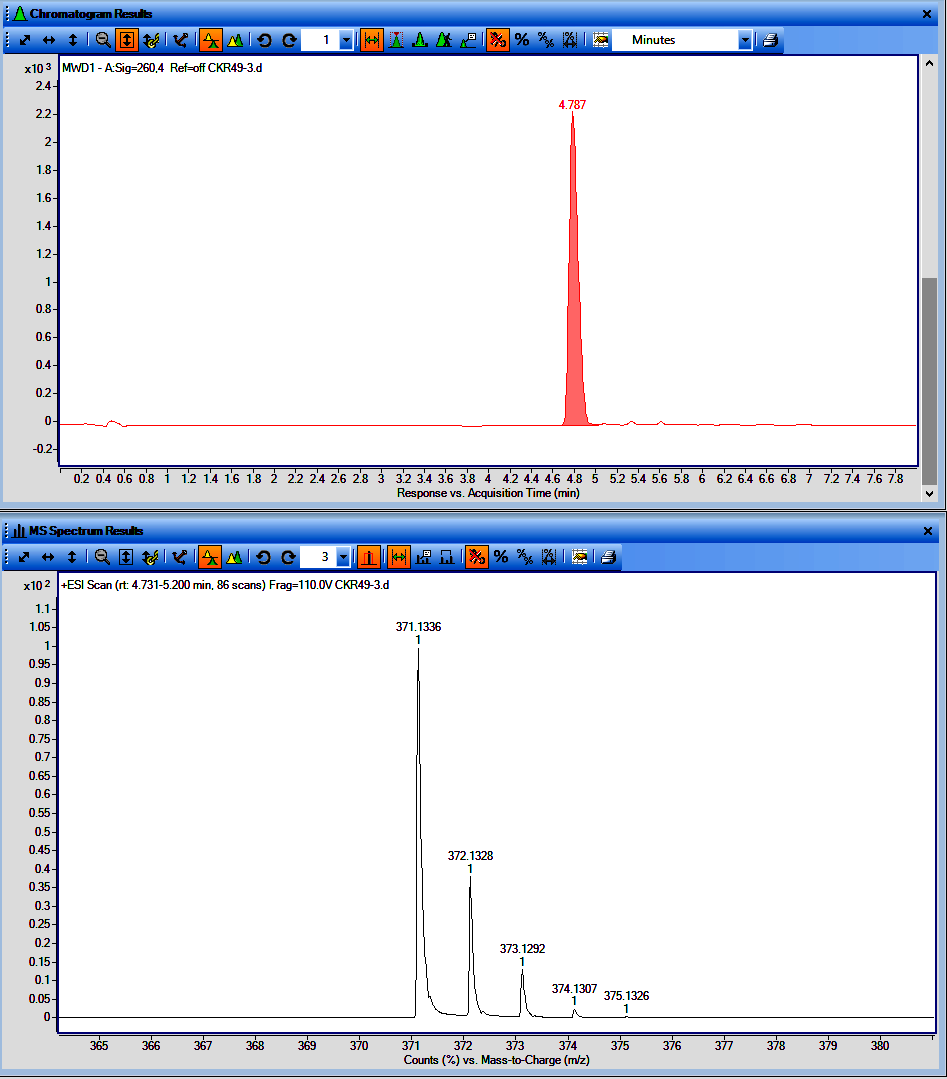

**Figure**. LCMS chromatogram of CKR49-3 (UV with t_R_ and MS spectrum)

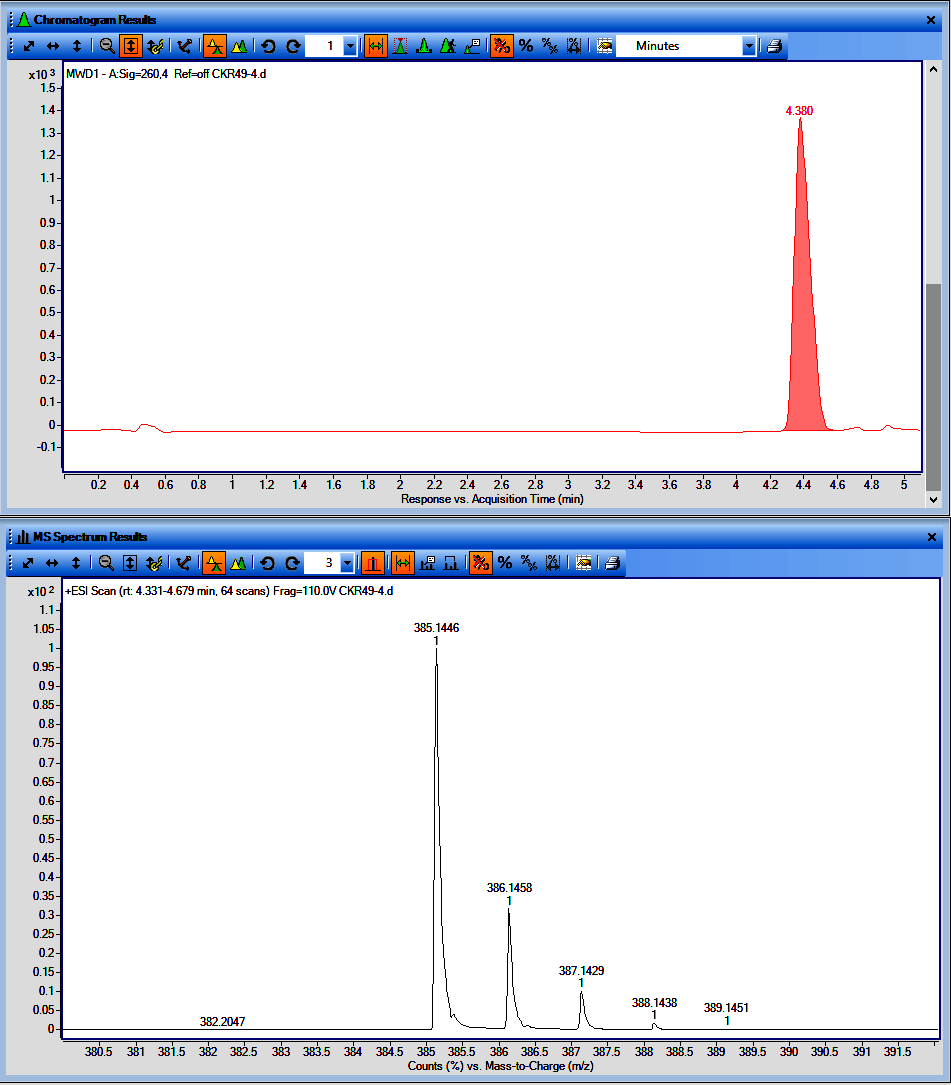

**Figure**. LCMS chromatogram of CKR49-4 (UV with t_R_ and MS spectrum)

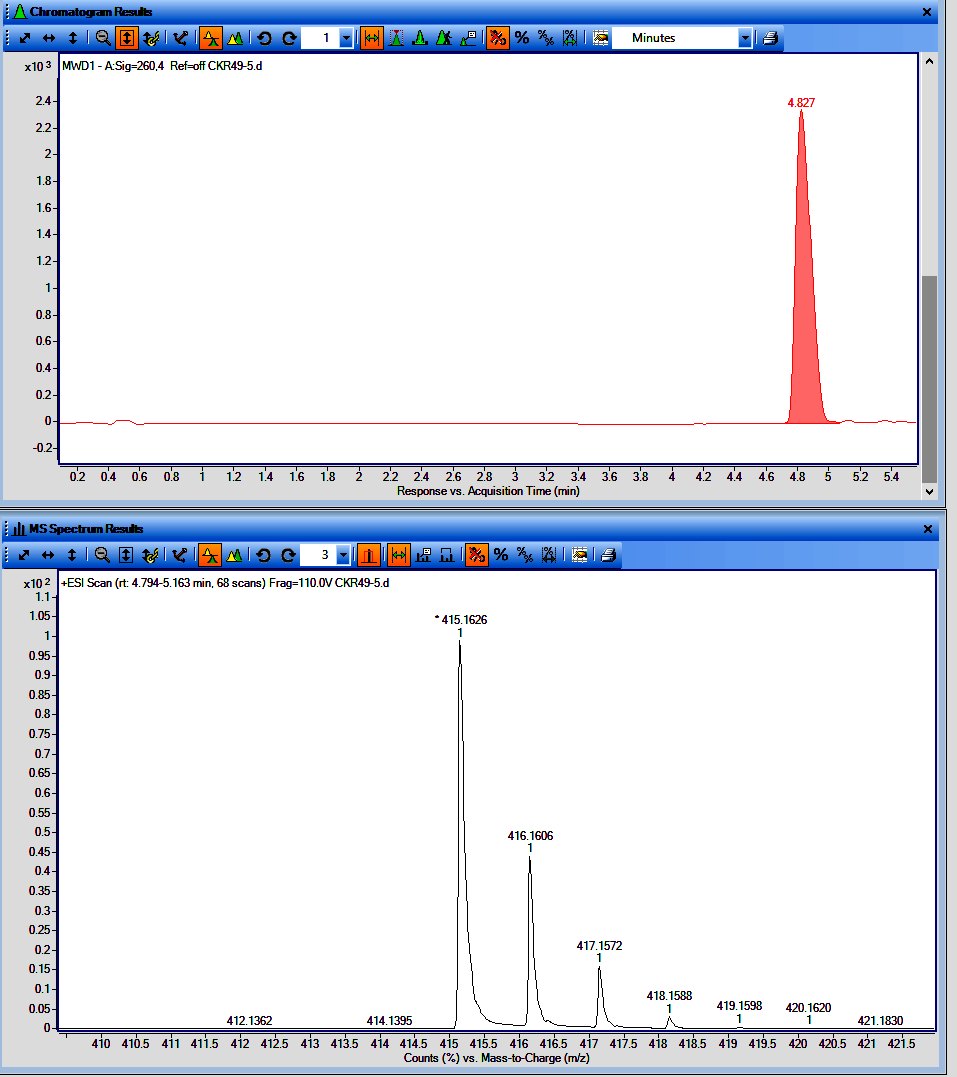

**Figure**. LCMS chromatogram of CKR49-5 (UV with t_R_ and MS spectrum)

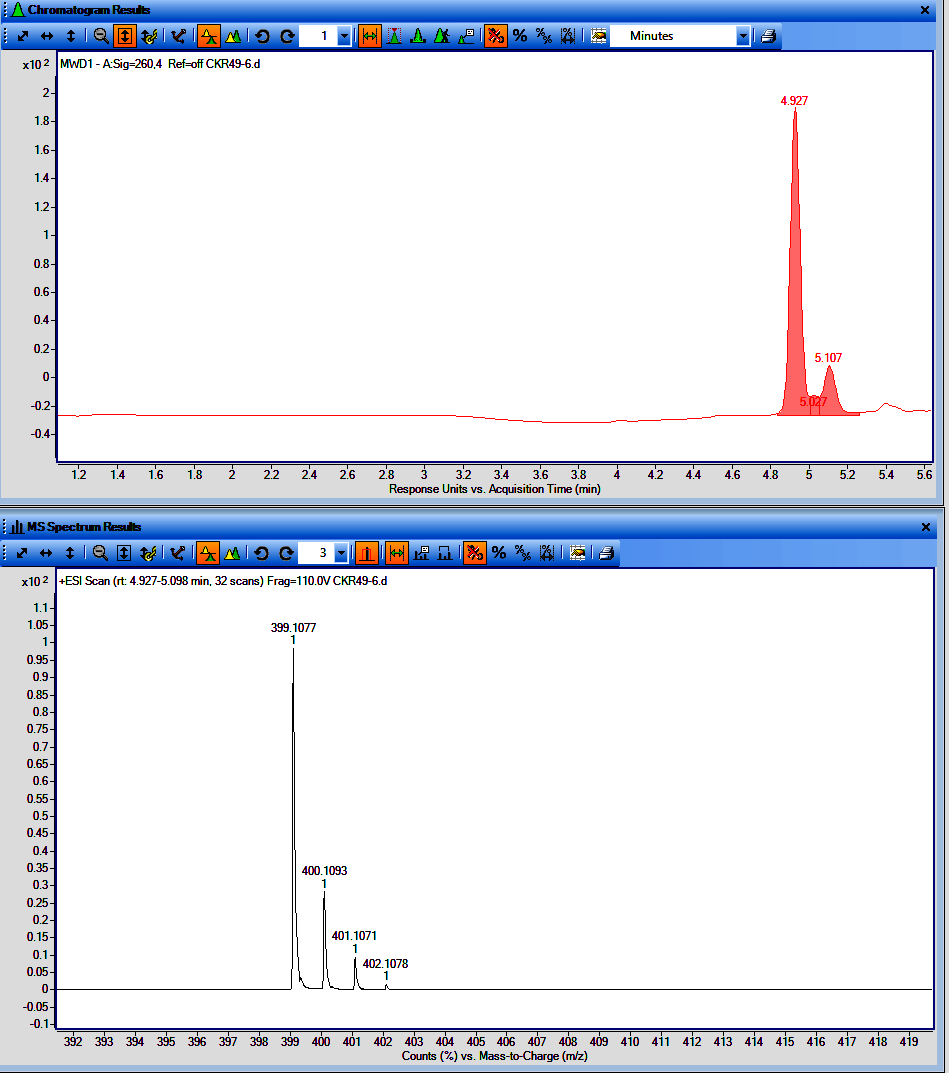

**Figure**. LCMS chromatogram of CKR49-6 (UV with t_R_ and MS spectrum)

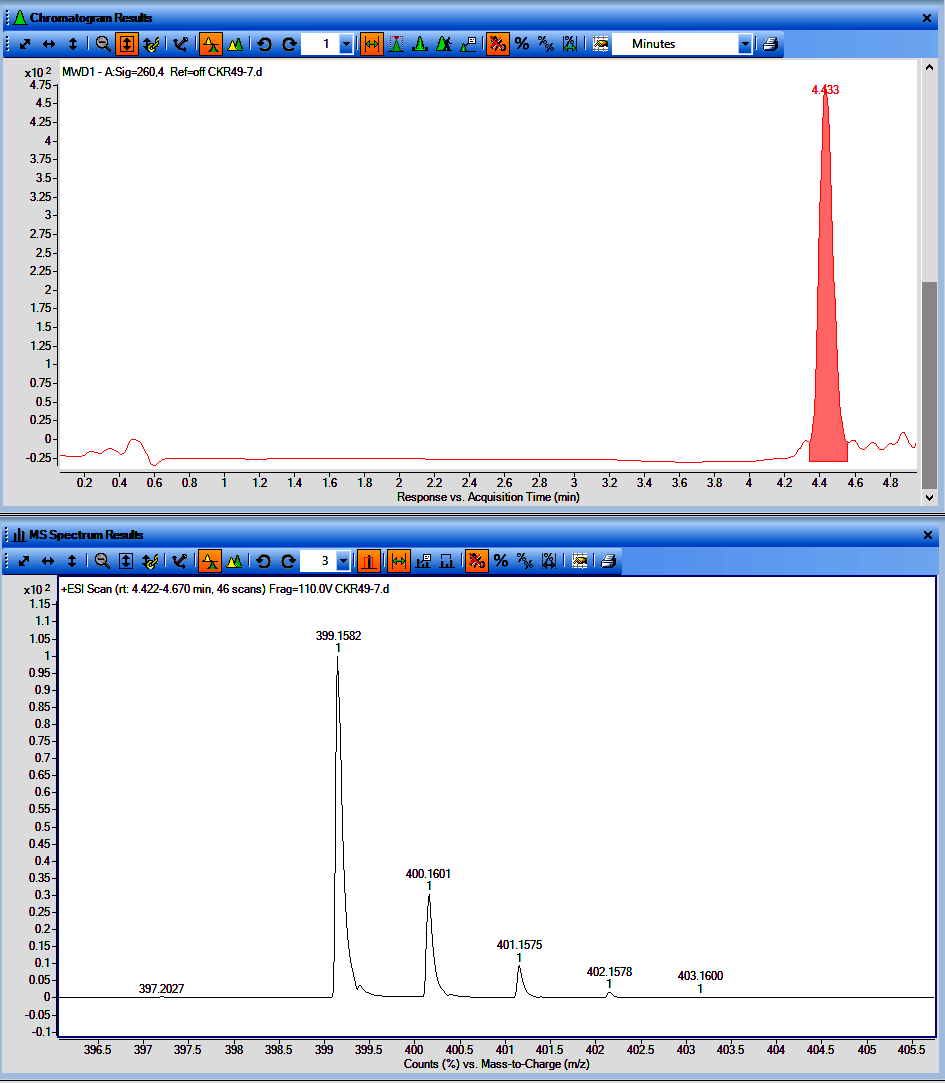

**Figure**. LCMS chromatogram of CKR49-7 (UV with t_R_ and MS spectrum)

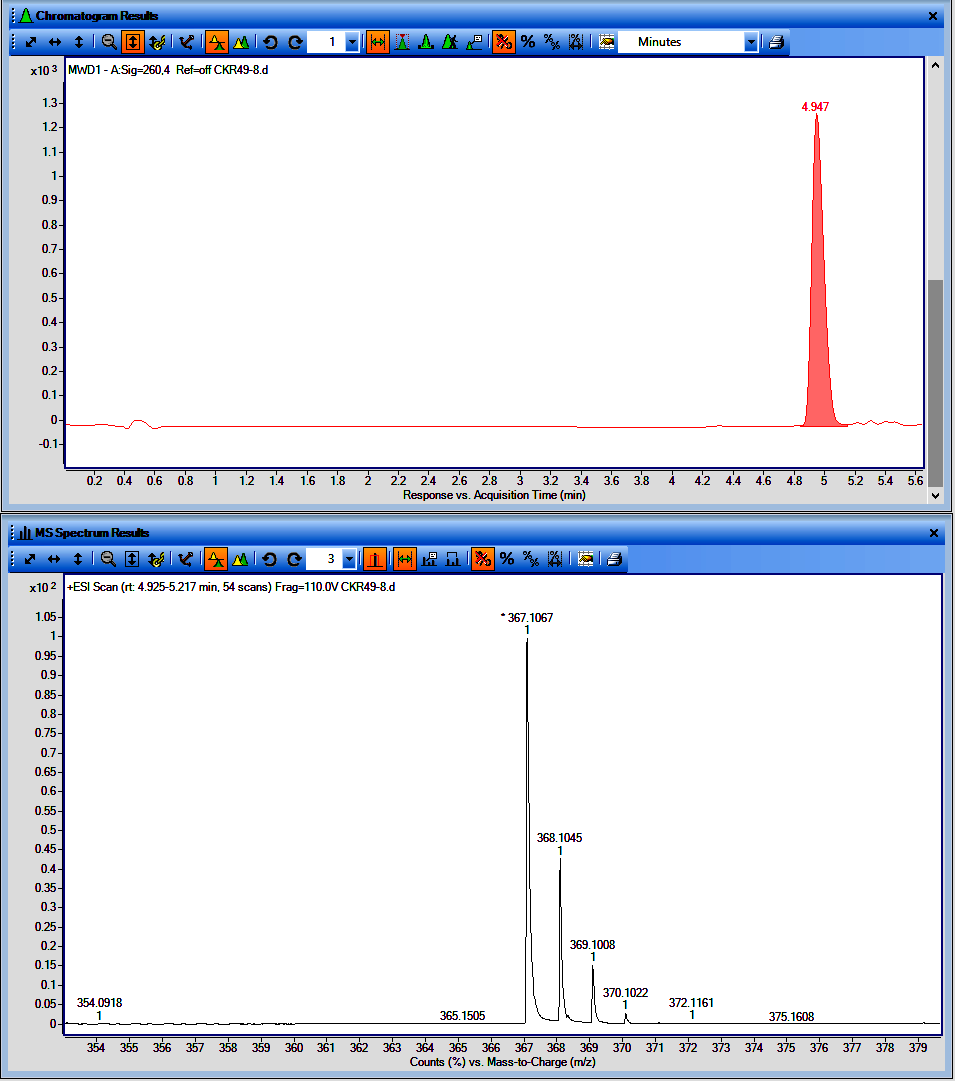

**Figure**. LCMS chromatogram of CKR49-8 (UV with t_R_ and MS spectrum)

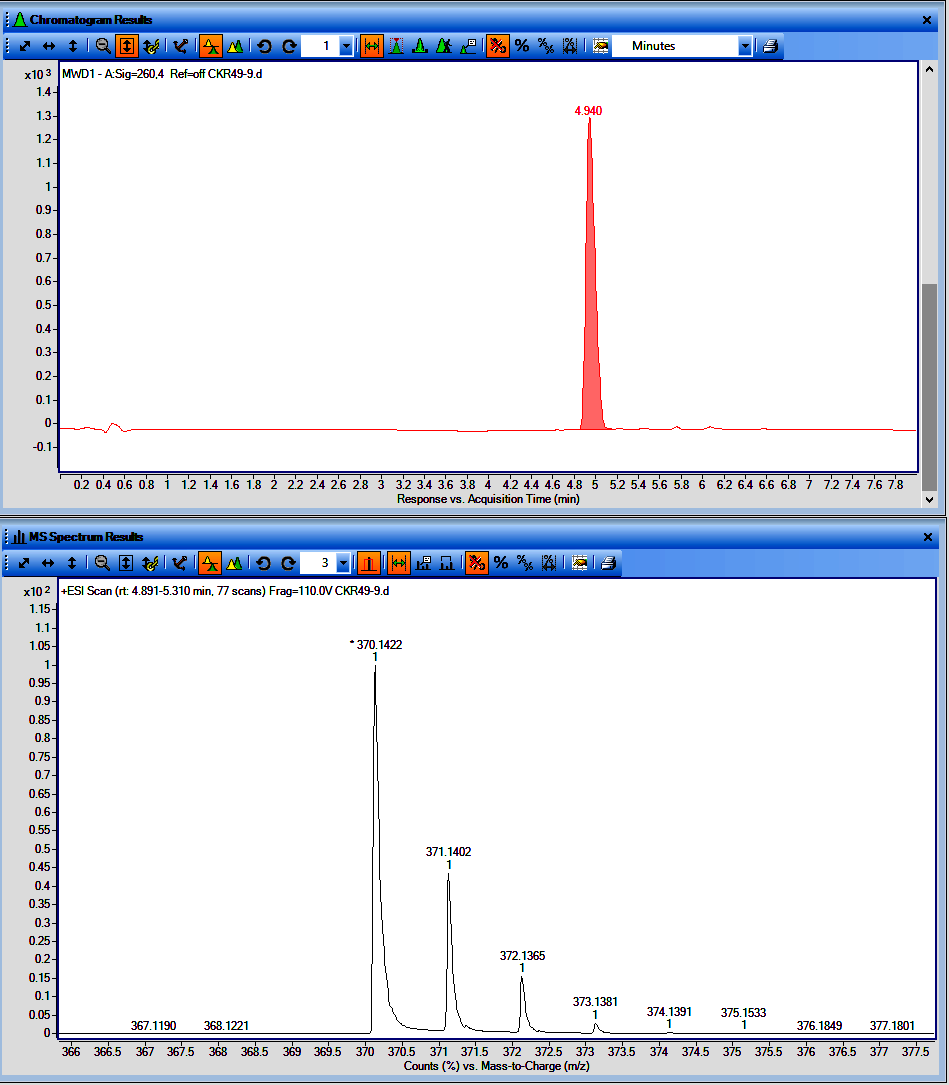

**Figure**. LCMS chromatogram of CKR49-9 (UV with t_R_ and MS spectrum)

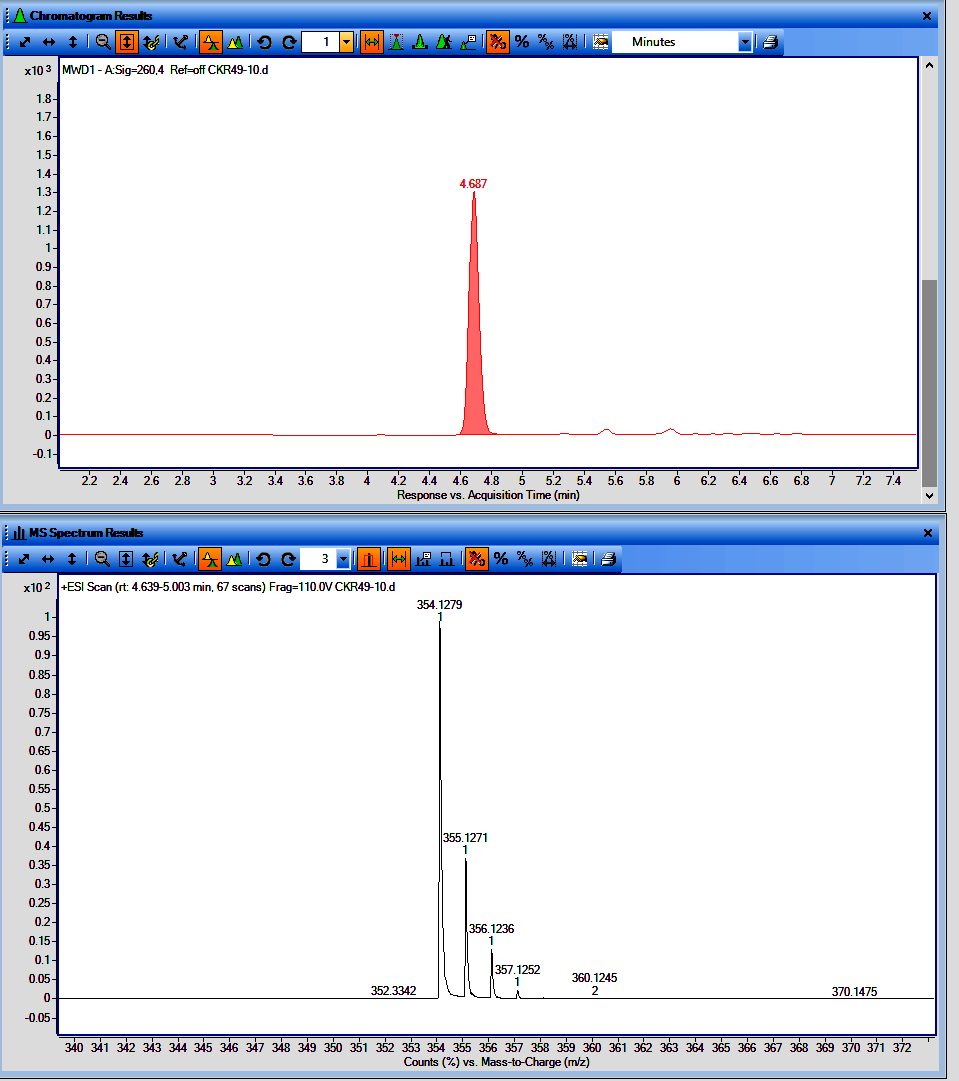

**Figure**. LCMS chromatogram of CKR49-10 (UV with t_R_ and MS spectrum)

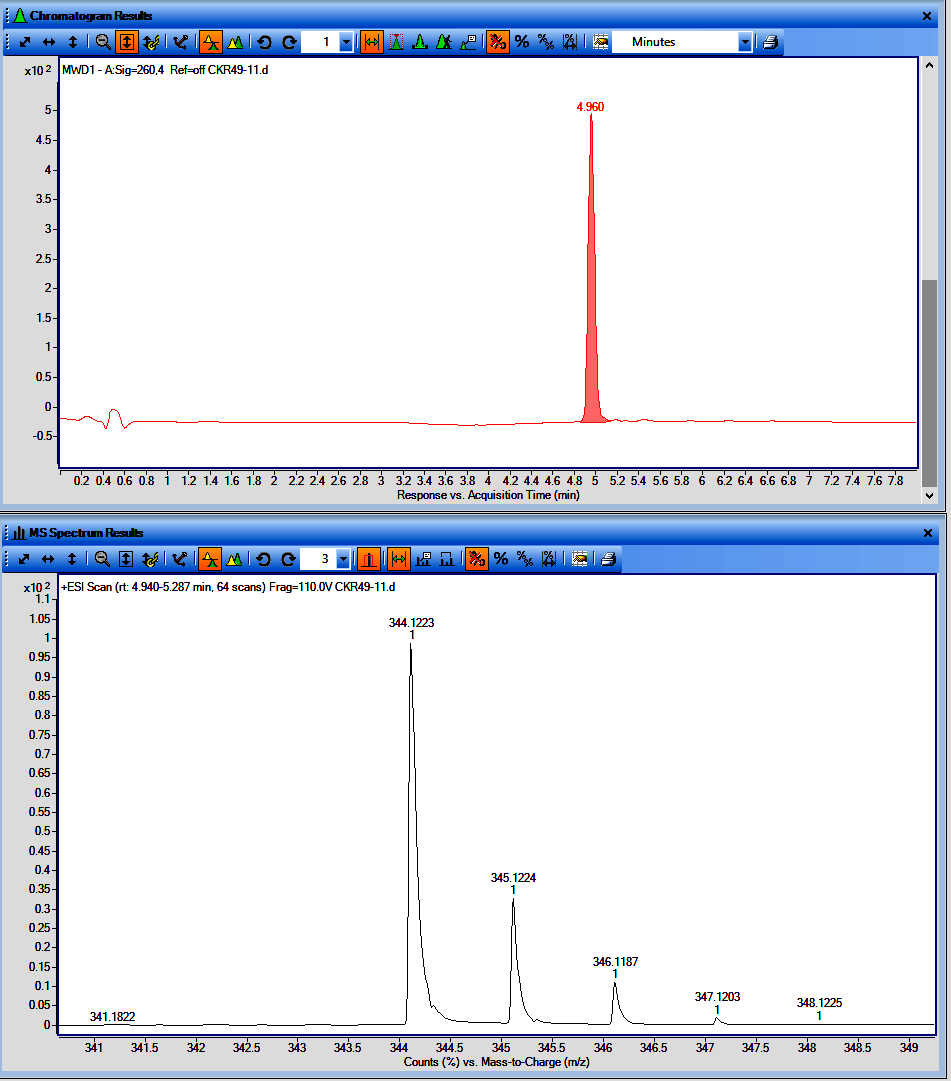

**Figure**. LCMS chromatogram of CKR49-11 (UV with t_R_ and MS spectrum)

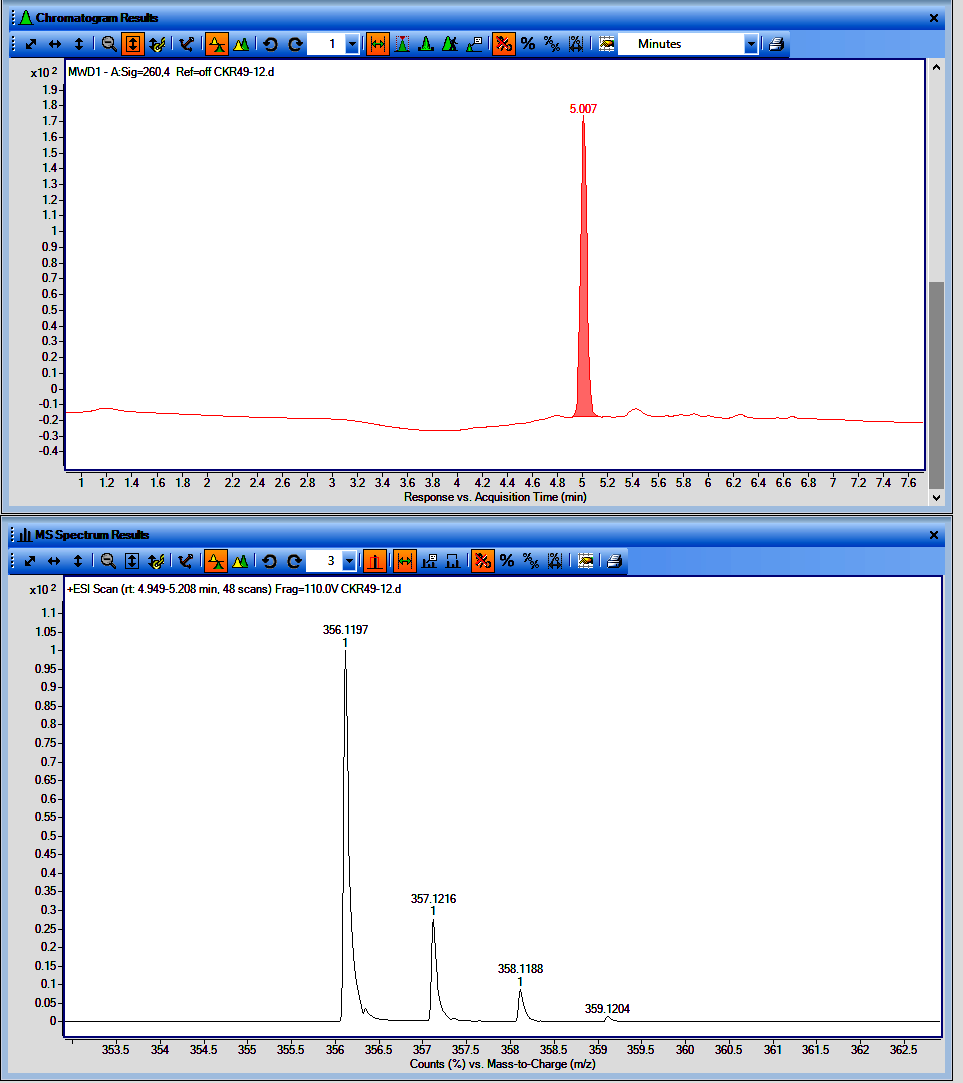

**Figure**. LCMS chromatogram of CKR49-12 (UV with t_R_ and MS spectrum)

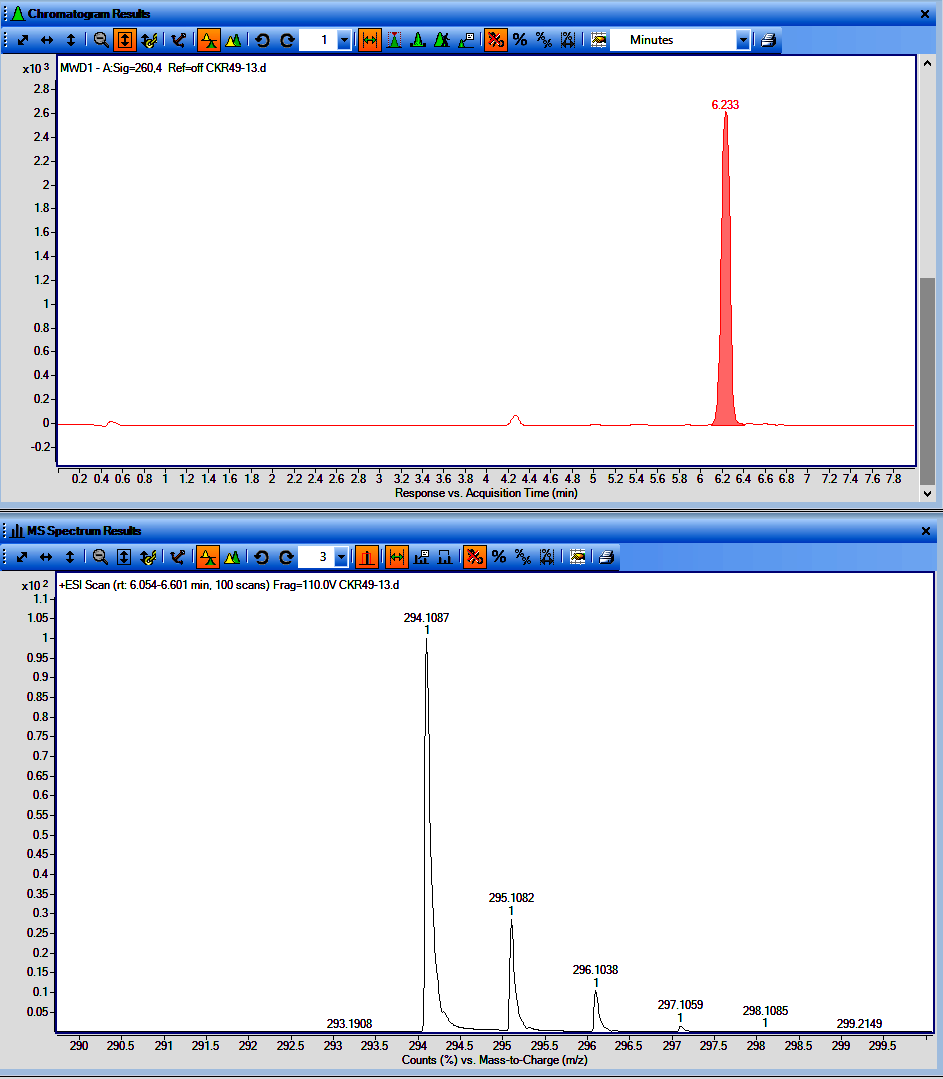

**Figure**. LCMS chromatogram of CKR49-13 (UV with t_R_ and MS spectrum)

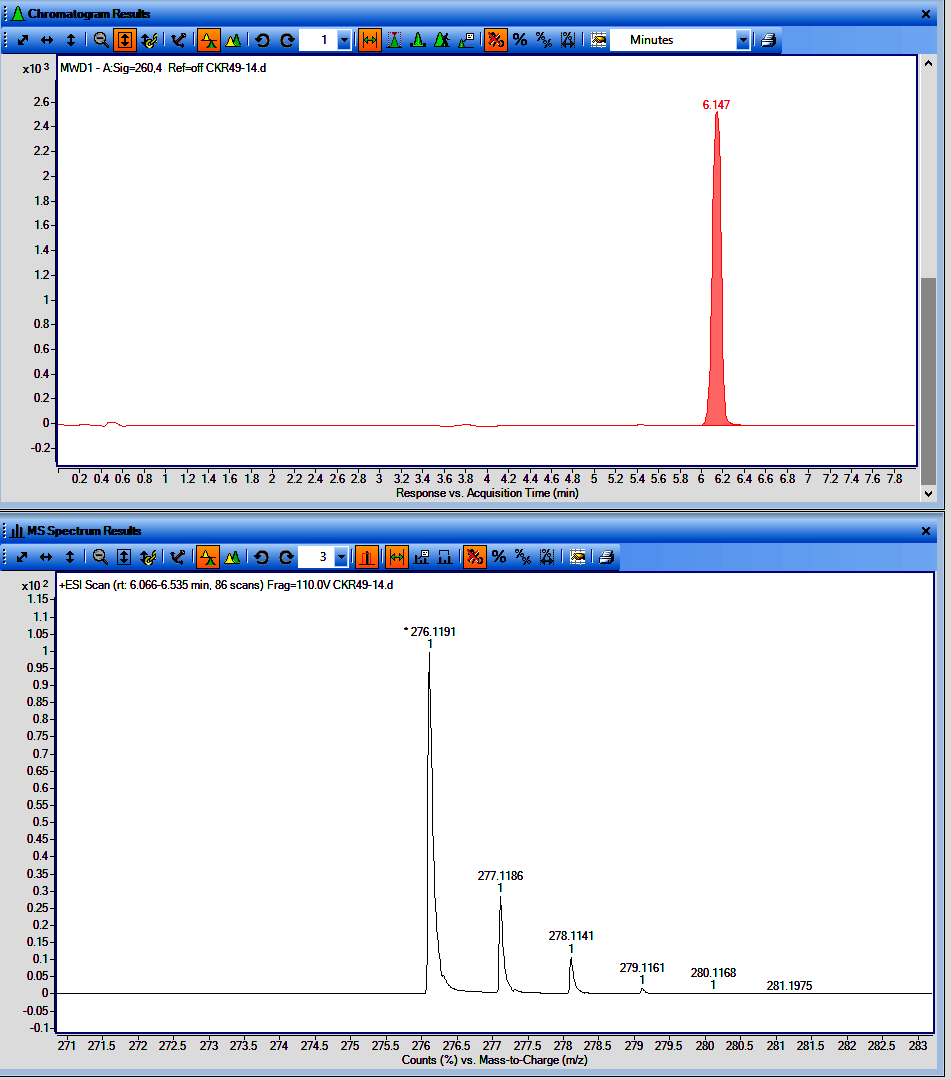

**Figure**. LCMS chromatogram of CKR49-14 (UV with t_R_ and MS spectrum)

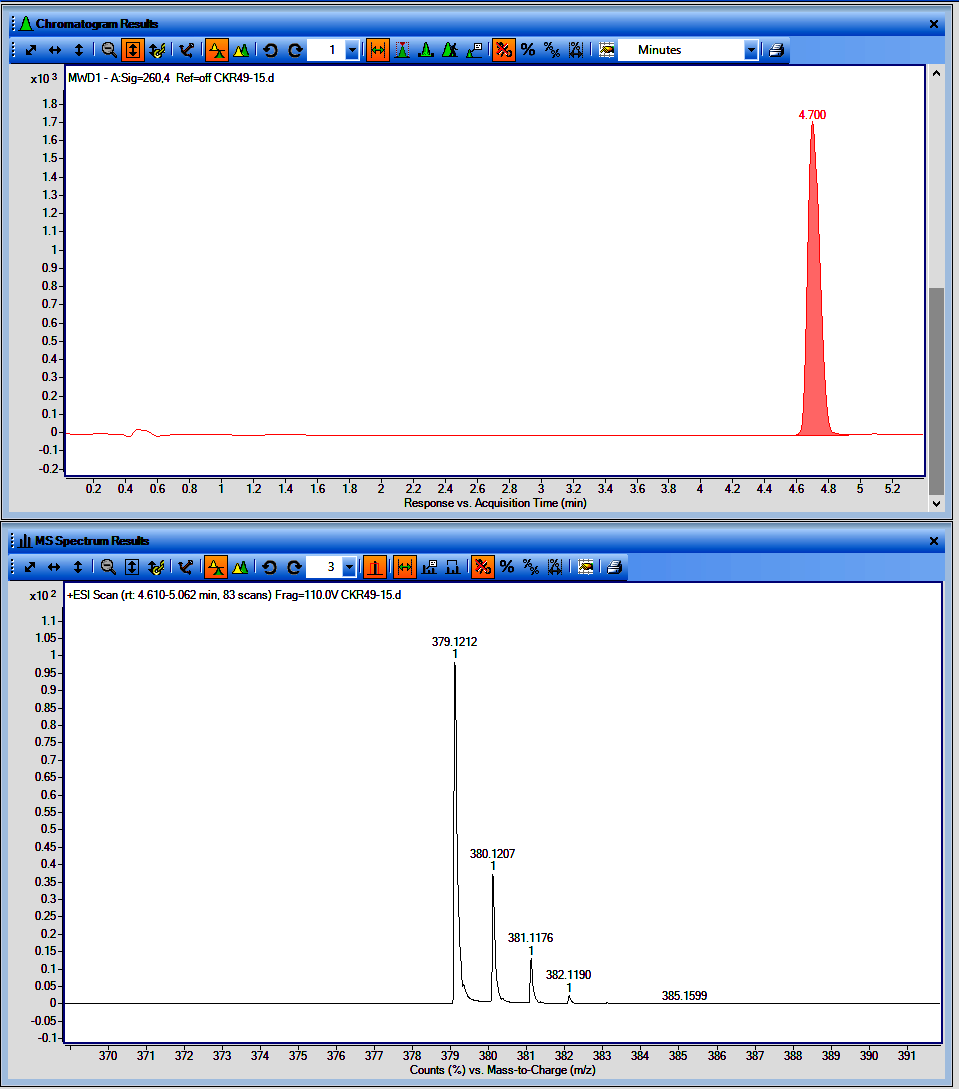

**Figure**. LCMS chromatogram of CKR49-15 (UV with t_R_ and MS spectrum)

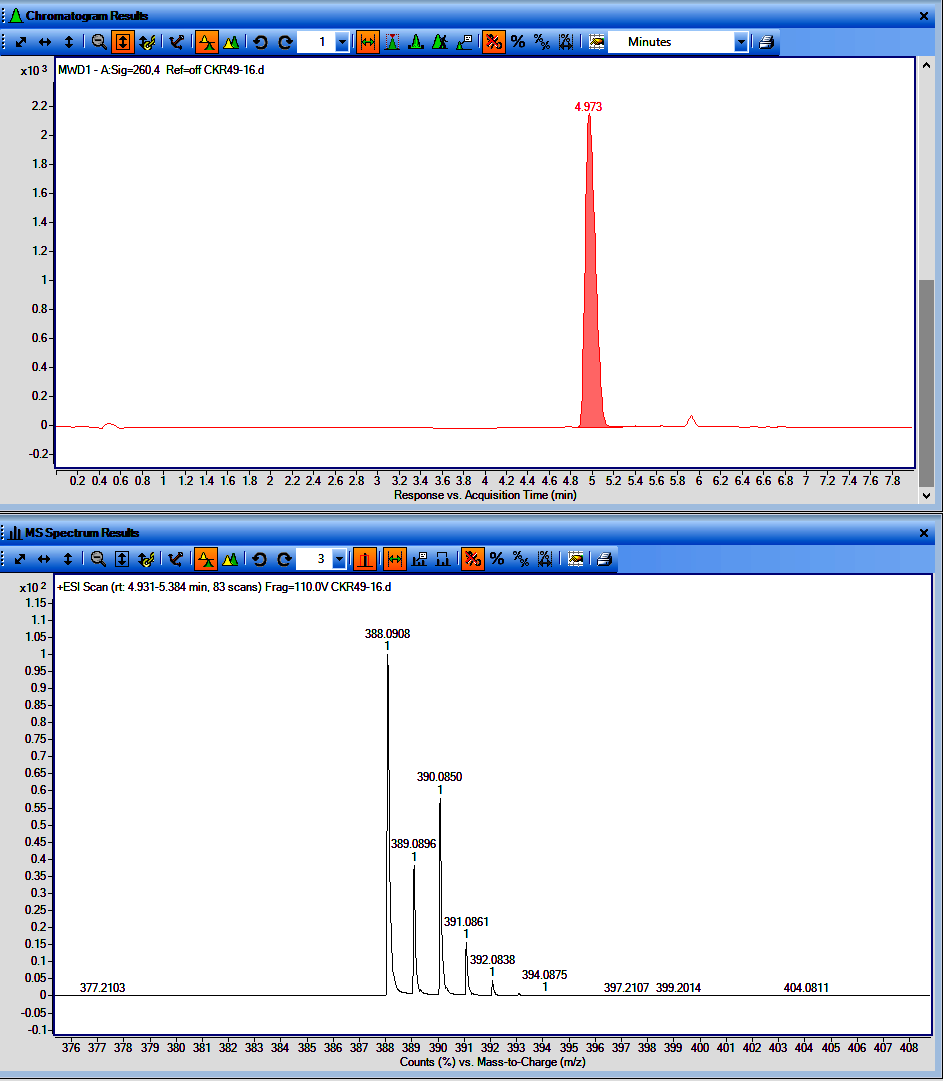

**Figure**. LCMS chromatogram of CKR49-16 (UV with t_R_ and MS spectrum)

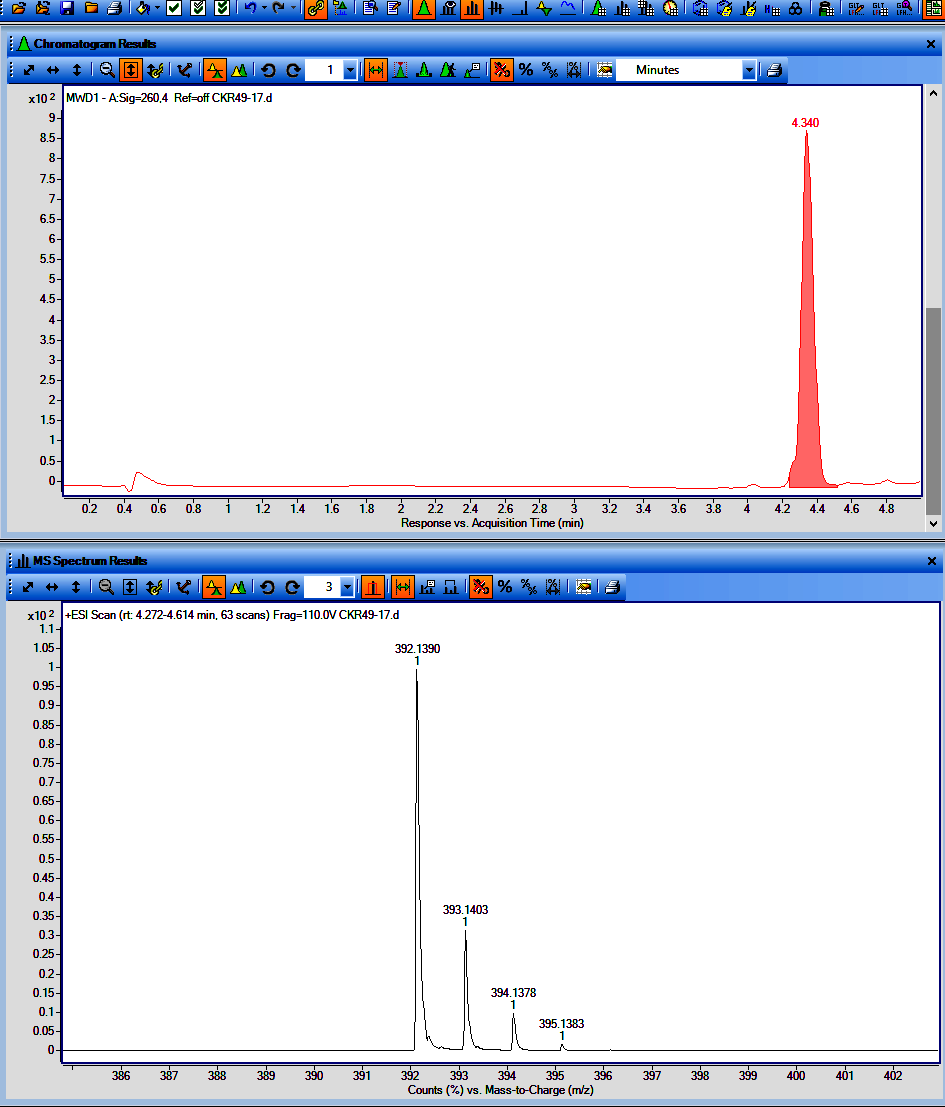

**Figure**. LCMS chromatogram of CKR49-17 (UV with t_R_ and MS spectrum)

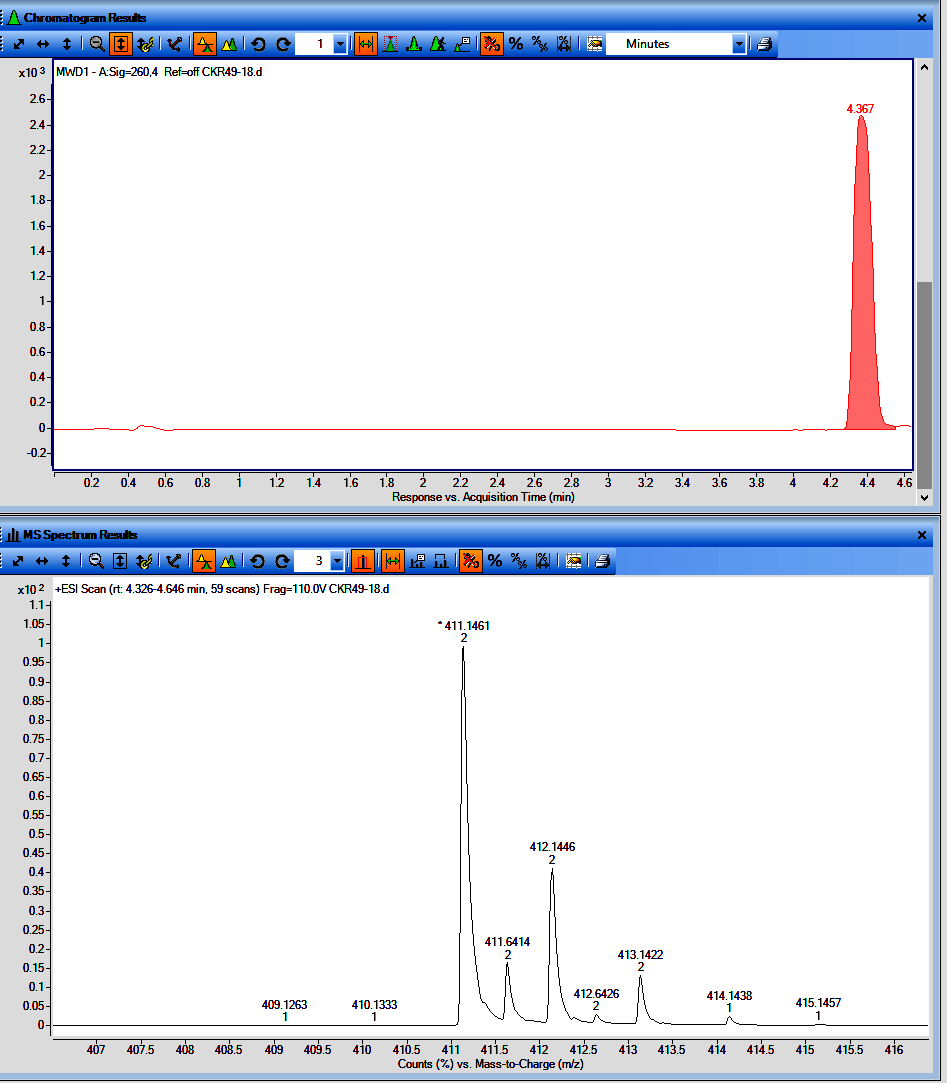

**Figure**. LCMS chromatogram of CKR49-18 (UV with t_R_ and MS spectrum)

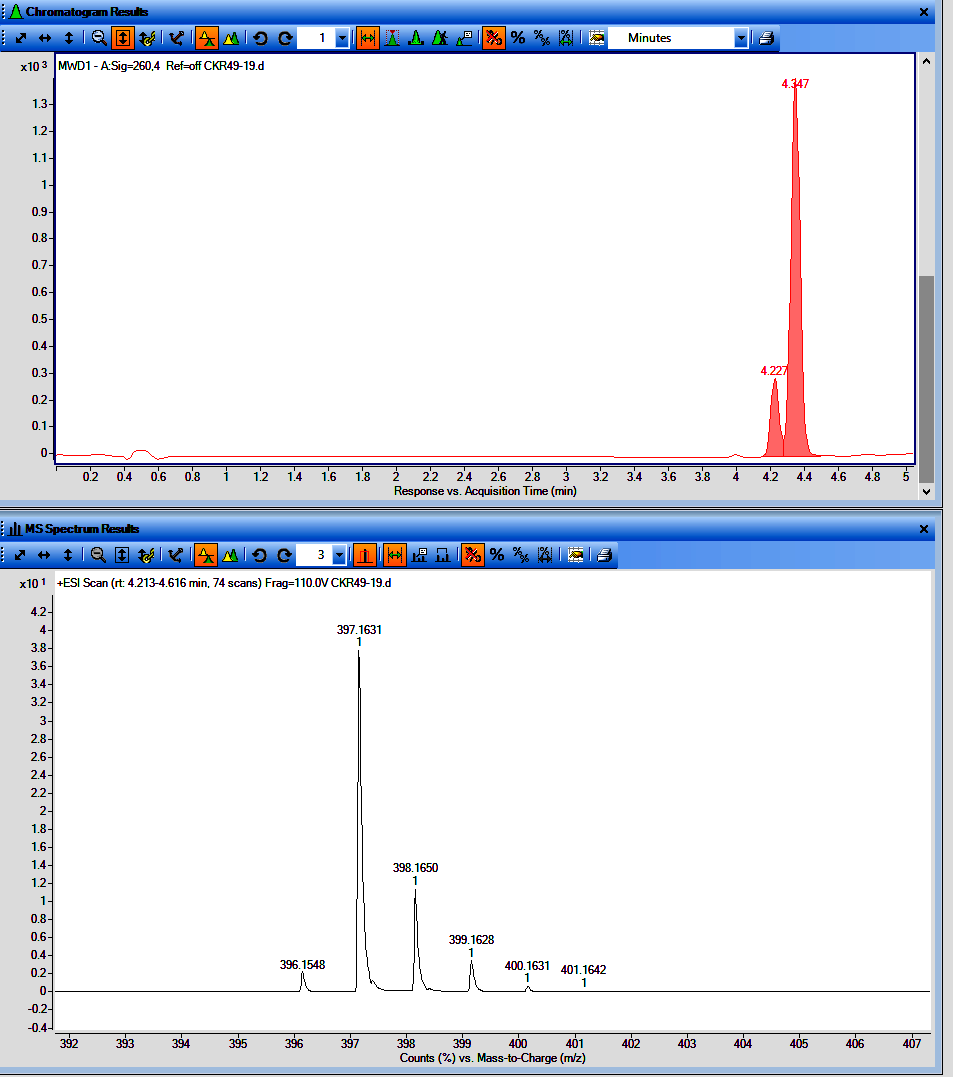

**Figure**. LCMS chromatogram of CKR49-19 (UV with t_R_ and MS spectrum)

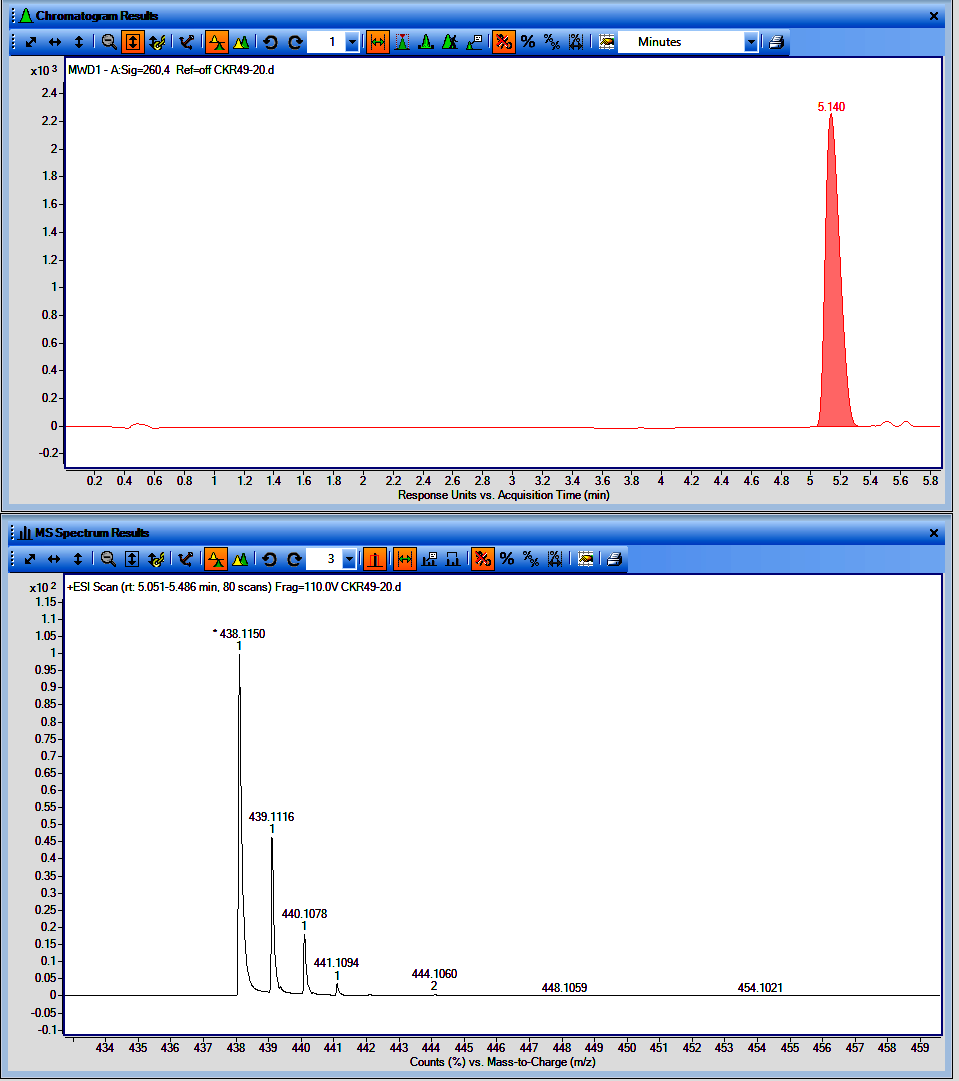

**Figure**. LCMS chromatogram of CKR49-20 (UV with t_R_ and MS spectrum)

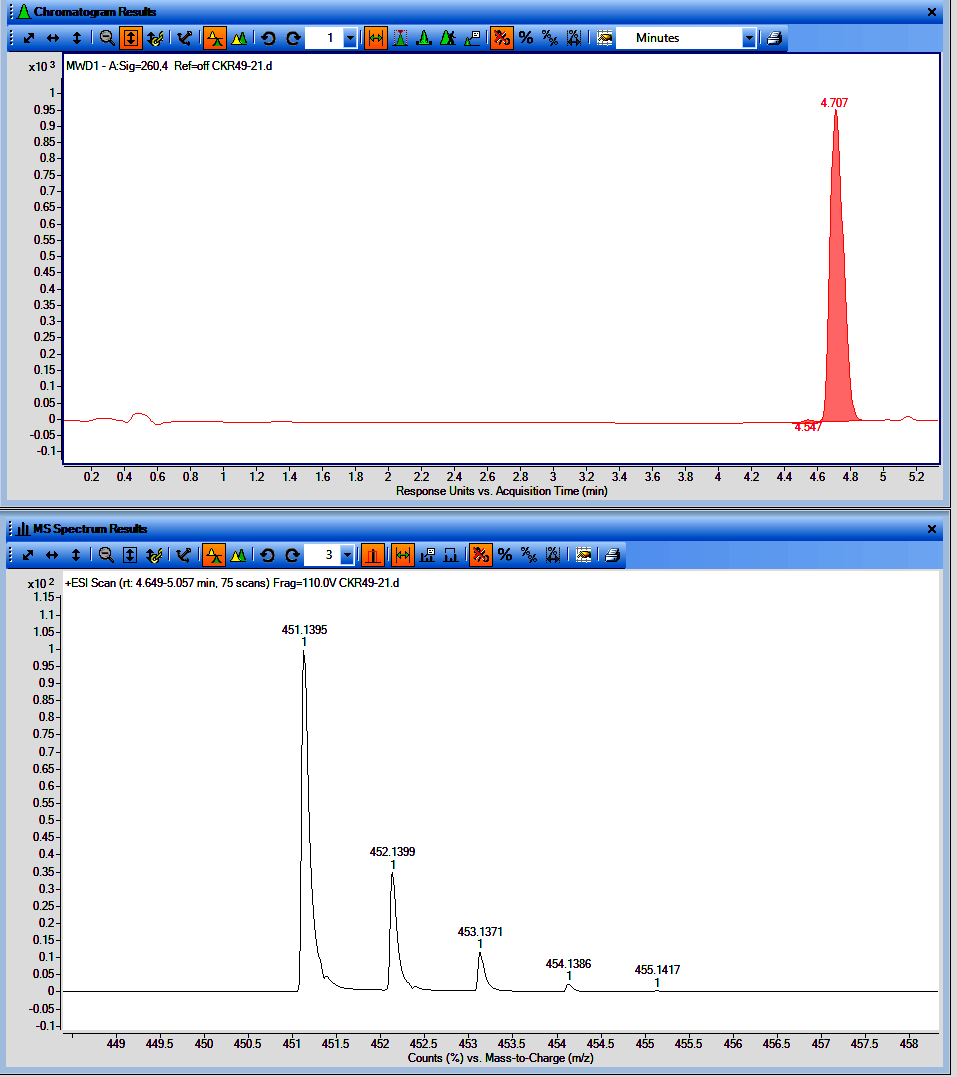

**Figure**. LCMS chromatogram of CKR49-21 (UV with t_R_ and MS spectrum)

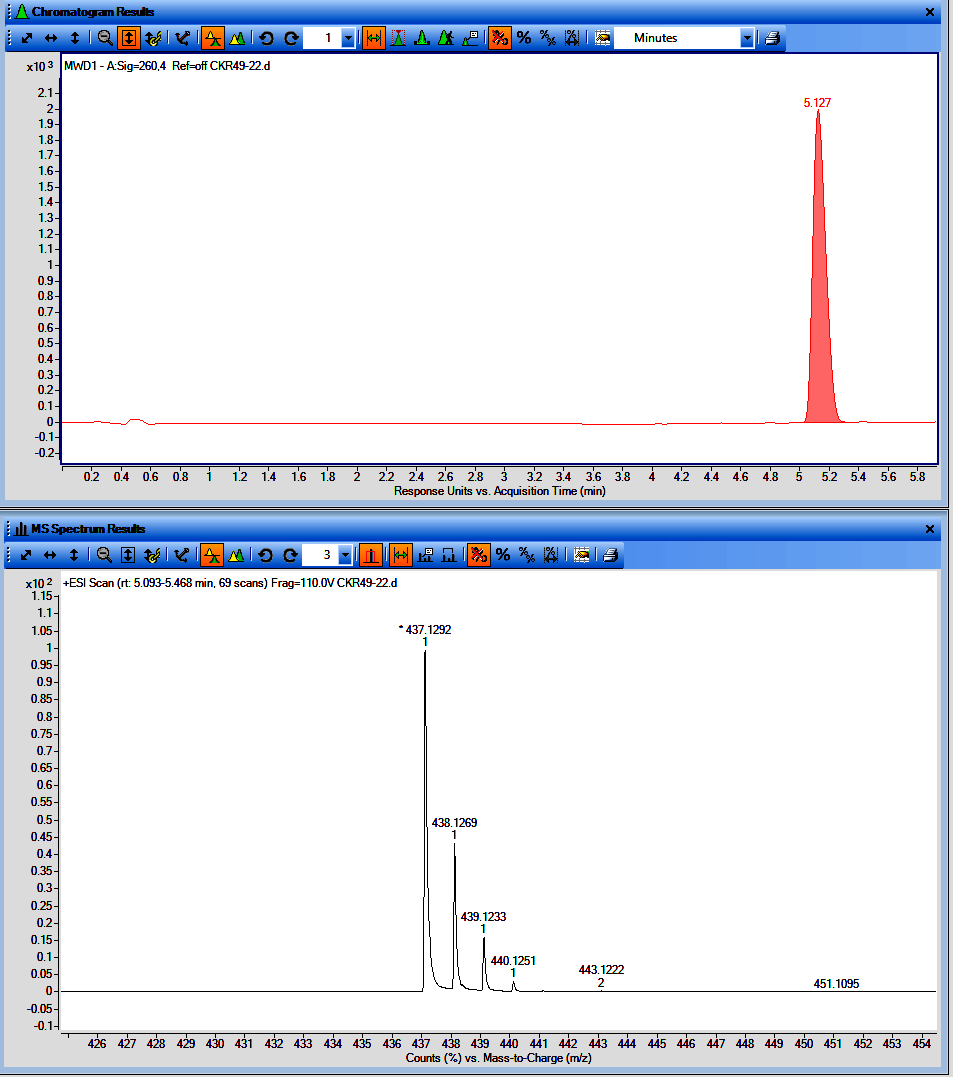

**Figure**. LCMS chromatogram of CKR49-22 (UV with t_R_ and MS spectrum)

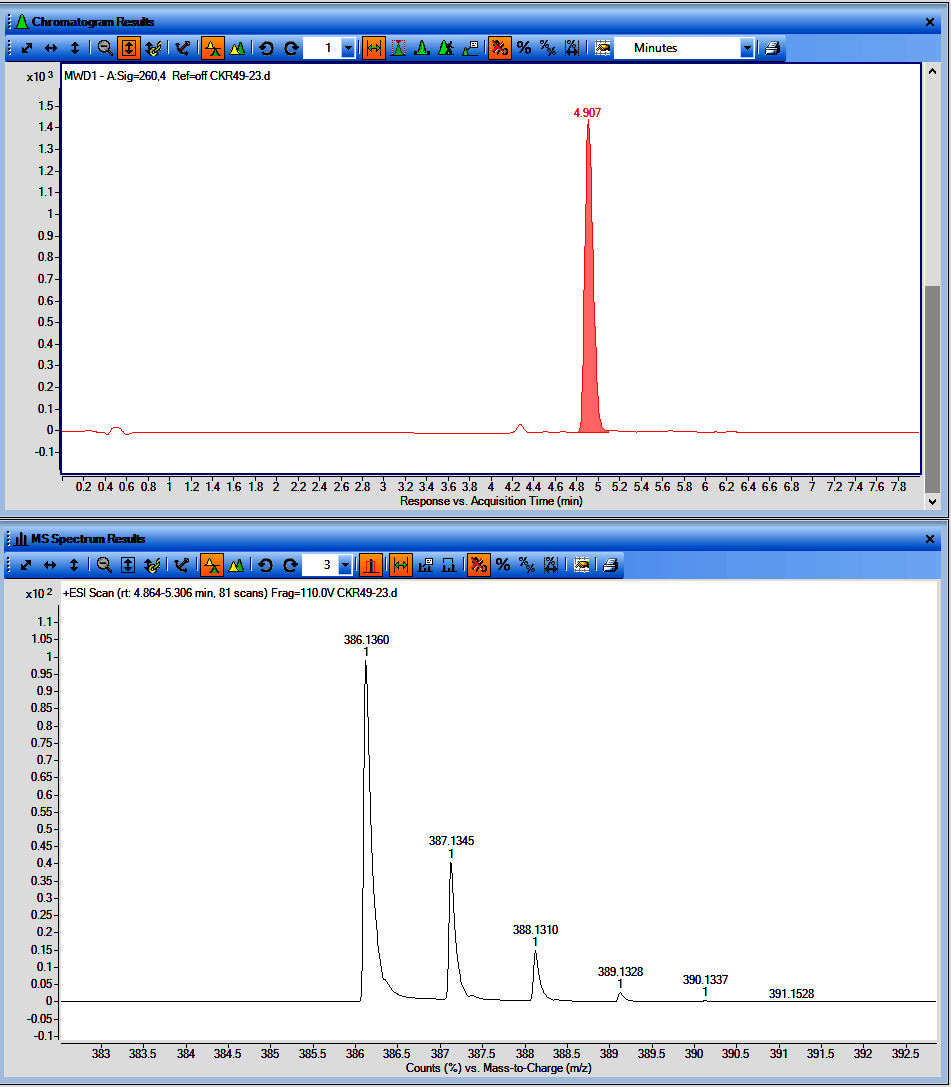

**Figure**. LCMS chromatogram of CKR49-23 (UV with t_R_ and MS spectrum)

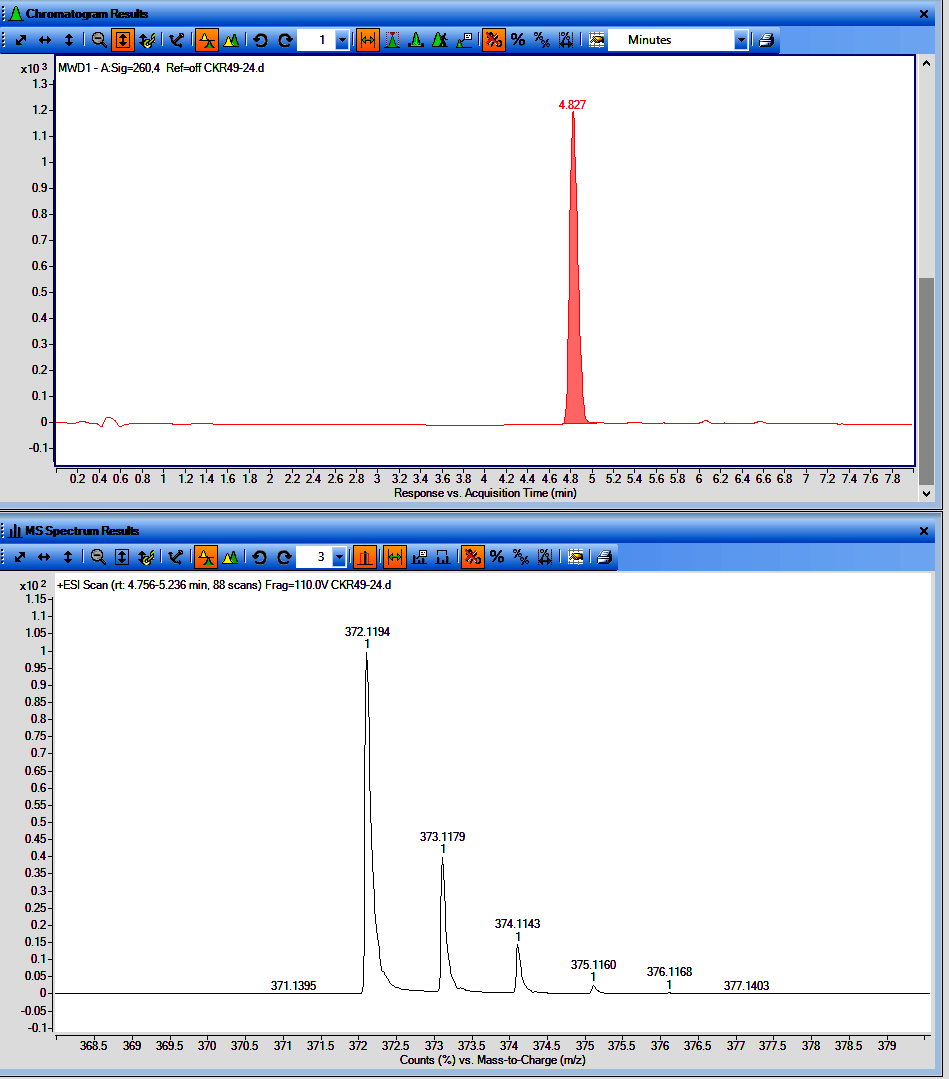

**Figure**. LCMS chromatogram of CKR49-24 (UV with t_R_ and MS spectrum)

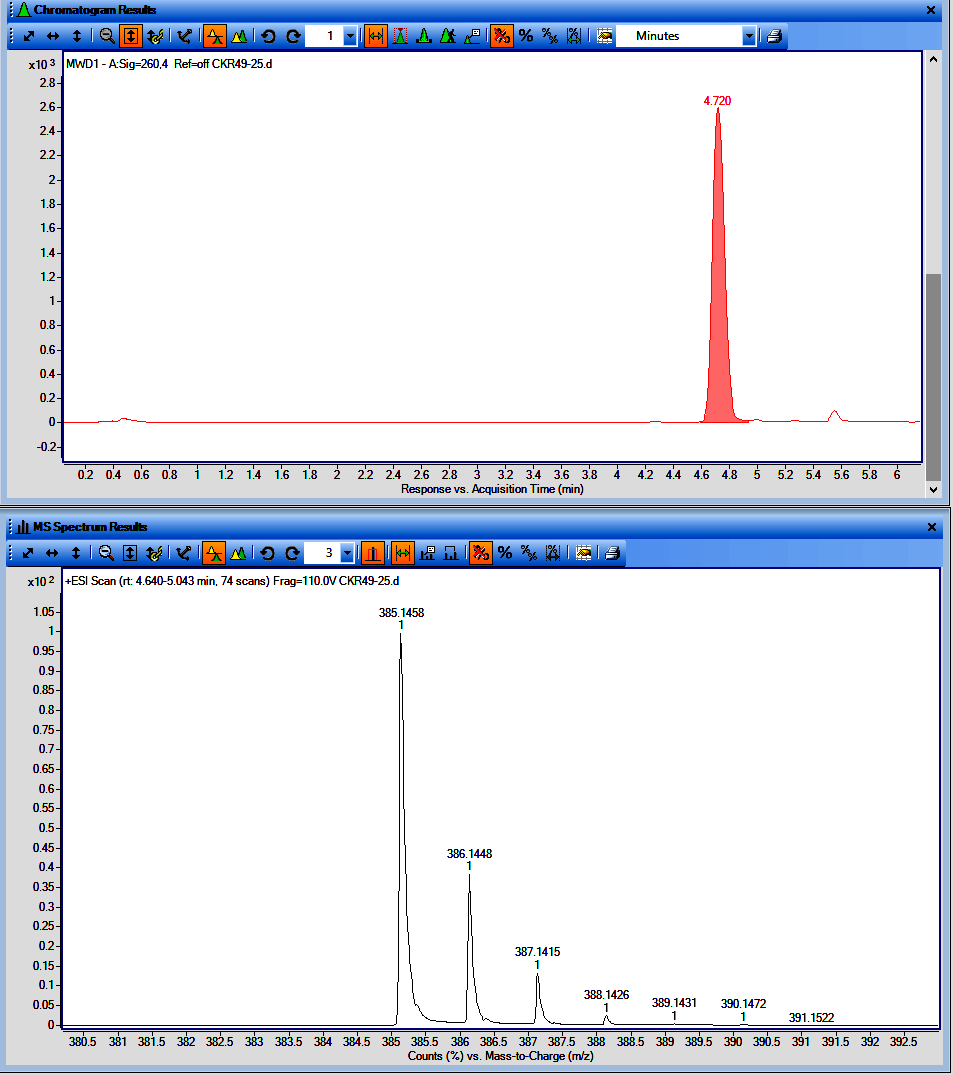

**Figure**. LCMS chromatogram of CKR49-25 (UV with t_R_ and MS spectrum)

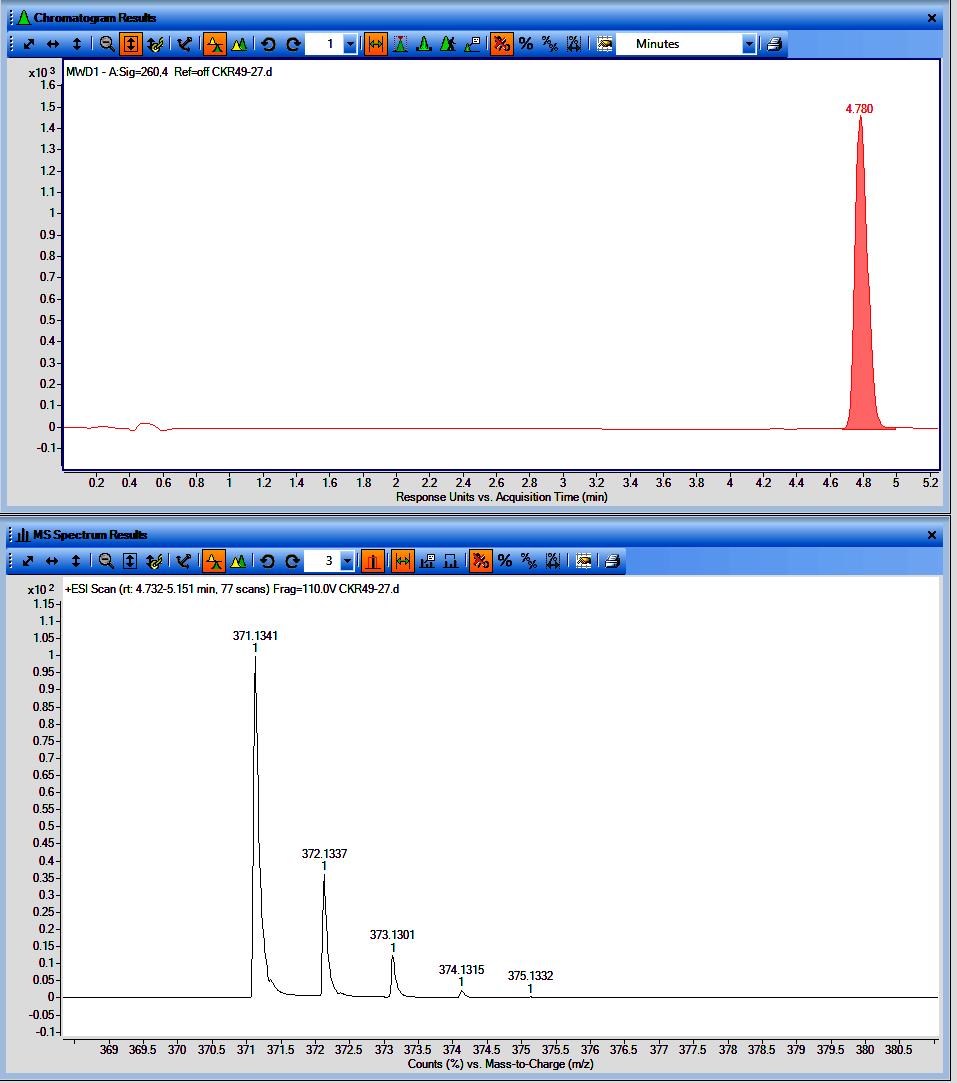

**Figure**. LCMS chromatogram of CKR49-27 (UV with t_R_ and MS spectrum)

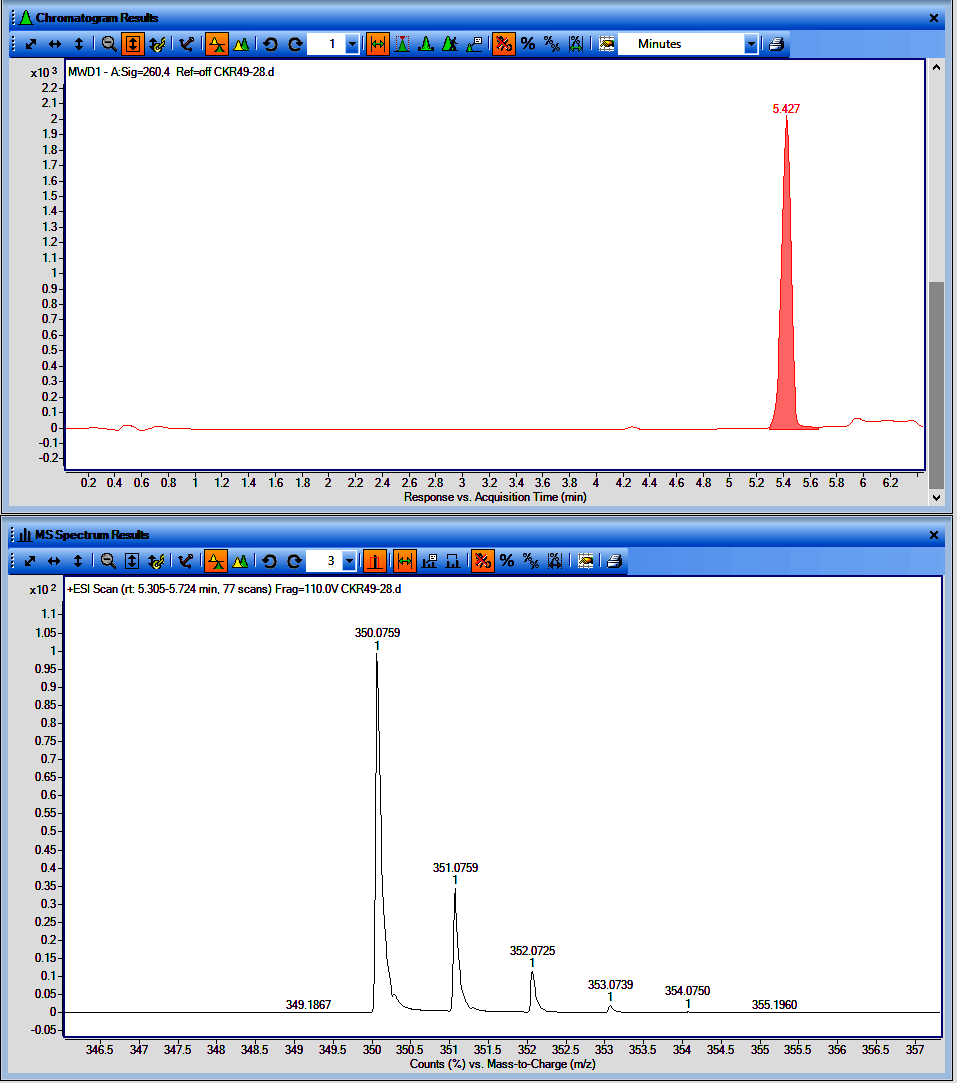

**Figure**. LCMS chromatogram of CKR49-28 (UV with t_R_ and MS spectrum)

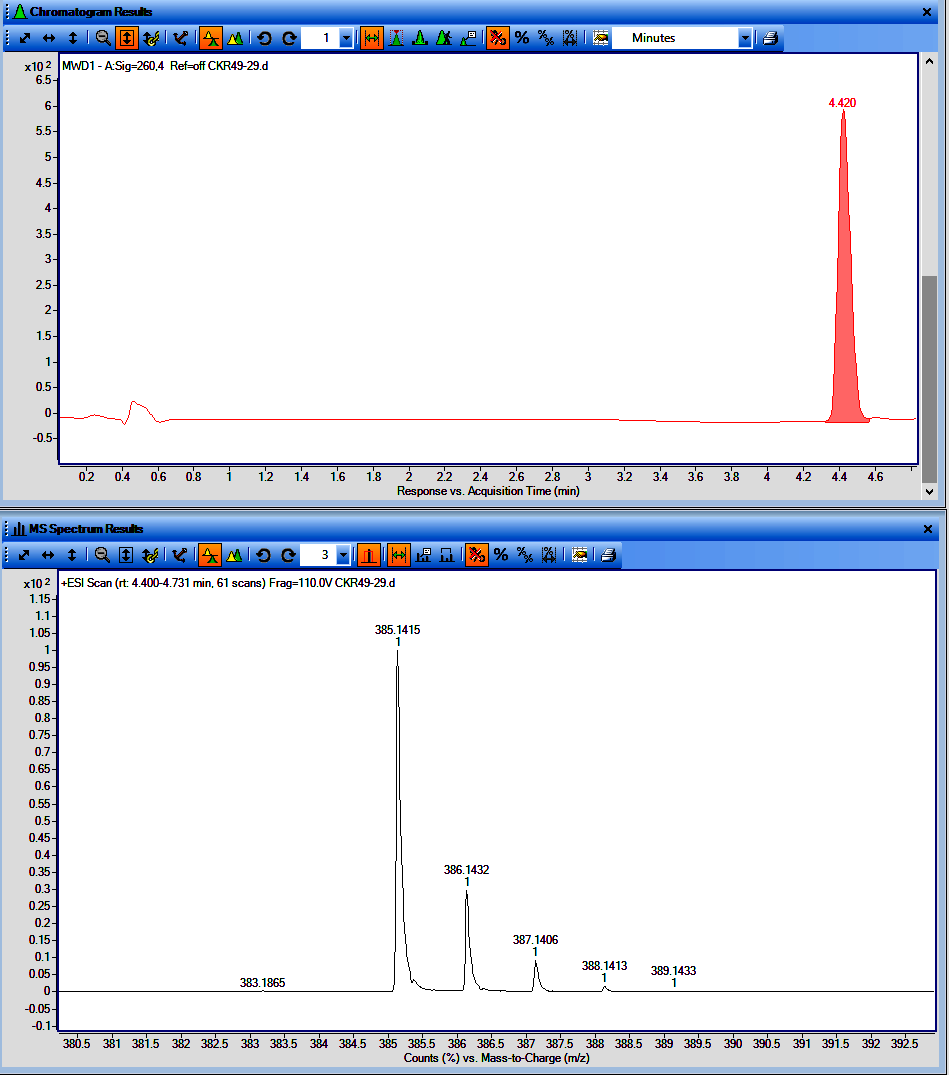

**Figure**. LCMS chromatogram of CKR49-29 (UV with t_R_ and MS spectrum)

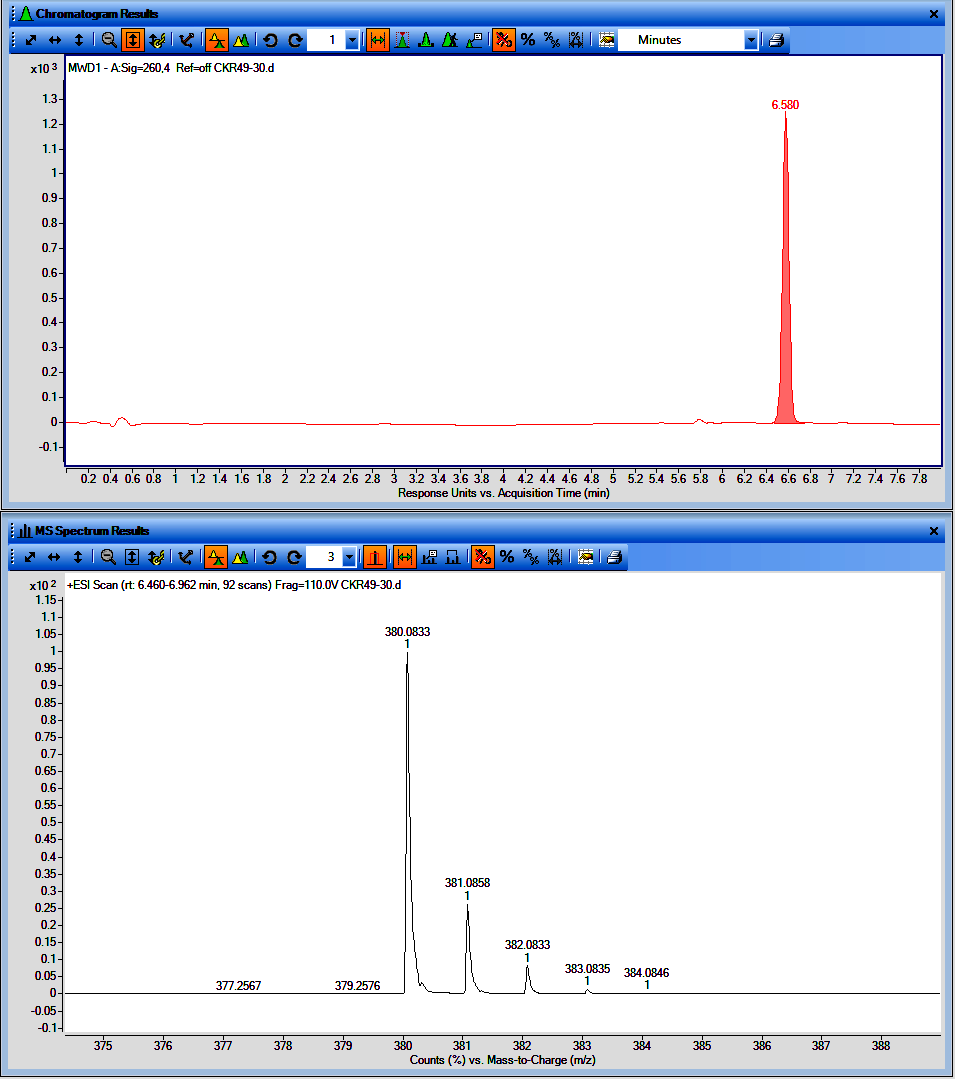

**Figure**. LCMS chromatogram of CKR49-30 (UV with t_R_ and MS spectrum)

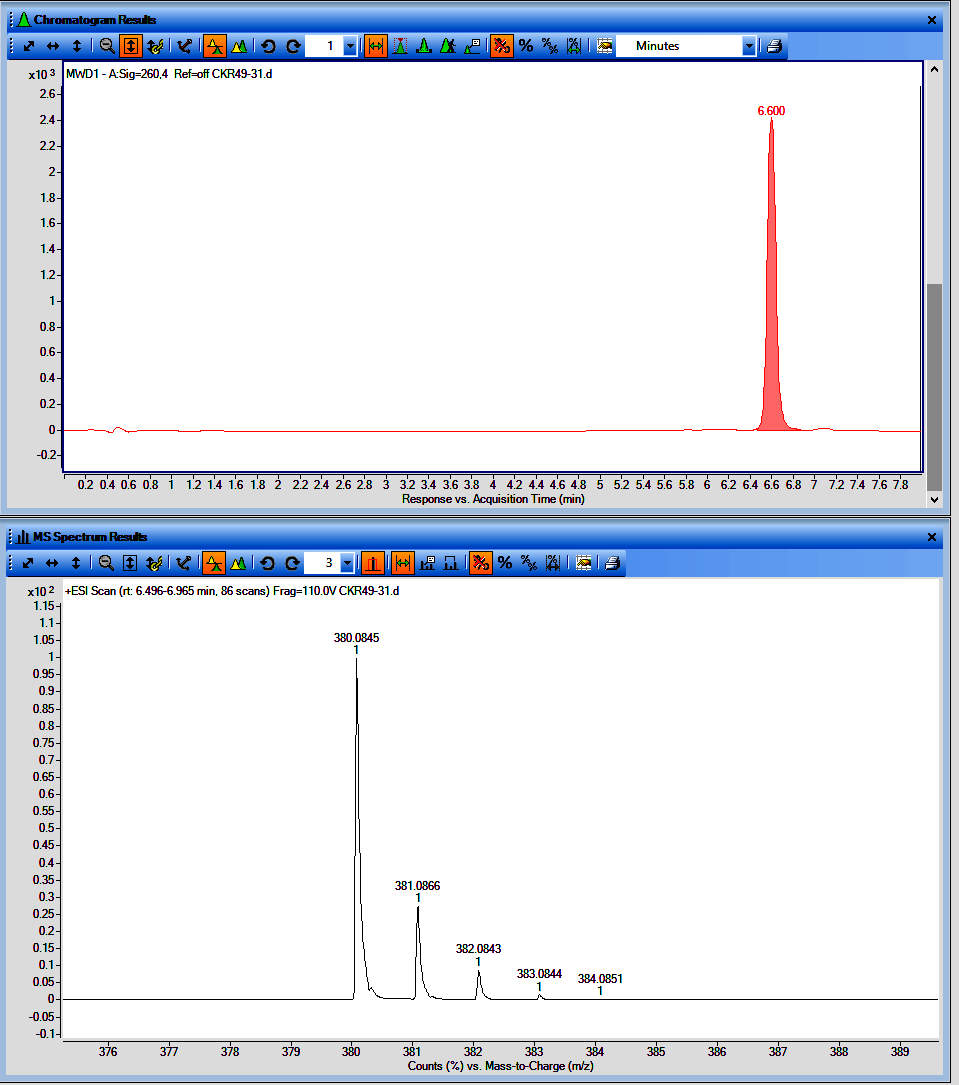

**Figure**. LCMS chromatogram of CKR49-31 (UV with t_R_ and MS spectrum)

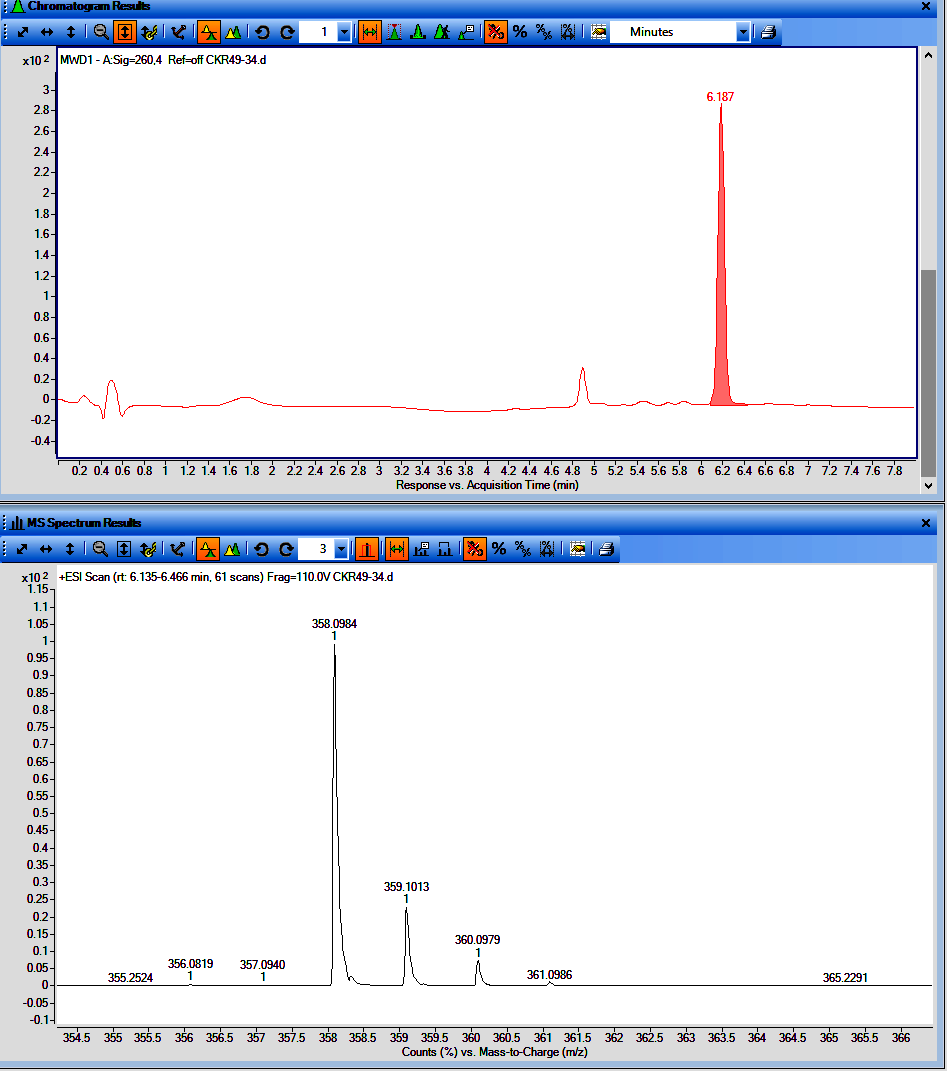

**Figure**. LCMS chromatogram of CKR49-34 (UV with t_R_ and MS spectrum)

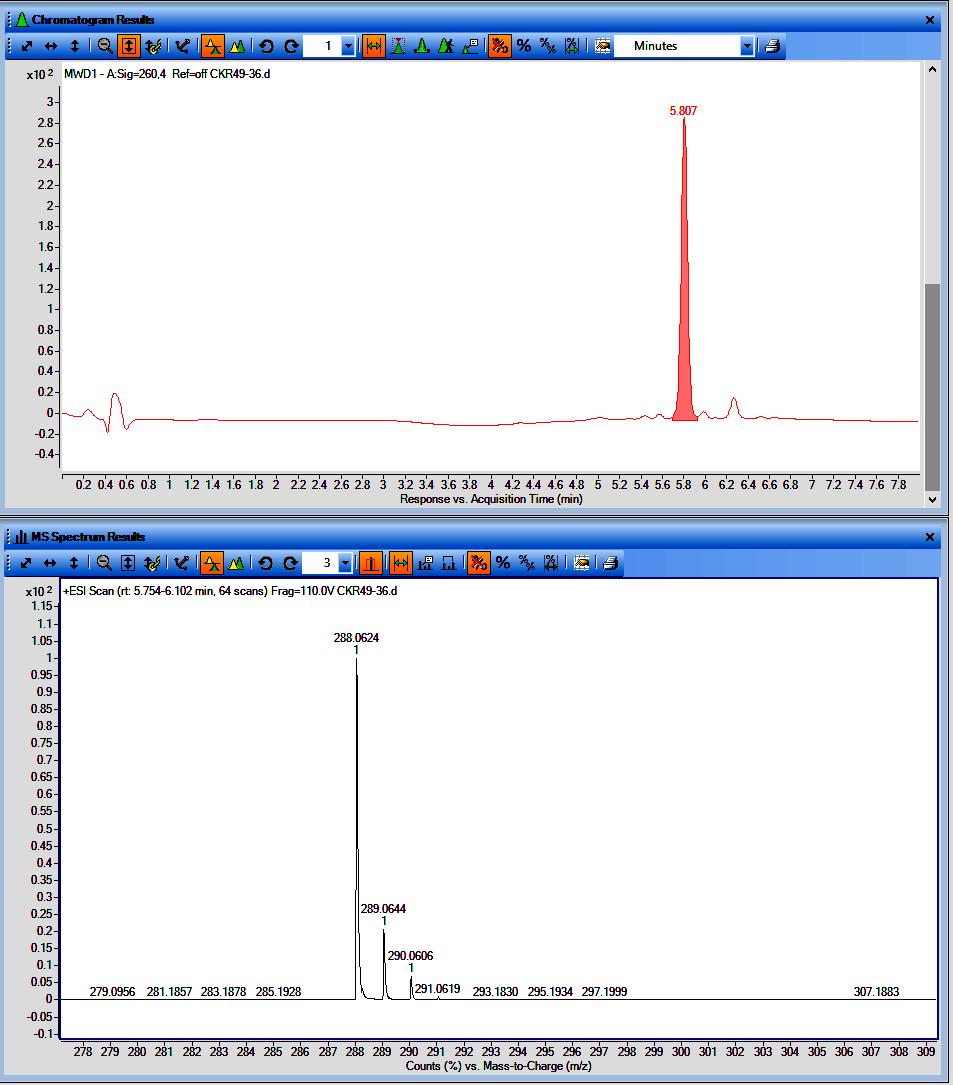

**Figure**. LCMS chromatogram of CKR49-36 (UV with t_R_ and MS spectrum)

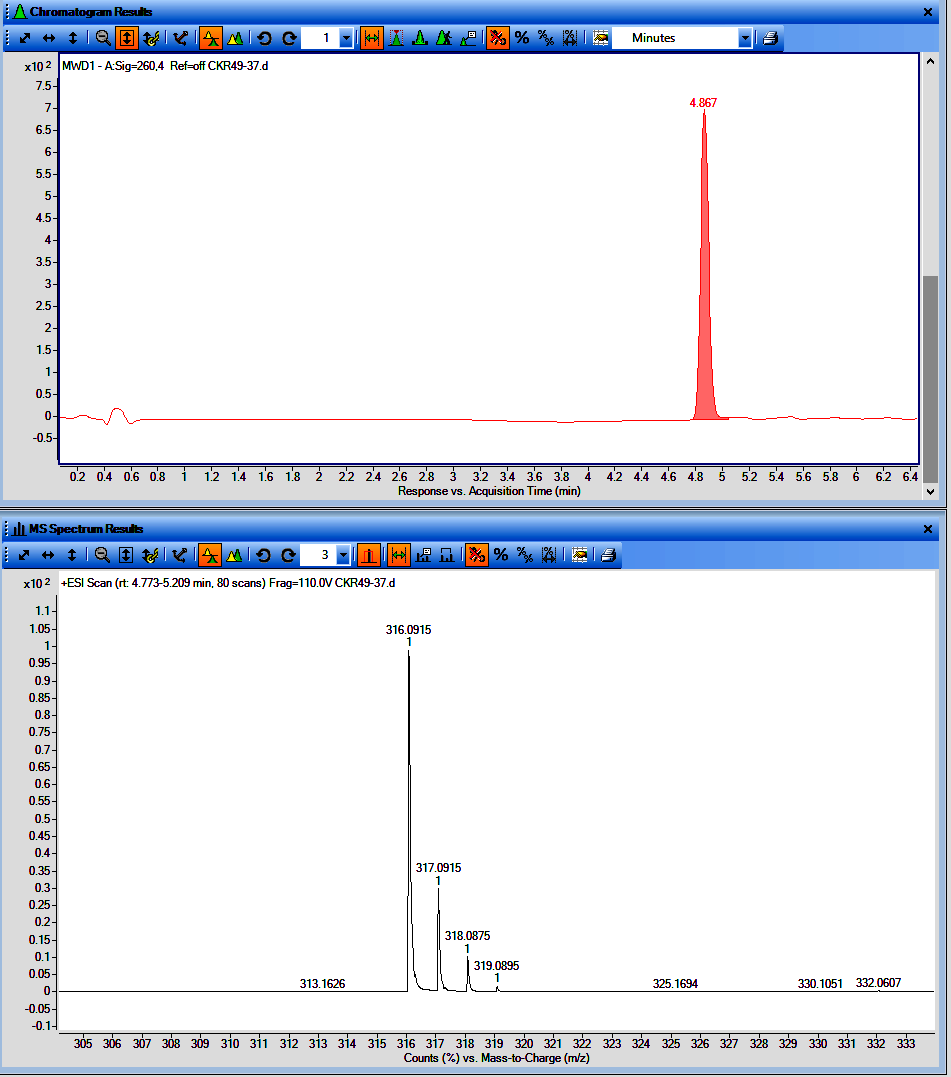

**Figure**. LCMS chromatogram of CKR49-37 (UV with t_R_ and MS spectrum)

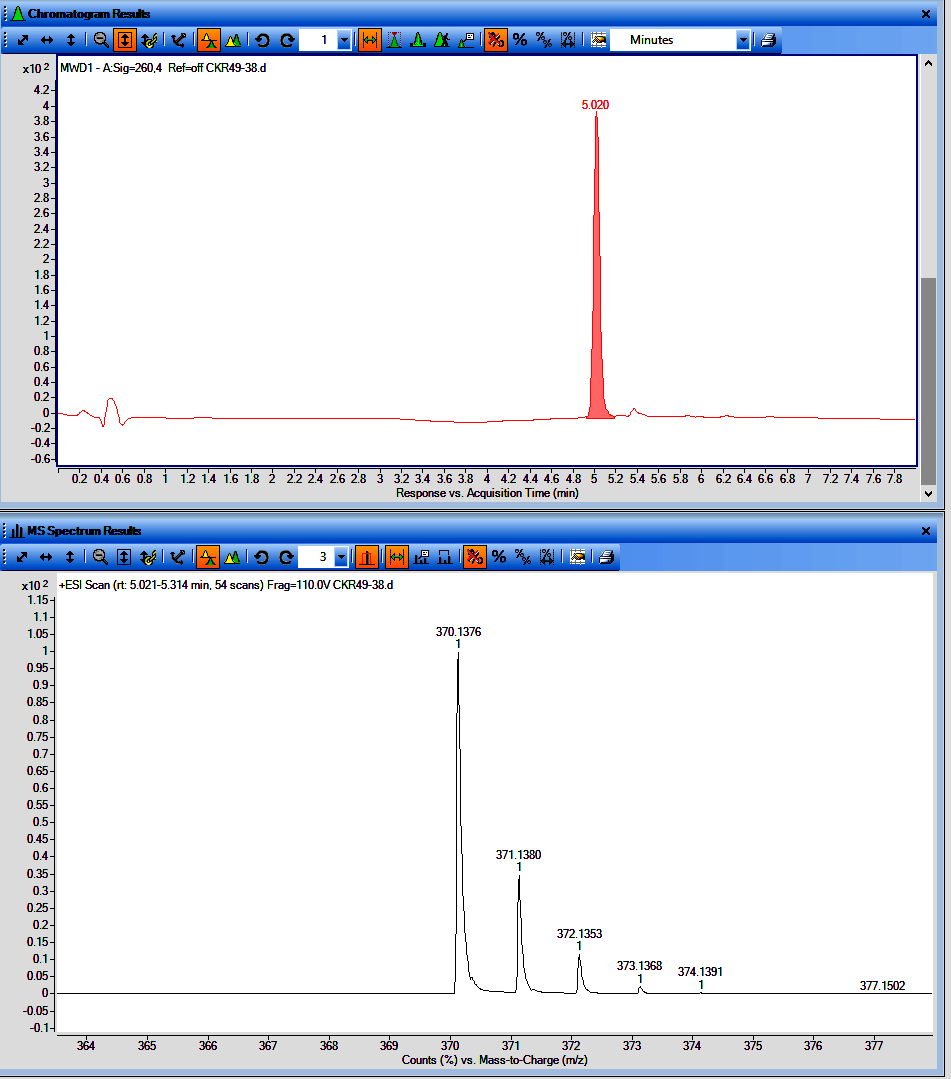

**Figure**. LCMS chromatogram of CKR49-38 (UV with t_R_ and MS spectrum)

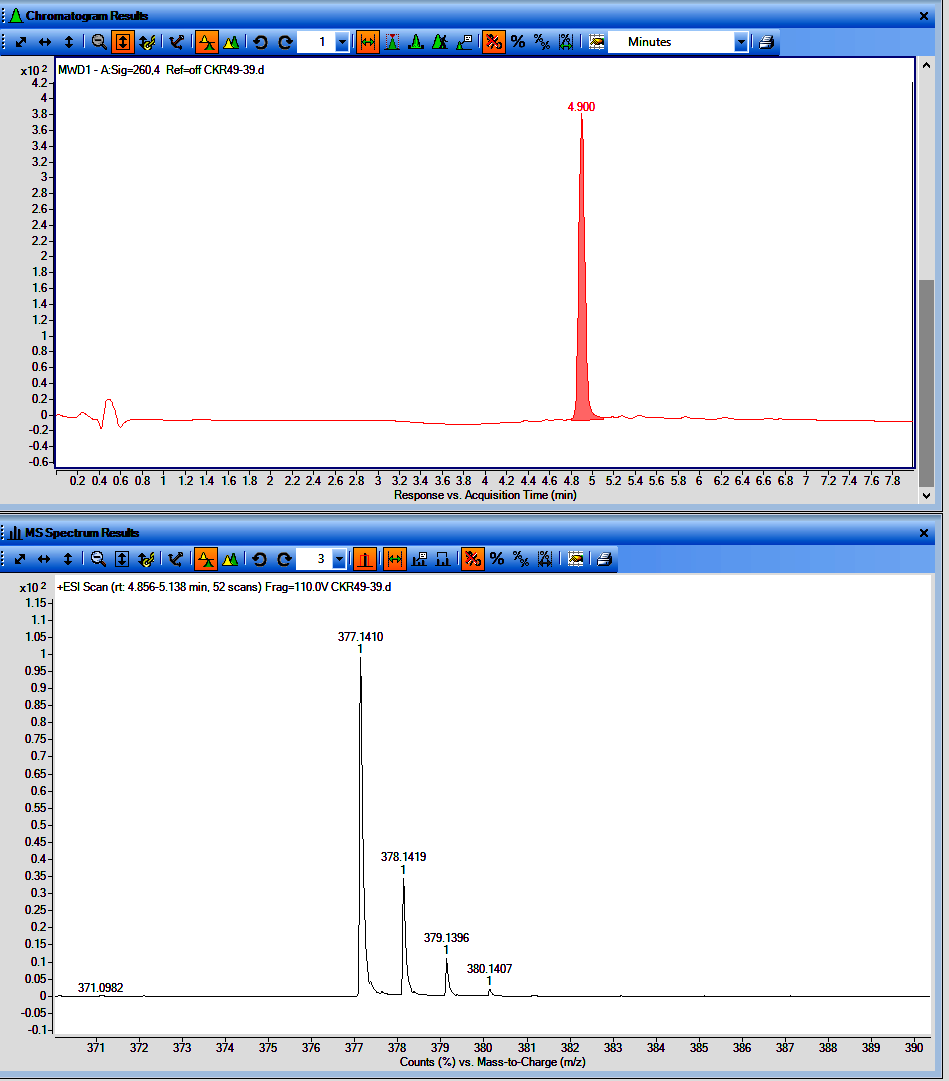

**Figure**. LCMS chromatogram of CKR49-39 (UV with t_R_ and MS spectrum)

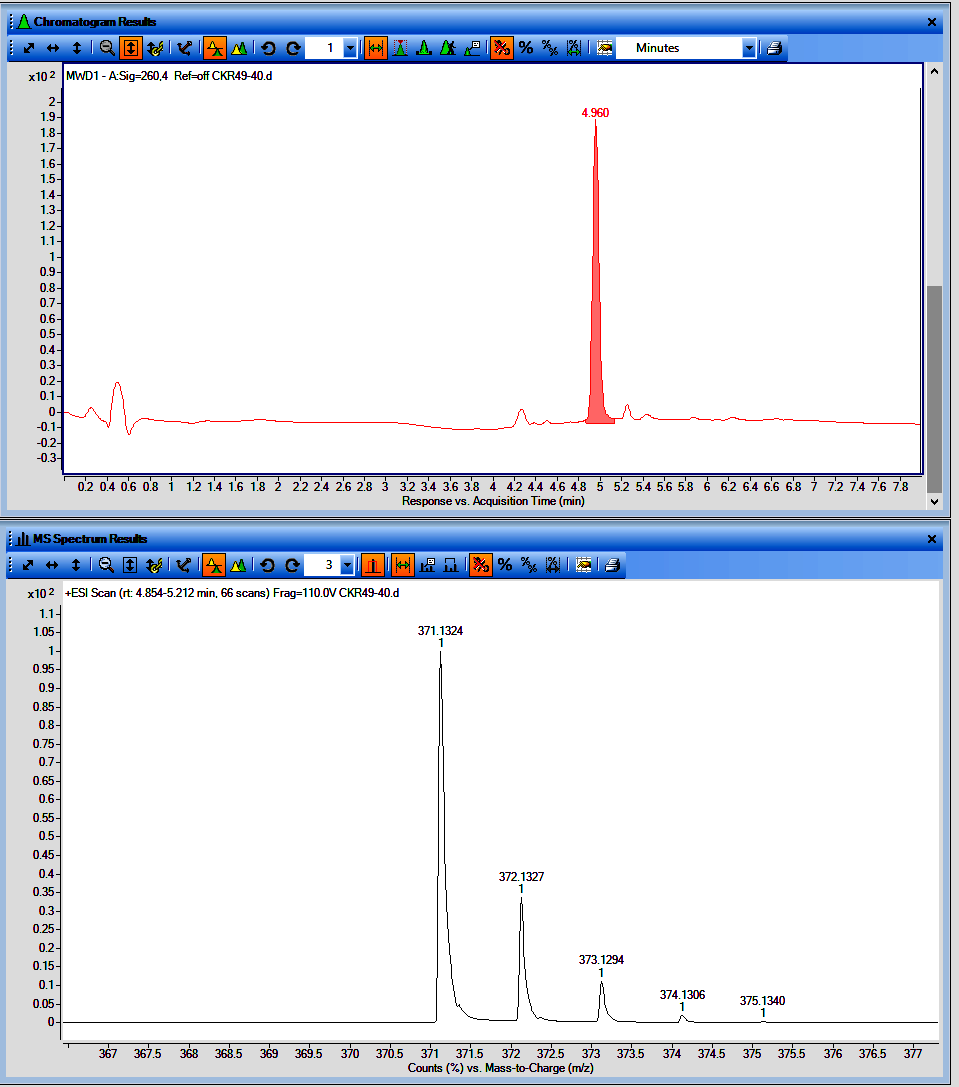

**Figure**. LCMS chromatogram of CKR49-40 (UV with t_R_ and MS spectrum)

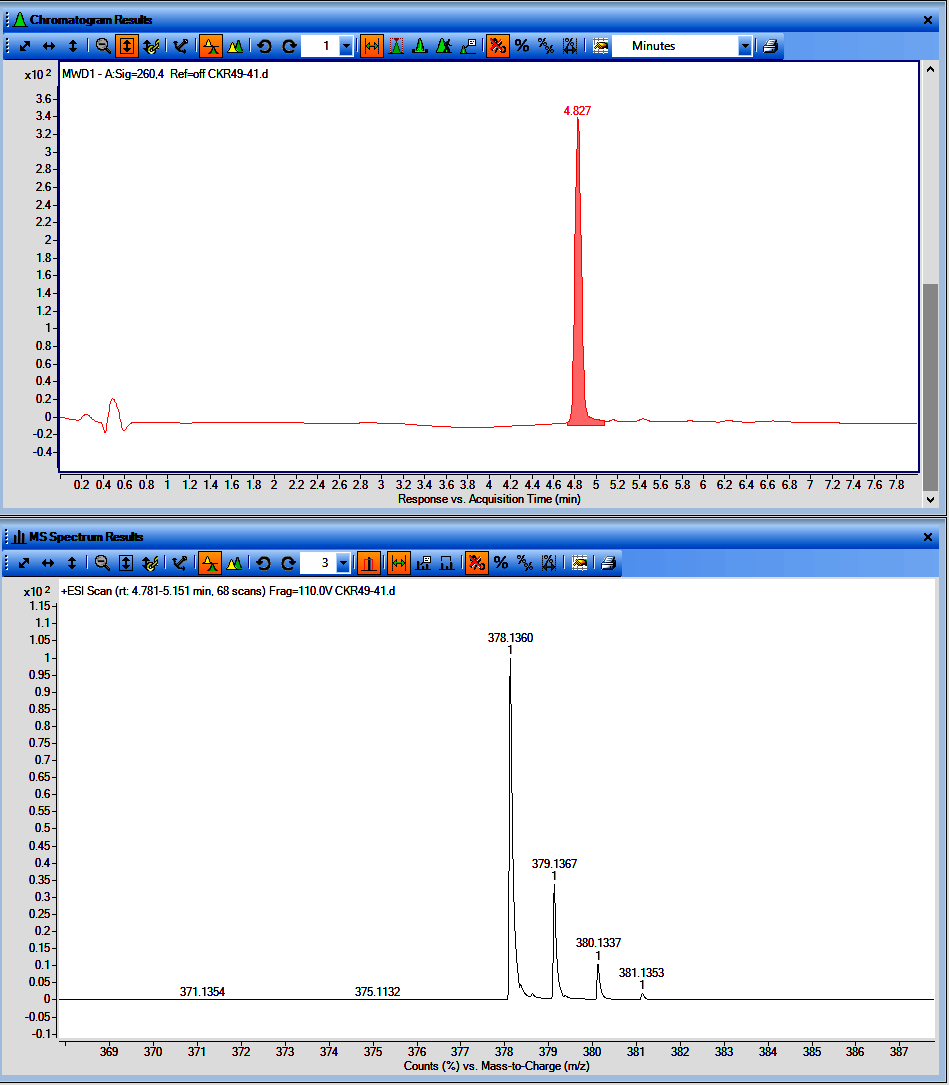

**Figure**. LCMS chromatogram of CKR49-41 (UV with t_R_ and MS spectrum)

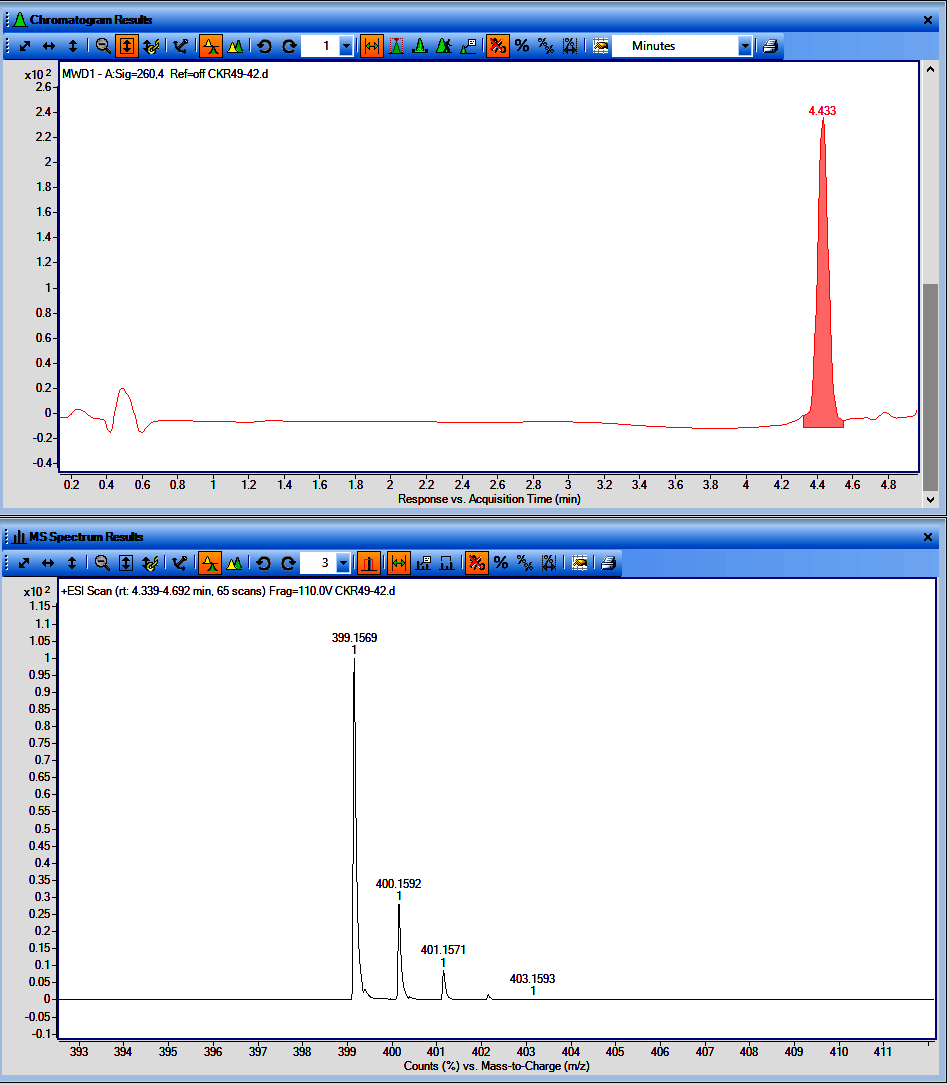

**Figure**. LCMS chromatogram of CKR49-42 (UV with t_R_ and MS spectrum)
